# Discovery of the sEH Inhibitor Epoxykynin as Potent Kynurenine Pathway Modulator

**DOI:** 10.1101/2023.11.09.566383

**Authors:** Lara Dötsch, Caitlin Davies, Elisabeth Hennes, Julia Schönfeld, Adarsh Kumar, Celine Da Cruz Lopes Guita, Johanna H.M. Ehrler, Kerstin Hiesinger, Sasikala Thavam, Petra Janning, Sonja Sievers, Stefan Knapp, Ewgenij Proschak, Slava Ziegler, Herbert Waldmann

**Affiliations:** Max Planck Institute of Molecular Physiology, Department of Chemical Biology, Otto-Hahn-Strasse 11, 44227 Dortmund (Germany); Technical University of Dortmund, Department of Chemical Biology, Otto-Hahn-Strasse 6, 44227 Dortmund (Germany); Vertex Pharmaceuticals, 86-88 Jubilee Ave, Milton, Abingdon OX14 4RW (United Kingdom); Compound Management and Screening Center (COMAS), Otto-Hahn-Strasse 15, 44227 Dortmund (Germany); Goethe University Frankfurt, Institute of Pharmaceutical Chemistry, Max-von-Laue-Strasse 9, 60438 Frankfurt (Germany); Structural Genomics Consortium, Buchmann Institute for Molecular Life Sciences, Goethe University Frankfurt, Max-von-Laue-Strasse 15, 60438 Frankfurt (Germany)

## Abstract

Disease-related phenotypic assays enable unbiased discovery of novel bioactive small molecules and may provide novel insights into physiological systems and unprecedented molecular modes-of-action (MMOA). Herein we report the identification and characterization of epoxykynin, a potent inhibitor of the soluble epoxide hydrolase (sEH). Epoxykynin was discovered by means of a cellular assay monitoring modulation of kynurenine (Kyn) levels in BxPC-3 cells upon stimulation with the cytokine interferon-γ (IFN-γ) and subsequent target identification employing affinity-based chemical proteomics. Increased Kyn levels are associated with immune suppression in the tumor microenvironment and, thus, the Kyn pathway and its key player indoleamine 2,3-dioxygenase 1 (IDO1) are appealing targets in immuno-oncology. However, targeting IDO1 directly has led to limited success in clinical investigations, demonstrating that alternative approaches to reduce Kyn levels are in high demand. We uncover a cross-talk between sEH and the Kyn pathway that may provide new opportunities to revert cancer-induced immune tolerance.

## Introduction

Phenotypic screening allows for the identification of small molecules that modulate specific cellular processes in their natural environment, i.e. in the cell. This may lead to the discovery of novel bioactive compounds, as well as better understanding of biological pathways. Compared to target-based approaches, phenotypic assays enable a less biased detection of biologically active compounds.^1^ The design of physiologically relevant screening assays should consider utilizing a disease-relevant system, employing a physiological stimulus and an appropriate downstream readout (phenotypic screening “rule of 3”).^2^

Indoleamine 2,3-dioxygenase 1 (IDO1) is a heme-containing enzyme that catalyzes the conversion of L-tryptophan (Trp) into the metabolite kynurenine (Kyn).^3^ IDO1 plays a critical role in the suppression of the immune system, particularly in the context of cancer.^4-6^ The reduction of Trp in the tumor microenvironment and the simultaneous production of Kyn leads to T cell dysfunction and immune tolerance.^7^ Targeting IDO1 directly has shown promising results in the treatment of cancer in preclinical studies and more than 50 different clinical trials with the most advanced IDO1 inhibitor epacadostat have been launched.^8^ However, the recent failure of a large phase III trial, testing epacadostat in combination with the anti-PD1 antibody pembrolizumab (ECHO-301/KN-252),^9^ has put many trials on hold. Reasons for failure include that the patients for the study were not specifically selected for IDO1 expression in the tumor, compensatory expression of the two other Trp-catabolizing dioxygenases tryptophan 2,3-dioxygenase (TDO) and indoleamine 2,3-dioxygenase 2 (IDO2) or activation of the aryl hydrocarbon receptor (AhR) by epacadostat, leading to immune tolerance.^10-12^ The negative outcome of ECHO-301 showed that consideration of the Kyn pathway as a whole is crucial for its relevance in immuno-oncology. Therefore, alternative approaches for reduction of Kyn levels and new targets are in high demand.

We recently reported on the development of a cell-based assay monitoring Kyn levels after stimulation of BxPC-3 cells with the cytokine interferon-γ (IFN-γ) to induce expression of IDO1.^13^ Using this assay, we have now identified *N*-substituted indoles as compound class that potently reduce cellular Kyn levels upon stimulation of cancer cells with IFN-γ.^13,14^ These small molecules do not inhibit IDO1 activity or expression, but modulate the Kyn pathway by targeting the soluble epoxide hydrolase (sEH). The most potent derivative, termed epoxykynin, inhibits the hydrolase activity of sEH (sEH-H) but not its phosphatase activity (sEH-P). Our results demonstrate a previously unrecognized dependence of the Kyn pathway on sEH function. These findings open up new avenues for the application of sEH inhibitors to modulate disease states linked to increased Kyn production.

## Results and Discussion

### A cell-based screening identifies N-substituted indoles as inhibitors of the Kyn pathway

We employed a cell-based assay and screened 157,332 commercial and in-house synthesized compounds to identify modulators of Kyn levels in BxPC-3 cells upon stimulation with IFN-γ.^13^ In contrast to IDO1, the expression of the two other Trp-catabolizing enzymes TDO and IDO2 cannot be induced by cytokines.^15-17^ Thereby, compounds identified by this screening method most likely interfere with IDO1-mediated Kyn production.

Among the identified hits, 1,3,5-substituted indole **1a** stood out and potently reduced cellular Kyn levels by 95.5 ± 3.0% at 7.1 μM with an IC50 value of 90 ± 26 nM (Table 1, Table S1, Figure 1). Exploration of the structure-activity relationship (SAR) for the indole derivative class defined by the structure of **1a** revealed that the *N*-substituent R^1^ is crucial for the biological activity (Table 1 and Table S1). Generally, amide and ether groups as *N*-alkyl substituents in the R^1^ position were well tolerated (Table 1, entries 1-16), while ester or ketone substitutions of the *N*-alkyl group decreased the activity (Table 1, entry 17 and Table S1). Comparison of aromatic ether substituents at the *N*-alkyl group revealed that small, electron-withdrawing groups on the phenyl ring were favorable (Table 1, entries 2-8). Introduction of hydrophobic and lipophilic moieties on the indole nitrogen improved the potency (Table 1, entries 11-13). Aromatic residues with a halogen in *para*-position were favorable (Table 1, entries 18-21), while most heterocycles were less active or inactive (entry 16 and Table S1). Comparison of compounds **1l** and **1m** (Table 1, entries 12-13) showed that substituting the R^4^ position with bromine led to a 50-fold increase in activity (see also Table 1, entries 20-21). Introducing substituents in R^2^, R^3^ and R^6^ positions generally resulted in less active compounds (Table 1, entries 22-23 and Table S1). Interestingly, only the trifluoroacetyl group in R^5^ position gave good IC50 values, which suggests that a small and strong electron-withdrawing group in this position is beneficial for biological activity (Table 1, entries 9 and 24, Table S1). Ultimately, compound **1l**, termed epoxykynin (Table 1, entry 12), which combines a hydrophobic *N*-cycloheptyl acetamide at R^1^ with bromine and trifluoroacetyl residues at R^4^ and R^5^ positions, was identified as the most potent derivative with an IC50 value of 36 ± 15 nM.

**Table 1:**
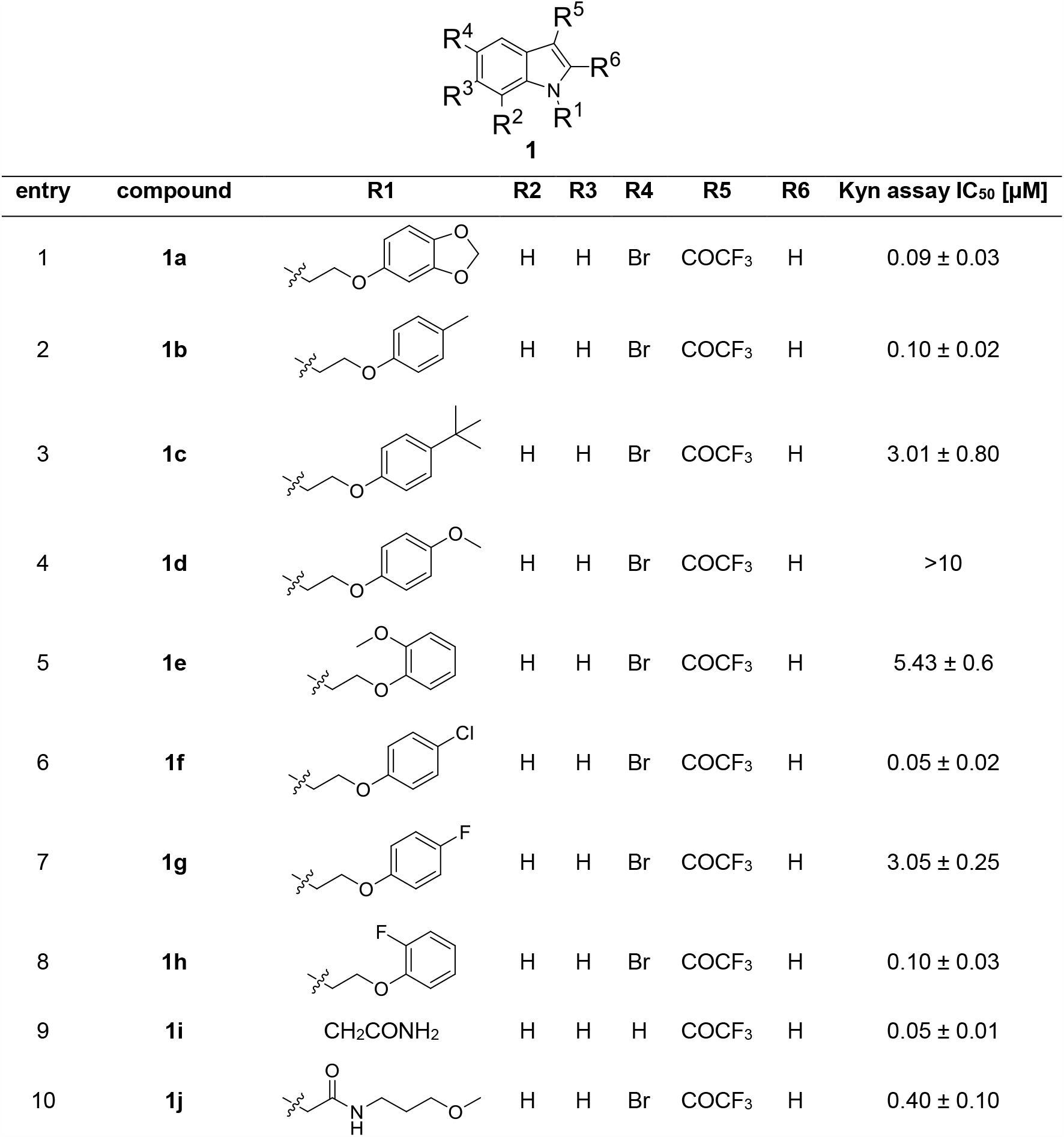
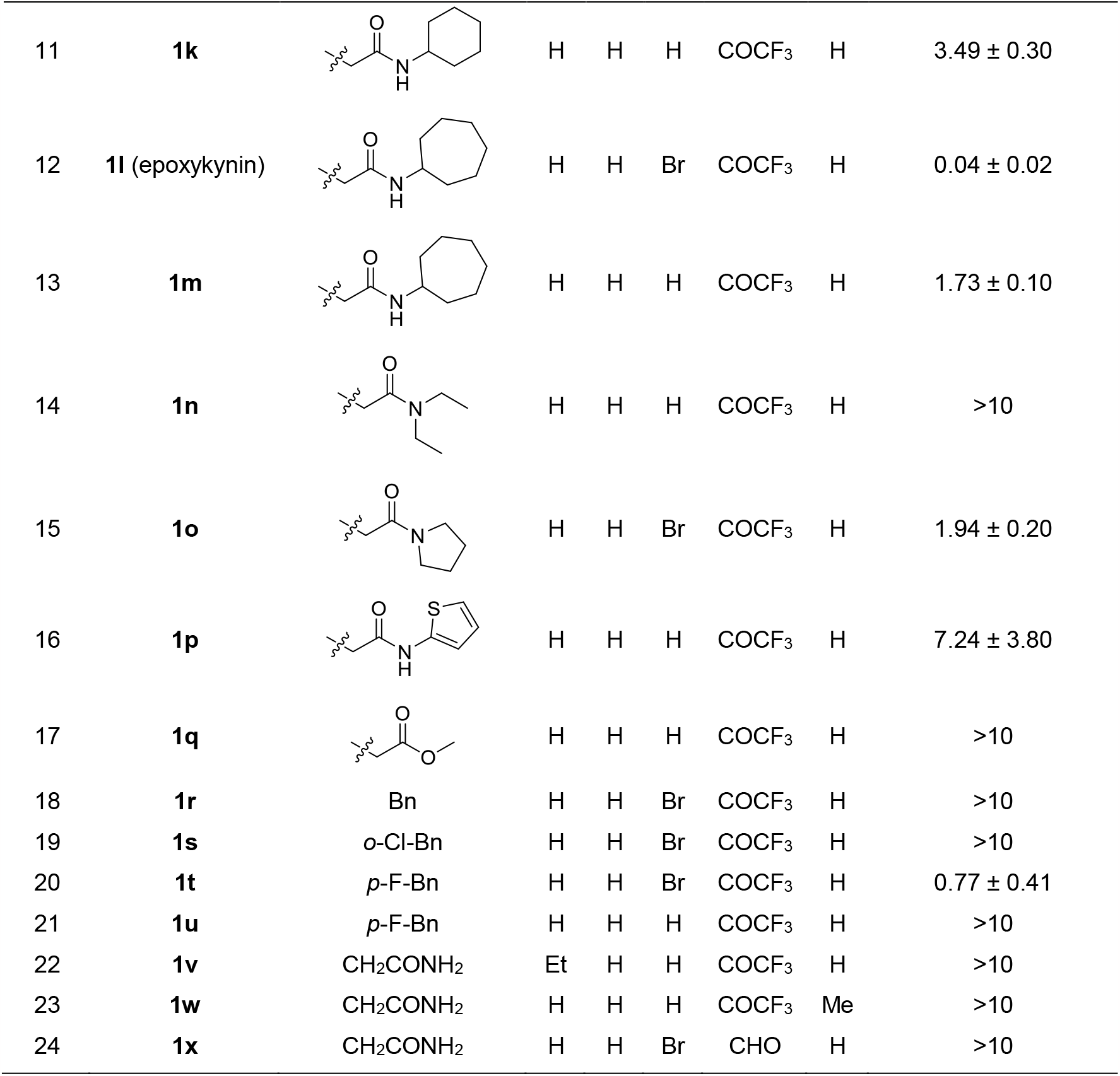
Structure-activity relationship (SAR) determined for selected compounds for reduction of Kyn levels (see also Table S1). IC50 values were determined in BxPC-3 cells using the automated Kyn assay. Data are mean values ± SD (n≥3).

| 1 |  |  |  |  |  |  |  |  |  |
| --- | --- | --- | --- | --- | --- | --- | --- | --- | --- |
| entry | compound | R1 | R2 | R3 | R4 | R5 | R6 | Kyn assay IC<sub>50</sub> [μM] | |
| 1 | 1a | | H | H | Br | COCF<sub>3</sub> | H | 0.09 ± 0.03 | |
| 2 | 1b | | H | H | Br | COCF<sub>3</sub> | H | 0.10 ± 0.02 | |
| 3 | 1c | | H | H | Br | COCF<sub>3</sub> | H | 3.01 ± 0.80 | |
| 4 | 1d |  | H | H | Br | COCF<sub>3</sub> | H | >10 |  |
| 5 | 1e | | H | H | Br | COCF<sub>3</sub> | H | 5.43 ± 0.6 | |
| 6 | 1f | | H | H | Br | COCF<sub>3</sub> | H | 0.05 ± 0.02 | |
| 7 | 1g | | H | H | Br | COCF<sub>3</sub> | H | 3.05 ± 0.25 | |
| 8 | 1h | | H | H | Br | COCF<sub>3</sub> | H | 0.10 ± 0.03 | |
| 9 | 1i | CH<sub>2</sub>CONH<sub>2</sub> | H | H | H | COCF<sub>3</sub> | H | 0.05 ± 0.01 | |
| 10 | 1j | | H | H | Br | COCF<sub>3</sub> | H | 0.40 ± 0.10 | |

| entry | compound | R1 | R2 | R3 | R4 | R5 | R6 | Kyn assay IC<sub>50</sub> [μM] |
| --- | --- | --- | --- | --- | --- | --- | --- | --- |
| 11 | **1k** |  | H | H | H | COCF<sub>3</sub> | H | 3.49 ± 0.30 |
| 12 | **1l** (epoxykynin) |  | H | H | Br | COCF<sub>3</sub> | H | 0.04 ± 0.02 |
| 13 | **1m** |  | H | H | H | COCF<sub>3</sub> | H | 1.73 ± 0.10 |
| 14 | **1n** |  | H | H | H | COCF<sub>3</sub> | H | >10 |
| 15 | **1o** |  | H | H | Br | COCF<sub>3</sub> | H | 1.94 ± 0.20 |
| 16 | **1p** |  | H | H | H | COCF<sub>3</sub> | H | 7.24 ± 3.80 |
| 17 | **1q** |  | H | H | H | COCF<sub>3</sub> | H | >10 |
| 18 | **1r** | Bn | H | H | Br | COCF<sub>3</sub> | H | >10 |
| 19 | **1s** | *o*-Cl-Bn | H | H | Br | COCF<sub>3</sub> | H | >10 |
| 20 | **1t** | *p*-F-Bn | H | H | Br | COCF<sub>3</sub> | H | 0.77 ± 0.41 |
| 21 | **1u** | *p*-F-Bn | H | H | H | COCF<sub>3</sub> | H | >10 |
| 22 | **1v** | CH<sub>2</sub>CONH<sub>2</sub> | Et | H | H | COCF<sub>3</sub> | H | >10 |
| 23 | **1w** | CH<sub>2</sub>CONH<sub>2</sub> | H | H | H | COCF<sub>3</sub> | Me | >10 |
| 24 | **1x** | CH<sub>2</sub>CONH<sub>2</sub> | H | H | Br | CHO | H | >10 |

**Figure 1:**
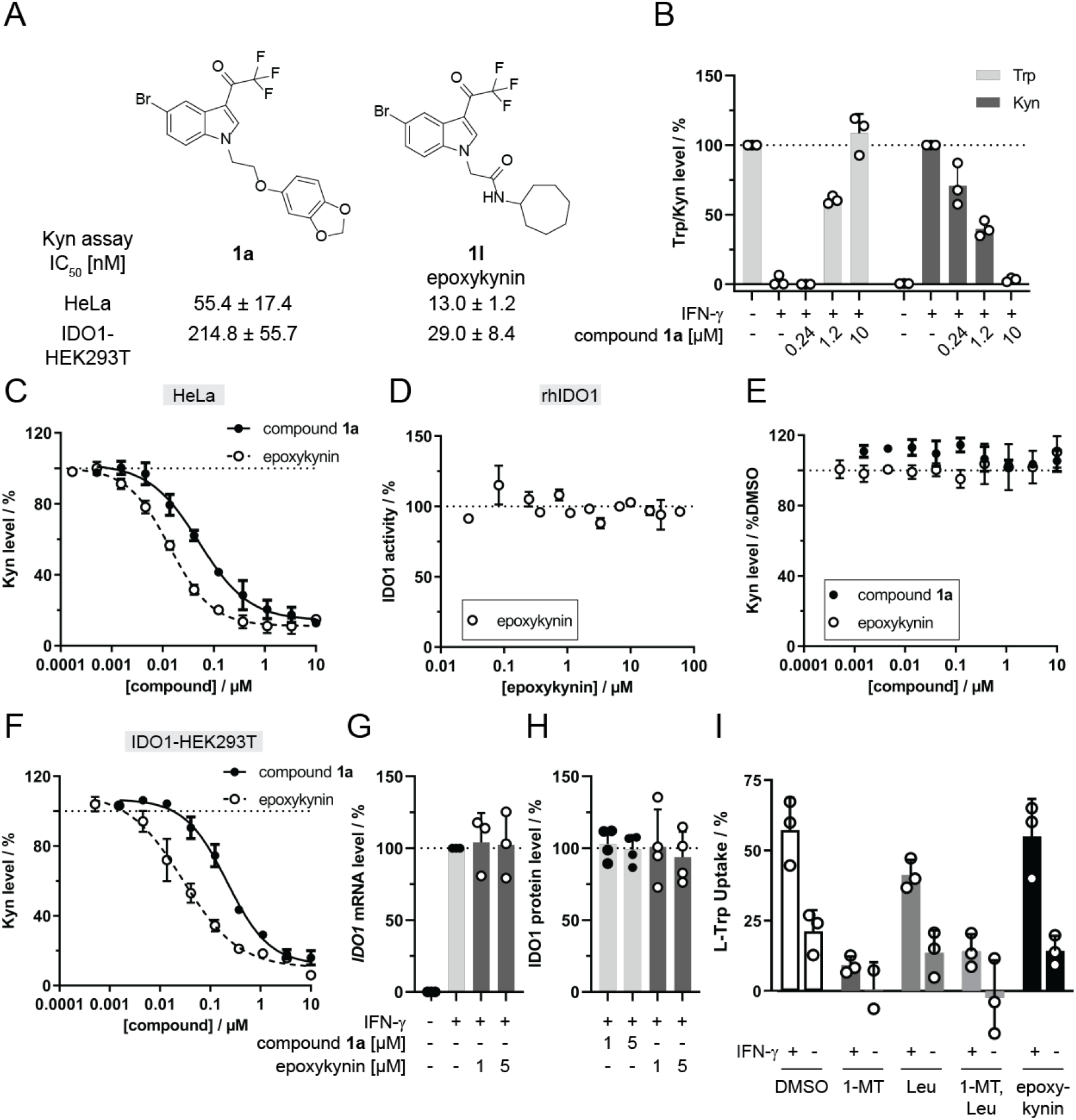
Reduction of cellular Kyn levels by compound **1a** and epoxykynin (**1l**) and influence on IDO1 expression and Trp uptake. A) Structures of the initial hit compound **1a** and the most potent compound epoxykynin (**1l**) and the respective IC50 values for Kyn reduction using the manual Kyn assay in HeLa and IDO1-expressing HEK293T cells. B) Trp/Kyn level in BxPC-3 cells upon treatment with IFN-γ, Trp and compound **1a**. Kyn and Trp levels were determined by LC-MS (mean values ± SD, n≥2). C) Kyn assay in HeLa cells. Cells were treated with IFN-γ, Trp and compounds for 48 h prior to measuring Kyn levels using *para*-dimethylaminobenzaldehyde (*p*-DMAB, mean values ± SD, n=3). D) *In vitro* IDO1 enzymatic activity. Purified IDO1 was treated with epoxykynin or DMSO for 40 min at 37°C prior to addition of Trp and incubation for 60 min at 37°C. Kyn levels were detected using *p*-DMAB (mean values ± SD, n=2). E) *IDO1* promoter-dependent reporter gene assay in HEK293T cells expressing firefly luciferase (Fluc) under the control of the *IDO1* promoter and constitutive *Renilla* luciferase expression (Rluc). Cells were treated with IFN-γ to induce Fluc expression and simultaneously with compounds for 48 h. Fluc values were normalized to the Rluc signal (mean values ± SD, n=3). F) Kyn assay in HEK293T cells transiently expressing human IDO1. Cells were treated with Trp and compounds for 24 h prior to measuring Kyn levels with *p*-DMAB (mean values ± SD, n=3). G) *IDO1* mRNA expression in HeLa cells that were treated with IFN-γ and epoxykynin or DMSO for 24 h prior to quantification of mRNA levels *via* qPCR (mean values ± SD, n=3). H) IDO1 protein levels in HeLa cells that were treated with IFN-γ and compound **1a** and epoxykynin or DMSO for 24 h prior to quantification of protein levels via immunoblotting (mean values ± SD, n=4). See also Figure S1 for complete blots. I) Trp uptake in BxPC-3 cells for epoxykynin. BxPC-3 cells were starved for Trp for 72 h and treated with IFN-γ for 24 h prior to addition 5 mM L-leucine (L-Leu), 1 mM 1-methyl-L-tryptophan (1-MT) or 5 μM epoxykynin for 30 min. Afterwards, 50 μM Trp was added and the Trp uptake after 30 min was quantified with HPLC-MS/MS (mean values ± SD, n=3).

To validate the screening results, the impact of **1a** on cellular Trp and Kyn levels was quantified *via* LC-MS as an orthogonal assay readout (Figure 1B). Treatment with IFN-γ stimulates the expression of IDO1 in Bx-PC3 cells and, thereby, increases Kyn levels and reduces Trp levels. In the presence of IFN-γ, compound **1a** dose-dependently inhibited Kyn production (Figure 1B). At 10 μM, hardly any Kyn was detectable, and the consumption of supplemented Trp increased in a concentration-dependent manner (Figure 1B). In HeLa cells, epoxykynin was approximately 4-fold more active than the initial hit compound **1a** (Figure 1C), with an IC50 value of 13.0 ± 1.2 nM, which is comparable to the IC50 determined for the screening assay in BxPC-3 cells of 36 ± 15 nM (Table 1, entry 12). Interestingly, epoxykynin did not affect the *in vitro* enzymatic activity of IDO1 (Figure 1D), and both, hit **1a** and epoxykynin did not alter *IDO1* expression as detected using a reporter gene under the control of the *IDO1* promoter (Figure 1E). Compound **1a** and epoxykynin decreased the Kyn levels in HEK293T cells that transiently express IDO1 in the absence of IFN-γ (Figure 1F), demonstrating that the *IDO1* promoter and signaling events upstream of the promoter are not modulated by the compound. In agreement with these findings, compound treatment did not alter the *IDO1* mRNA or IDO1 protein levels (Figure 1G-H). Epoxykynin may interfere with the uptake of the IDO1 substrate Trp and thereby reduce Kyn production. Large essential amino acids, such as Trp or leucine (Leu), can be imported by L-type amino acid transporters (LAT) which can be inhibited by saturating concentrations of Leu.^18,19^ Furthermore, Trp can be transported into the cell by IFN-γ-inducible tryptophanyl-tRNA synthetases (TrpRS),^20^ and treatment with the Trp analogue 1-methyl-L-tryptophan (1-MT) inhibits TrpRS.^20-22^ To analyze both uptake routes, BxPC-3 cells were starved for Trp in the presence or absence of IFN-γ prior to treatment with Leu, 1-MT and epoxykynin for 30 min (Figure 1I). As expected, IFN-γ increased Trp uptake due to upregulation of TrpRS and IDO1, thus leading to a higher demand for the IDO1 substrate Trp. Trp import was reduced by Leu and 1-MT, but not by epoxykynin. Hence, compounds **1a** and epoxykynin **1l** decrease cellular Kyn levels in the presence and absence of IFN-γ, but neither by reduction of IDO1 expression and inhibition of IDO1 enzymatic activity, nor by modulation of the uptake of the IDO1 substrate Trp.

### Epoxykynin reduces cellular Kyn levels by inhibition of sEH

Based on the structure-activity relationship analysis, affinity probes for chemical proteomics (pulldown) were generated. To this end, a Boc-protected amine-PEG4-alkyne linker was attached to epoxykynin and compound **1r** (Table 1, entry 18) to form precursors **2a** and **3a** respectively (Figure 2A) and removal of the Boc group yielded free amine probes **2b** and **3b**. Active probe **2a** decreased cellular Kyn levels, while negative probe **3a** did not (Figure 2B).

**Figure 2:**
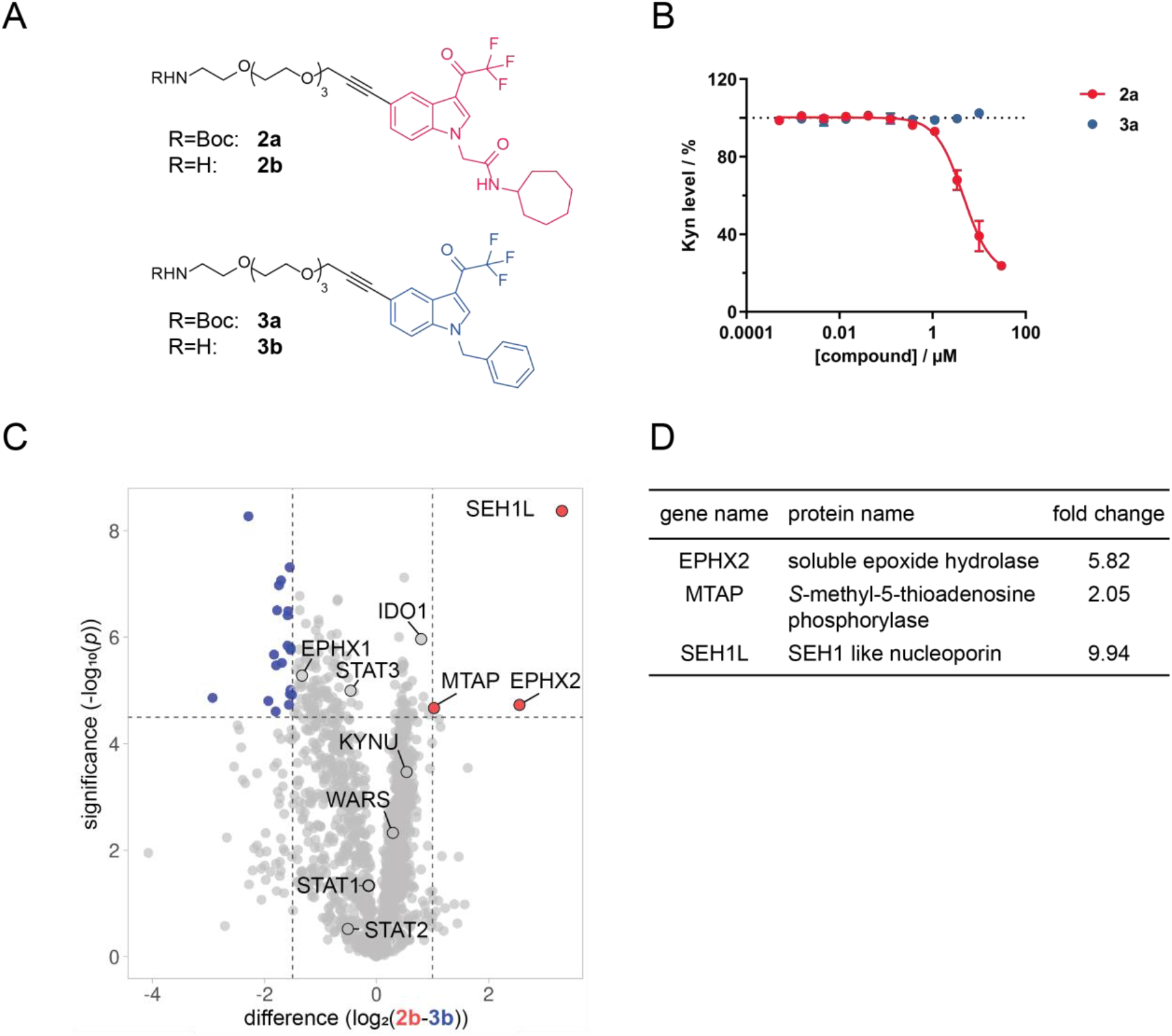
Target identification for epoxykynin. A) Structures of affinity probes **2a** and **2b** and control affinity probes **3a** and **3b**. B) Influence of the pulldown probes **2a** and **3a** on Kyn levels. HeLa cells were treated with IFN-γ, Trp and compounds for 48 h prior to measuring Kyn levels using *p*-DMAB (mean values ± SD, n=3). C) Volcano plot for proteins enriched using affinity-based chemical proteomics (pulldown) with probe **2b** (red) or control probe **3b** (blue) created with VolcaNoseR.23 The affinity probes **2b** and **3b** were immobilized on NHS-activated beads and incubated for 2 h at 4°C with lysate of HeLa cells that were treated with IFN-γ. Enriched proteins were analyzed using HRMS (n=2, N=4, FDR 0.01), representative replicate is shown, see also Figure S2. D) Proteins from C) that were significantly enriched with the affinity probe **2b**. For a complete list of enriched proteins see Table S2 and S3.

Subsequently, probes **2b** and **3b** were immobilized on NHS-activated beads for the affinity pulldown and incubated with HeLa cell lysate followed by HRMS analysis of bound proteins (Figure 2C). In total, 1,579 proteins were identified but only SEH1 like nucleoporin (SEH1L), methylthioadenosine phosphorylase (MTAP) and the soluble epoxide hydrolase (sEH, EPHX2) bound selectively to active affinity probe **2b** (Figure 2C-D and S2, Tables S2 and S3). Several additional proteins involved in the IDO1 pathway were found, namely IDO1, STAT1-3,^24^ TrpRS (WARS)^21^ and kynureninase (KYNU),^25^ but were not significantly enriched by either of the affinity probes (Figure 2C and S2). Unlike sEH, both MTAP and SEH1L were also enriched using the control probe **3b** to a certain extent (Table S4). The CRAPome^26,27^ database for background contaminants in affinity pulldowns lists SEH1L and other nucleoporins as frequently detected under control conditions. Thus, SEH1L might non-specifically interact with the sample matrix, whereas sEH and MTAP may represent *bona fide* interactors. To explore MTAP as a possible target of epoxykynin, we employed HCT116 MTAP^(-/-)^ cells that transiently express IDO1 (Figure S3). Epoxykynin inhibited cellular Kyn production in both HCT116 wildtype (wt) and MTAP knockout cells with comparable IC50 values of 20.7 ± 11.5 nM and 20.5 ± 15.9 nM, respectively. Therefore, epoxykynin does not suppress Kyn levels *via* modulation of MTAP.

In the pulldown, sEH was selectively enriched by affinity probe **2b** in comparison to probe **3b**. Moreover, epoxykynin competed with probe **2b** for binding to sEH (Figure S2). Direct binding of epoxykynin to sEH was analyzed using nano differential scanning fluorimetry (nanoDSF, Figure 3A and S4). Treatment of purified sEH with epoxykynin dose-dependently shifted the thermal denaturation temperature *T*m of sEH by 5.5 ± 1.5°C at 10 μM, suggesting binding of epoxykynin to sEH.

**Figure 3:**
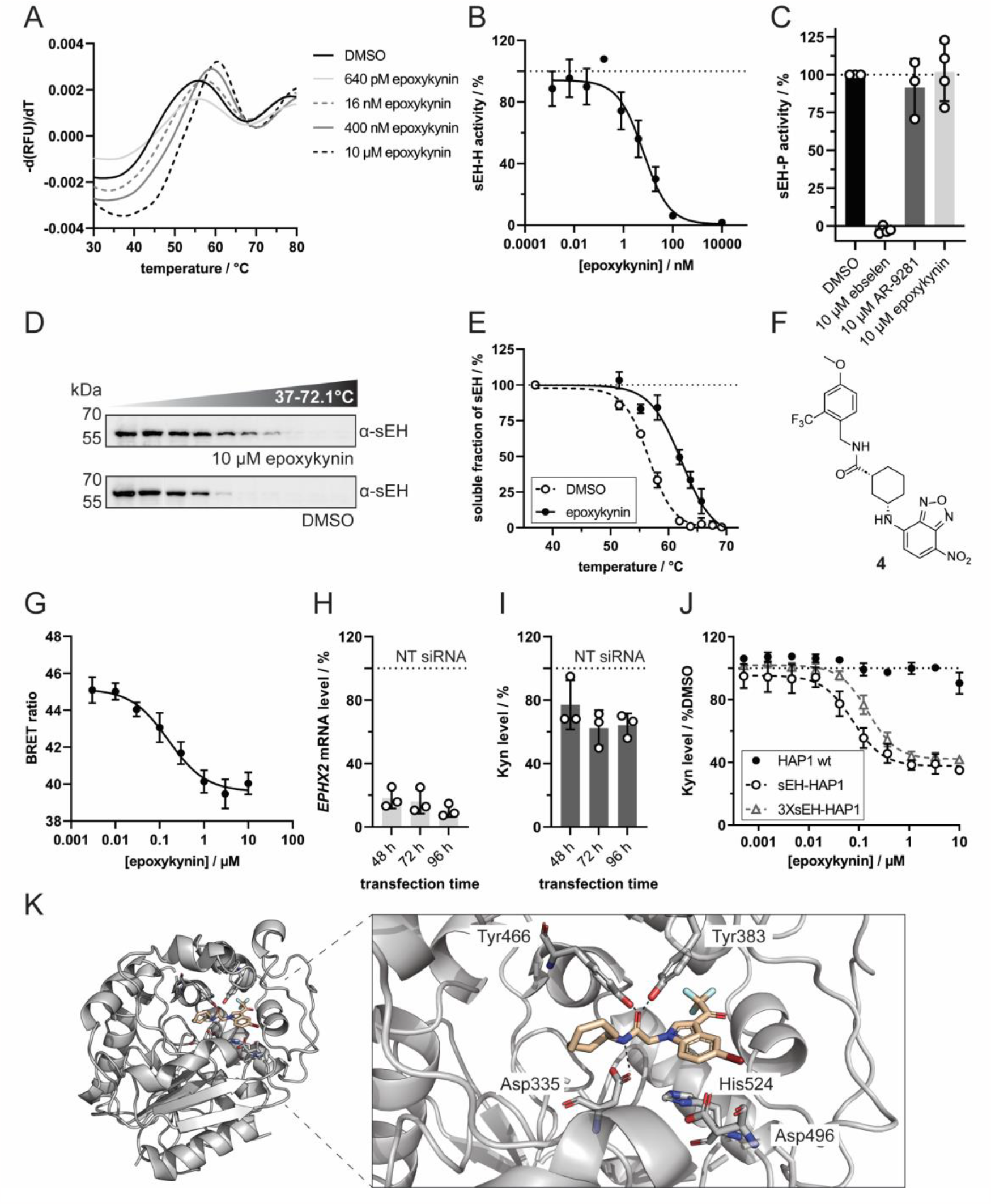
Epoxykynin binds to sEH *in vitro* and *in cellulo* and inhibits the C-terminal lipid epoxide hydrolase activity (sEH-H). A) Dose-dependent binding of epoxykynin to sEH as detected using nanoDSF. Purified sEH was treated with epoxykynin or DMSO for 10 min at room temperature prior to detection of the intrinsic tryptophan/tyrosine fluorescence upon thermal denaturation. Representative first derivatives of melting curves are shown (n=4, see also Figure S4). B) Dose-dependent inhibition of sEH-H by epoxykynin. The epoxide hydrolase activity of purified sEH (sEH-H) was measured by means of the conversion of the fluorogenic sEH-H substrate PHOME upon treatment with epoxykynin (mean values ± SD, n=3). See also Figure S5. C) Epoxykynin does not inhibit sEH-P. The phosphatase activity of purified sEH (sEH-P) was measured by means of an AttoPhos-based assay upon treatment with epoxykynin or AR9281 and ebselen as controls. Representative curves are shown (n=3, see also Figure S6). D) Cellular thermal shift assay (CETSA) for sEH in Jurkat cells. Cells were treated with 10 μM epoxykynin or DMSO for 15 min prior to heat treatment and cell lysis. Soluble proteins were analyzed using immunoblotting. Representative immunoblots are shown (n=3, see also Figure S7). E) Thermal stability of sEH upon compound treatment. Quantification of sEH band intensities from D (mean values ± SD, n=3). F) Structure of the fluorescent sEH ligand **4**.28 G) Dose-dependent displacement of the fluorescent tracer **4** by epoxykynin in HEK293T cells expressing NanoLuc-sEH. HEK293T cells that transiently express NanoLuc-sEH were treated with 60 nM of tracer **4** and epoxykynin for 5 h prior to determination of the bioluminescence resonance energy transfer (BRET) ratio (mean values ± SD, n=5). H and I) Knockdown (KD) of sEH decreases Kyn levels. HeLa cells were transfected with 50 nM non-targeting (NT) or *EPHX2*-targeting siRNA for 48-96 h and treated with Trp and IFN-γ for 48 h prior to detection of *EPHX2* mRNA (H) and Kyn levels with *p*-DMAB (I) (mean values ± SD, n=3). J) Overexpression of sEH in HAP1 cells. HAP1 cells were transiently transfected with different amounts of sEH expression plasmid (1 μg (sEH-HAP1) or 3 μg (3XsEH-HAP1) plasmid DNA per 96-well plate) prior to treatment with epoxykynin, Trp and IFN-γ for 48 h. Kyn levels were quantified using *p*-DMAB (mean values ± SD, n=3). K) Crystal structure of epoxykynin bound to human sEH-H (aa 228-547, pdb 8qzd). Epoxykynin (wheat sticks) binds to the sEH-H active site (grey cartoon and sticks) and is stabilized by polar interactions with the two stabilizing residues Tyr383 and Tyr466 and with Asp335 of the catalytic triad Asp335-Asp496-His524 (indicated by the dotted black lines). The amino acids in the active site are labeled with the three-letter code. Heteroatoms of the ligand and amino acid side chains are depicted in red (oxygen), blue (nitrogen), dark red (bromine) and cyan (fluorine). Amino acids 497-500 are omitted for clarity.

sEH is a bifunctional enzyme bearing a C-terminal lipid epoxide hydrolase domain (sEH-H) and an N-terminal lipid phosphatase moiety (sEH-P),^29-31^ both separated by a proline-rich linker.^32^ While the biological role of sEH-P still remains elusive,^33^ sEH-H was identified as part of the CYP epoxygenase branch of the arachidonic acid (AA) cascade.^34^ It catalyzes the hydrolysis of epoxy fatty acids, such as epoxyeicosatrienoic acids (EETs), to their corresponding vicinal diols.^32^ Thus, sEH-H plays a vital role in the catabolism of bioavailable epoxides, contributing to the detoxification of xenobiotics and regulation of signaling molecules.^35,36^ sEH-H is involved in a variety of disease-associated pathways, e.g. hypertension,^37^ diabetes,^38^ inflammation^39^ and cancer progression.^40^ Therefore, several classes of sEH inhibitors have been developed which mimic epoxides, such as ureas, carbamates and amides.^41,42^ Additionally, the synergism of sEH and proteins assigned to the other two major pathways of the AA cascade has been exploited to design multitarget dual-inhibitors.^43,44^

Epoxykynin inhibited purified sEH-H very potently with an IC50 value of 6.7 ± 3.2 nM (Figure 3B). Compounds less active in the Kyn level reduction assay were also weaker inhibitors of sEH-H (Table S5). Additional analysis of inhibition of the phosphatase domain sEH-P by means of an AttoPhos-based assay^33^ showed that the sEH-H and sEH-P inhibitor ebselen^33^, but not the sEH-H inhibitors AR9281^45^ and epoxykynin impeded the activity of sEH-P in this assay (Figure 3C). These findings demonstrate that epoxykynin selectively targets the C-terminal hydrolase domain of sEH, but not the N-terminal phosphatase domain. To show target engagement in cells, the thermal stability of sEH was investigated by means of a cellular thermal shift assay (CETSA, Figure 3D-E) in Jurkat cells, which have high sEH levels.^46-48^ Compared to the DMSO control, epoxykynin increased the melting temperature of sEH by 5.9 ± 1.2°C which correlates well with the Δ*T*m determined in the nanoDSF experiment (Figure 3A). For further confirmation of cellular target engagement by nano bioluminescence resonance energy transfer (nanoBRET), the fluorescent sEH ligand **4** (Figure 3F) was exposed to HEK293T cells that transiently express NanoLuc-sEH (Figure 3G)^28^. Transfer of energy by the bioluminescent NanoLuc-sEH donor to the fluorescent acceptor **4** can only occur in close proximity.^49^ Thus, a decrease in the BRET ratio indicates displacement of tracer **4** from the NanoLuc-sEH protein. Treatment with epoxykynin dose-dependently decreased the BRET ratio with an IC50 value of 159.4 ± 44.0 nM (Figure 3G). These findings demonstrate that epoxykynin binds and inhibits sEH-H both *in vitro* and in cells, but does not impair the catalytic activity of sEH-P. In addition, depletion of sEH in HeLa cells with siRNA resulted in sEH knockdown of 82 ± 5%, 84 ± 6% and 90 + 3% after 48, 72 and 96 h, respectively (Figure 3H). This partial knockdown decreased Kyn levels by 23 ± 13%, 38 ± 9% and 36 ± 6% after 48, 72 and 96 h, respectively (Figure 3I). Hence, depletion of sEH phenocopies treatment with epoxykynin. We noticed that epoxykynin did not affect IFN-γ-induced Kyn production in HAP1 cells, whereas IDO1 inhibitors like epacadostat reduced Kyn levels (Figure S8). HAP1 cells express hardly any sEH^46-48^ which explains the inactivity of epoxykynin in this cell line and makes it particularly useful for sEH overexpression studies. Accordingly, treatment of HAP1 cells that transiently express sEH with epoxykynin inhibited Kyn production with an IC50 value of 68.4 ± 9.7 nM (Figure 3J). Tripling the amount of transfected plasmid DNA shifted the IC50 to 159.7 ± 49.1 nM (Figure 3J). These findings prove sEH as the target of epoxykynin that mediates the reduction in Kyn levels.

A co-crystal structure of epoxykynin with human sEH-H was obtained to confirm the binding mode of the compound (Figure 3K, pdb 8qzd). The active site of sEH-H consists of the catalytic triad Asp335-Asp496-His524, as well as the two stabilizing residues Tyr381 and Tyr465 opposite of the catalytic triad serving as an oxyanion hole.^50^ The crystal structure revealed that epoxykynin binds to the catalytic center of sEH-H, occupying the binding site of the endogenous epoxide substrates. The ligand is stabilized by hydrogen bonds between the amide oxygen of epoxykynin and residues Tyr383 and Tyr466 and an additional hydrogen bond between the amide nitrogen of epoxykynin and Asp335 of the catalytic triad (Figure 3K). The co-crystal structure elucidates general trends in the SAR of the epoxykynin derivatives. The *N*-alkyl amide binds to the catalytic triad of sEH-H, while the lipophilic cycloheptane occupies a deep hydrophobic pocket. The constraint indole ring creates distance to the small, electron-withdrawing bromo and trifluoroacetyl substituents.

Structural alignments with previously published structures of sEH-H (Figure S9A-D, pdbs 1s8o, 5ai5, 3wke, 4hai) showed high overall similarity with root mean square deviation (RMSD) values below 1 Å. Besides the flexible C-terminus, there are minor structural deviations in a disordered loop of the cap domain of sEH-H between residues Ala411 and Lys421 (Figure S9E), indicating structural flexibility upon binding of different ligands. The two nitrogens of urea-derived sEH-H inhibitors (Figure S9F) act as hydrogen bond donors, while amide-based sEH-H inhibitors, such as epoxykynin (Figure 3K), only contain a single nitrogen as potential hydrogen bond donor to Asp335 of the catalytic triad. Yet, not all amide-based sEH-H inhibitors are stabilized by this hydrogen bond in the active site (Figure S9G-H).

### sEH regulates IDO1 protein levels

The metabolic level of reactive epoxides needs to be precisely balanced by biological systems,^35^ and as member of the arachidonic acid pathway sEH is involved in regulation of their levels. Kreiß *et al*. recently demonstrated that arachidonic acid metabolizing human 5-lipoxygenase regulates the expression of kynureninase,^51^ which suggests a possible functional link between the second branch of the AA acid cascade and the Kyn pathway. To investigate if IDO1 protein levels are regulated by sEH protein, we analyzed the IDO1 level after depletion (Figure 4A) or overexpression of sEH (Figure 4B) in HeLa cells. Knockdown of sEH decreased cellular IDO1 levels by 35 ± 10% after 72 h (Figure 4A), while overexpression of sEH increased IDO1 levels after 48 h, but not after 24 h (Figure 4B). In line with these findings, Zhou *et al*. found increased *IDO1* mRNA levels in sEH-overexpressing HCT116 cells^52^ which links sEH to regulation of Kyn levels through IDO1. When, instead of live cells, lysate of IFN-γ-stimulated BxPC-3 cells was treated with epoxykynin, in contrast to treatment with the IDO1 inhibitor epacadostat, Kyn levels were not reduced (Figure 4C). However, pre-treatment of BxPC-3 cells with either epoxykynin or *EPHX2*-targeting siRNA decreased Kyn levels in lysate (Figure 4D). These findings indicate that acute inhibition of sEH in cell lysate is not sufficient for Kyn reduction and a cross-talk between both pathways may operate *in cellulo* since sEH modulates the Kyn pathway and alters IDO1 protein.

**Figure 4:**
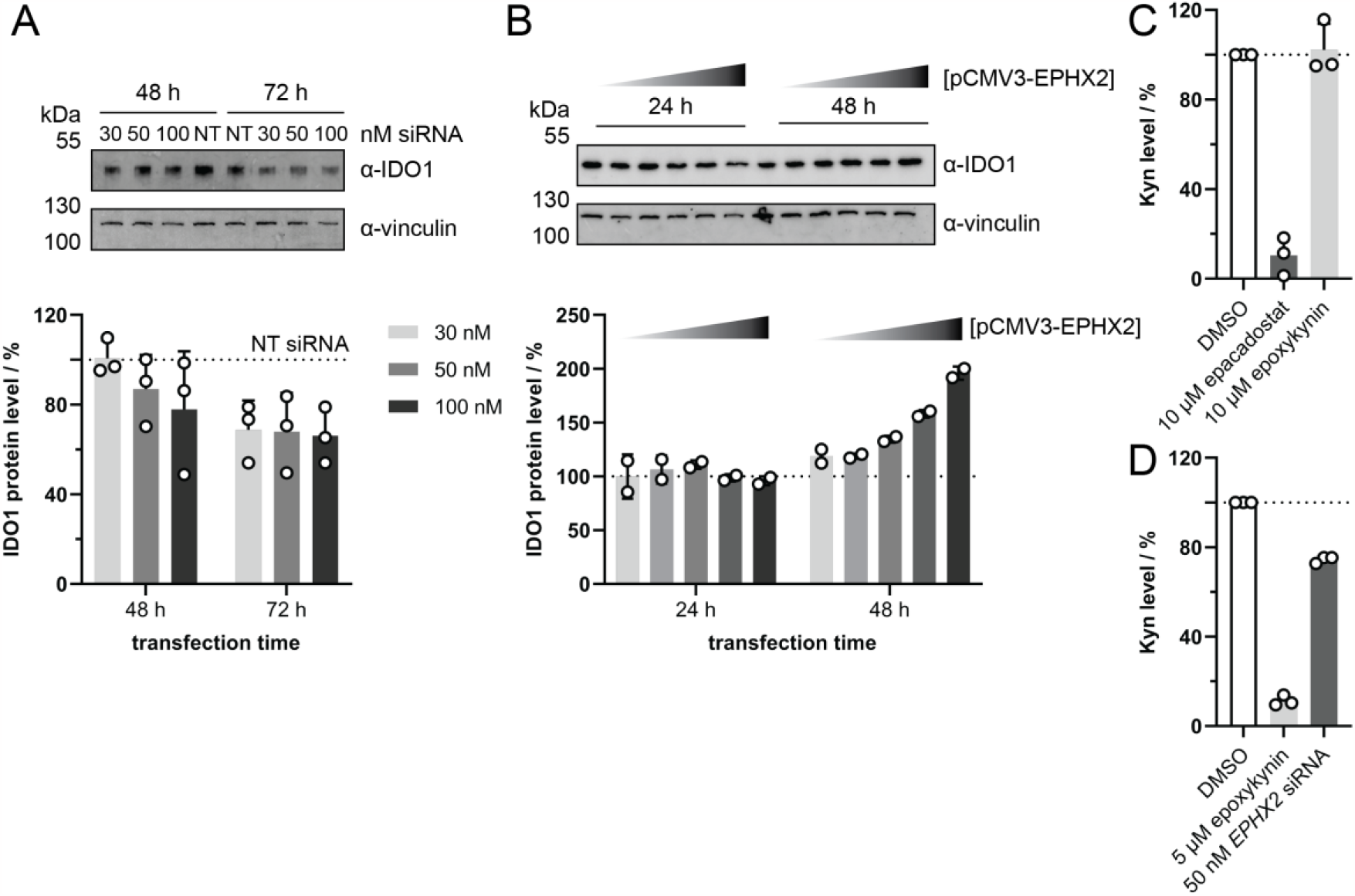
sEH cross-talks with the Kyn pathway, thereby modulating cellular IDO1 and Kyn levels. A) Knockdown (KD) of sEH. HeLa cells were transfected with non-targeting (NT) or *EPHX2*-targeting siRNA for 48-72 h and treated with IFN-γ for 48 h prior to quantification of IDO1 protein levels *via* immunoblotting (mean values ± SD, n=3). See also Figure S10 for complete blots. B) Overexpression of sEH. HeLa cells were transfected with empty vector or pCMV3-EPHX2 for 24-48 h and treated with IFN-γ for 48 h prior to quantification of IDO1 protein levels *via* immunoblotting (mean values ± SD, n=2). See also Figure S11 for complete blots. C-D) Kyn assay in cell lysate. C) BxPC-3 cells were treated with IFN-γ for 24 h prior to cell lysis. The lysate was treated with 10 μM epacadostat, 10 μM epoxykynin or DMSO for 45 min prior to detection of Kyn levels (mean values ± SD, n=3). D) BxPC-3 cells were treated with 5 μM epoxykynin, 50 nM *EPHX2*-targeting siRNA or DMSO and IFN-γ for 48 h prior to cell lysis and detection of Kyn levels (mean values ± SD, n=3).

Kyn is an endogenous agonist of the ligand-activated aryl hydrocarbon receptor (AhR).^53^ Activation of AhR induces IDO1 expression in antigen-presenting cells (APCs) or IDO1-positive cancer cells, promoting long-term immune tolerance.^54,55^ AhR antagonists can decrease plasma levels of AA in mice, while, vice versa, Kyn administration increases AA levels.^56^ Furthermore, the AA metabolite 12(R)-hydroxy-5(Z),8(Z),10(E),14(Z)-eicosatetraenoic acid (12(*R*)-HETE) activates AhR.^57^ Considering these findings, the connection between sEH overexpression and increased IDO1 levels reported here and noted previously^57^ may be due to an AhR-mediated feedback loop. This hypothesis is supported by the fact that inhibition of sEH by epoxykynin in cell lysate is not sufficient to inhibit Kyn production in lysates, indicating a possible functional cross-regulation of sEH and IDO1 expression by AhR. Since the AA derivative 12(*R*)-HETE acts as an AhR agonist,^57^ other eicosanoids may also bind to AhR, such as the sEH substrates epoxyeicosatrienoic acids (EETs) and dihydroxyeicosatrienoic acids (DHETs).^58^

sEH has been identified as a promising target for medicinal chemistry programs, and two sEH-targeting compounds have progressed to clinical trials, but have not found regulatory approval,^59-61^ such that new approaches are highly desirable. To this end, compound identification by means of a less biased cell-based assays may open up new advantageous alternatives to existing approaches. In conclusion, by means of a phenotypic screening for modulators of cellular Kyn production and subsequent target identification and validation, we have identified the potent sEH inhibitor epoxykynin. The compound engages with sEH in cells and inhibits sEH-H, resulting in reduced Kyn levels in cells expressing IDO1. Our data suggest a functional link between the third branch of the AA cascade and the Kyn pathway. Since the direct IDO1 inhibitor epacadostat recently failed clinical trials, alternative approaches to reduce Kyn levels are coveted, and inhibition of the Kyn pathway by modulating sEH may open up novel opportunities to revert cancer-related immune suppression.

## Author Contributions

S.Z. and H.W. designed the research. L.D., E.H., J.S., C.D.C.L.G. and K.H. performed the biological experiments. C.D. and S.T. synthesized the pulldown probes. J.H.M.E. produced recombinant protein for crystallization studies. A.K. solved the co-crystal structure. P.J. analyzed the data of the pulldown experiment. A.P. performed computational cluster analysis for the automated Kyn assay. S.S. adapted the conditions for the Kyn assay to an automated high-throughput format and performed the analysis of high-throughput data. S.K. supervised the crystallography. E.P. supervised protein production and nanoBRET studies. L.D., S.Z. and H.W. wrote the manuscript. All authors discussed the results and commented on the manuscript.

## Acknowledgements

This work was co-funded by the European Union (Drug Discovery Hub Dortmund (DDHD), EFRE-0200481) and Innovative Medicines Initiative (grant agreement number 115489) resources of which are composed of financial contribution from the European Union’s Seventh Framework Programme (FP7/2007-2013) and EFPIA companies’ in-kind contribution. E.P. and K.H. thanks German Research Foundation for financial support (E.P. DFG; SFB1039 A07; PR1405/13-1) and K.H. HI 2351/1-1). Dr. Axel Pahl and the compound management and screening center (COMAS) in Dortmund is acknowledged for performing the high-throughput screening and computational cluster analysis. We also thank Jens Warmers and Malte Metz for assistance with synthesis and HPLC-MS/MS-based experiments, as well as Christine Nowak and Laura Dragun for general support in the lab. We are grateful for Dr. Michael Grigalunas’s support during manuscript writing. We acknowledge Prof. Dr. Dieter Steinhilber and Maximilian Molitor for valuable scientific discussions. We thank the beamline scientists and local contacts at beamline X06SA, SLS Villigen for their assistance and support.

