## supporting information for "Discovery of the sEH Inhibitor Epoxykynin as Potent Kynurenine Pathway Modulator"

#### 1. Supplementary Data

##### 1.1 Supporting Tables

**Table S1:** Structure-activity relationship (SAR) for reduction of Kyn levels (related to Table 1). $IC_{50}$ values and Kyn assay inhibitions were determined in BxPC-3 cells using the automated Kyn assay. $IC_{50}$ values are mean values (mean values ± SD, n≥3) and Kyn assay inhibition values were determined as a single point measurement at 7.1 µM. Table is continued on following pages.

| entry | Kyn assay $IC_{50}$ [µM] | Kyn assay inhibition [%] | entry | Kyn assay $IC_{50}$ [µM] | Kyn assay inhibition [%] |
|---|---|---|---|---|---|
| 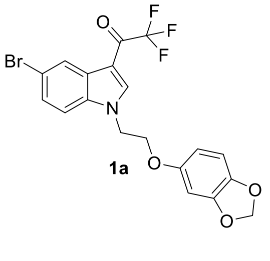 **1a** | 0.09 ± 0.03 | 95 | 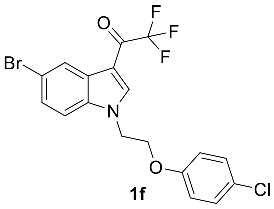 **1f** | 0.05 ± 0.02 | 98 |
| 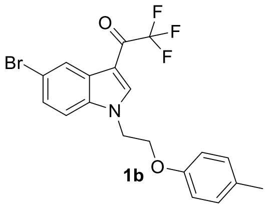 **1b** | 0.10 ± 0.02 | 83 | 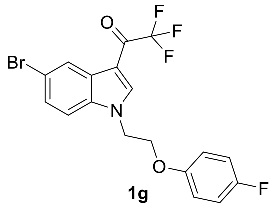 **1g** | 3.05 ± 0.25 | 100 |
| 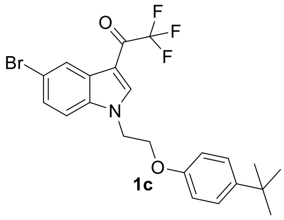 **1c** | 3.01 ± 0.80 | 20 | 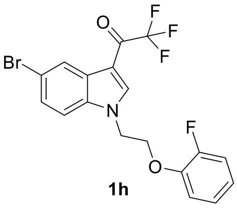 **1h** | 0.10 ± 0.03 | 99 |
| 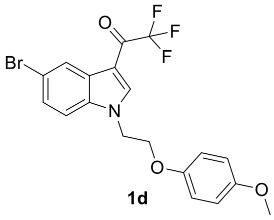 **1d** | >10 | 12 | 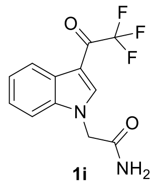 **1i** | 0.05 ± 0.01 | 112 |
| 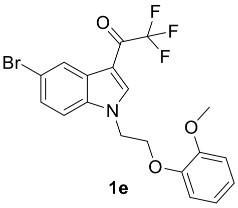 **1e** | 5.43 ± 0.6 | 54 | 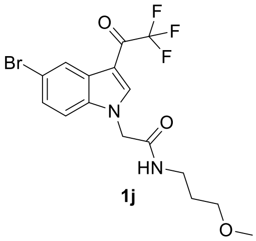 **1j** | 0.40 ± 0.10 | 95 |

| entry | Kyn assay $IC_{50}$ [μM] | Kyn assay inhibition [%] | entry | Kyn assay $IC_{50}$ [μM] | Kyn assay inhibition [%] |
|---|---|---|---|---|---|
| 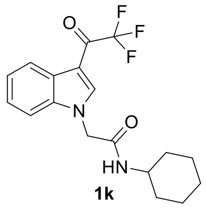 1k | 3.49 ± 0.30 | 41 | 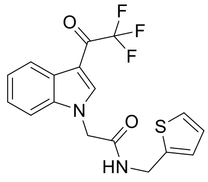 1p | 7.24 ± 3.80 | 49 |
| 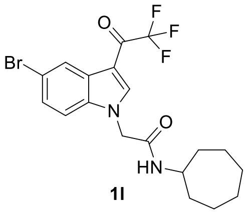 1l | 0.04 ± 0.02 | 98 | 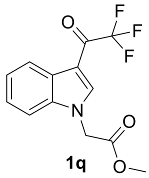 1q | >10 | -7 |
| 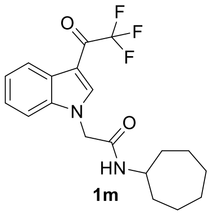 1m | 1.73 ± 0.10 | 55 | 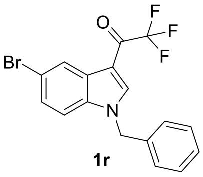 1r | >10 | 12 |
| 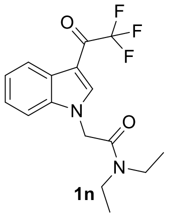 1n | >10 | 0 | 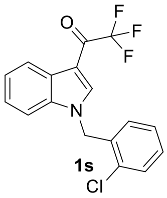 1s | >10 | 11 |
| 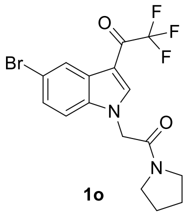 1o | 1.94 ± 0.20 | 70 | 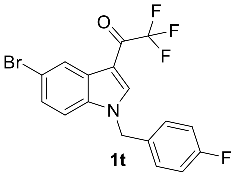 1t | 0.77 ± 0.41 | 67 |
| 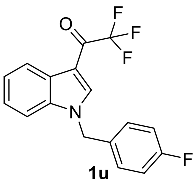 1u | >10 | 28 | 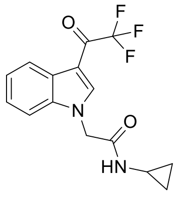 | >10 | 45 |

| entry | Kyn assay IC50 [µM] | Kyn assay inhibition [%] | entry | Kyn assay IC50 [µM] | Kyn assay inhibition [%] |
|---|---|---|---|---|---|
| 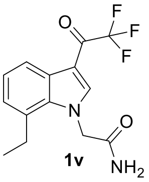 **1v** | >10 | 2 | 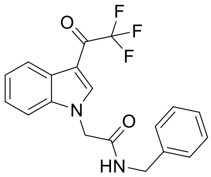 | >10 | 3 |
| 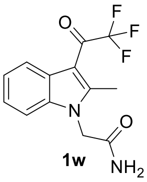 **1w** | >10 | -36 | 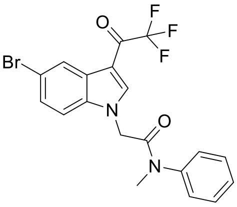 | 6.65 ± 2.6 | 12 |
| 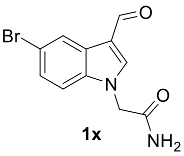 **1x** | >10 | -3 | 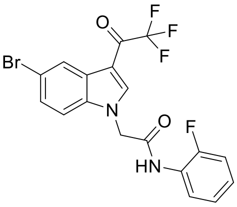 | 2.27 ± 0.6 | 52 |
| 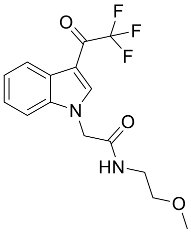 | >10 | 0 | 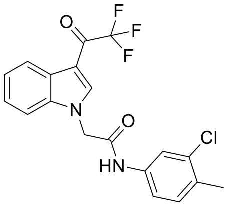 | 0.753 ± 0.4 | 85 |
| 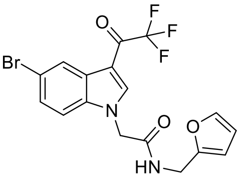 | 0.564 ± 0.12 | 93 | 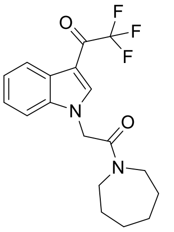 | >10 | 16 |
| 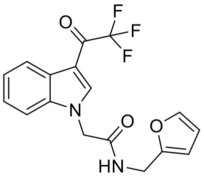 | 9.12 ± 1.1 | 34 | 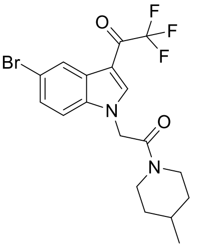 | 0.343 ± 0.04 | 96 |

| entry | Kyn assay IC50 [µM] | Kyn assay inhibition [%] | entry | Kyn assay IC50 [µM] | Kyn assay inhibition [%] |
|---|---|---|---|---|---|
| 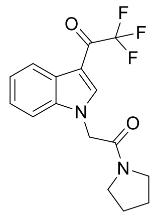 | >10 | -23 | 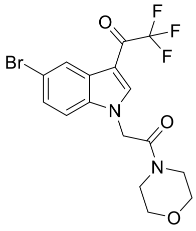 | >10 | -8 |
| 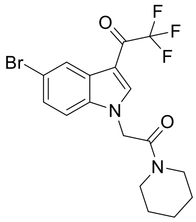 | 1.43 ± 0.2 | 88 | 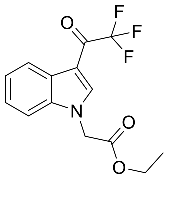 | >10 | -9 |
| 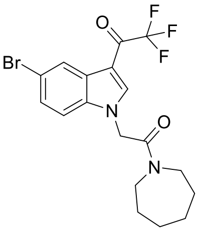 | 1.23 ± 0.2 | 88 | 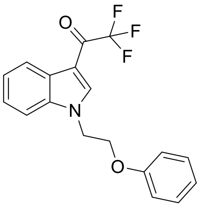 | 1.18 ± 0.7 | 84 |
| 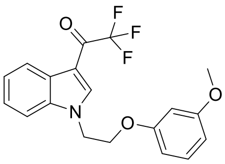 | >10 | -11 | 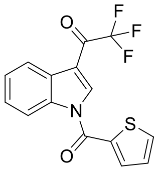 | not determined | 1 |
| 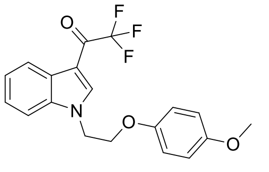 | >10 | 18 | 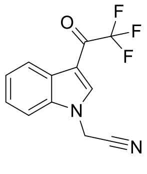 | >10 | 10 |
| 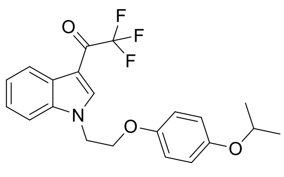 | 0.493 ± 0.12 | 96 | 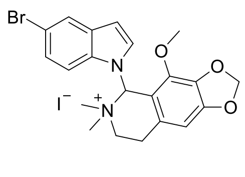 | not determined | 3 |

| entry | Kyn assay $IC_{50}$ [µM] | Kyn assay inhibition [%] | entry | Kyn assay $IC_{50}$ [µM] | Kyn assay inhibition [%] |
|---|---|---|---|---|---|
| 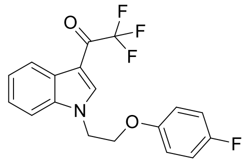 | 2.05 ± 0.1 | 108 | 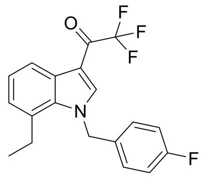 | >10 | -4 |
| 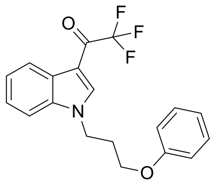 | >10 | 10 | 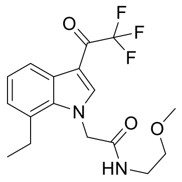 | >10 | -9 |
| 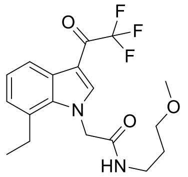 | >10 | -23 | 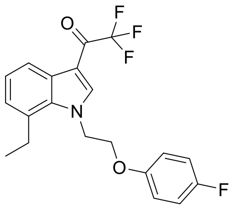 | 1.12 ± 0.4 | 77 |
| 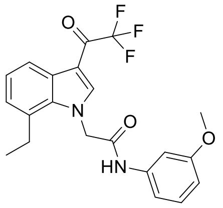 | >10 | -13 | 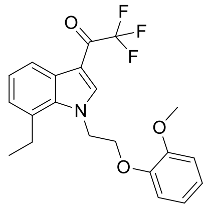 | >10 | 47 |
| 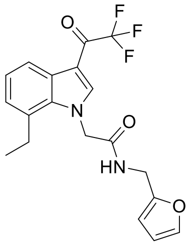 | >10 | 37 | 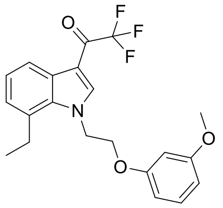 | 3.89 ± 2.4 | 74 |
| 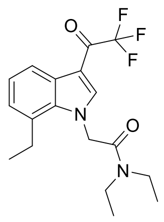 | >10 | 4 | 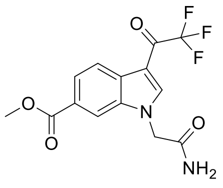 | >10 | -14 |

| entry | Kyn assay $IC_{50}$ [µM] | Kyn assay inhibition [%] | entry | Kyn assay $IC_{50}$ [µM] | Kyn assay inhibition [%] |
|---|---|---|---|---|---|
| 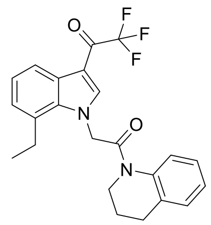 | >10 | 23 | 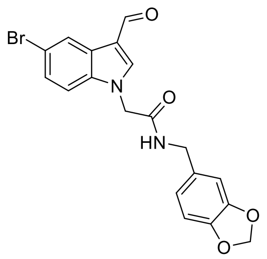 | not determined | -19 |
| 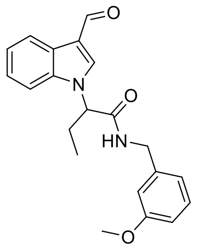 | 1.99 ± 0.4 | 71 | 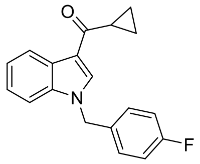 | 7.18 ± 1.4 | 59 |
| 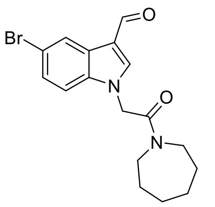 | not determined | -22 | 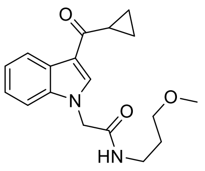 | >10 | -7 |
| 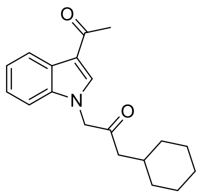 | >10 | -11 | 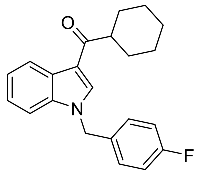 | 3.94 ± 0.7 | 21 |
| 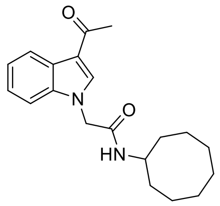 | >10 | -18 | 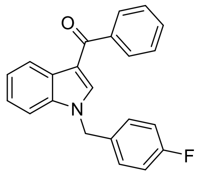 | >10 | 0 |
| 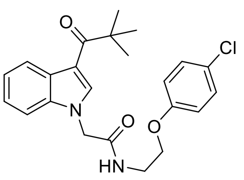 | 2.14 ± 0.1 | 70 | 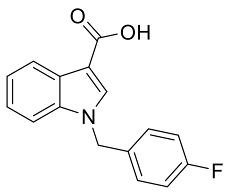 | >10 | 7 |

| entry | Kyn assay $IC_{50}$ [μM] | Kyn assay inhibition [%] | entry | Kyn assay $IC_{50}$ [μM] | Kyn assay inhibition [%] |
|---|---|---|---|---|---|
| 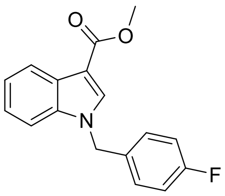 | 9.32 ± 1.0 | 42 | 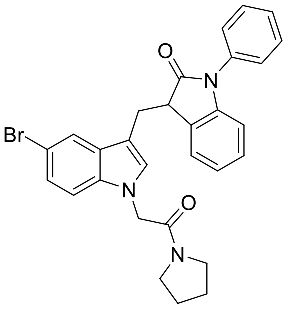 | not determined | 36 |
| 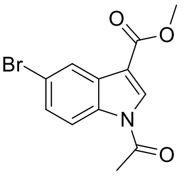 | not determined | -18 | 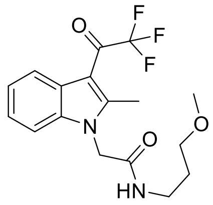 | >10 | -2 |
| 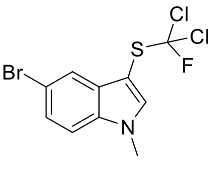 | not determined | 42 | 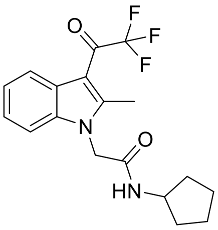 | 4.29 ± 1.6 | 63 |
| 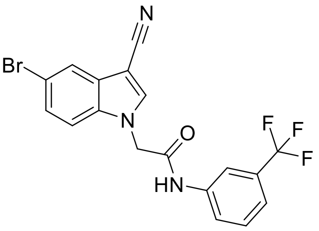 | not determined | -30 | 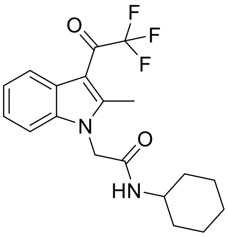 | 4.4 ± 0.4 | 12 |
| 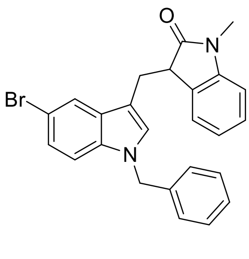 | not determined | 23 | 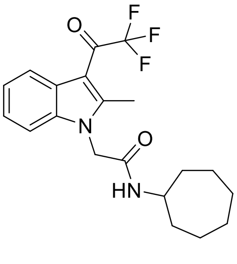 | 1.7 ± 0.1 | 82 |
| 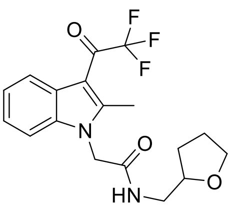 | >10 | 1 | 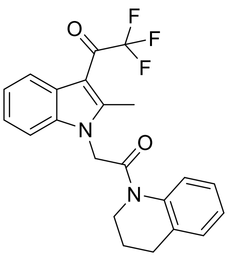 | >10 | -15 |

| entry | Kyn assay $IC_{50}$ [µM] | Kyn assay inhibition [%] | entry | Kyn assay $IC_{50}$ [µM] | Kyn assay inhibition [%] |
|---|---|---|---|---|---|
| 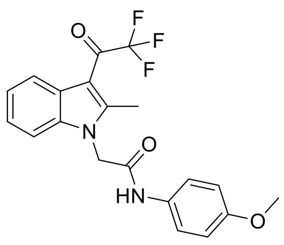 | >10 | -2 | 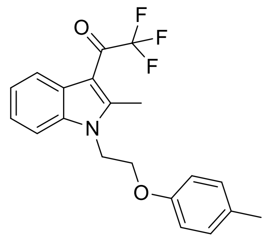 | >10 | -22 |
| 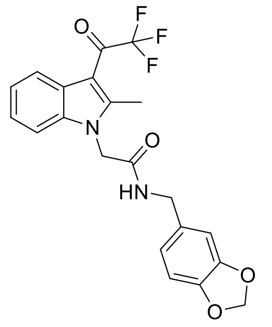 | >10 | -4 | 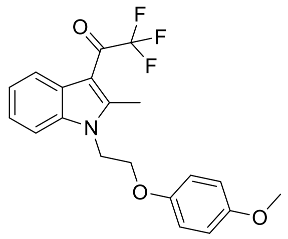 | 0.275 ± 0.08 | 97 |
| 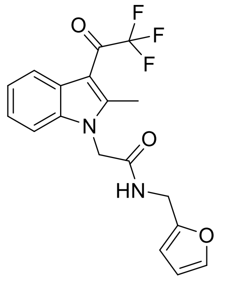 | >10 | 23 | 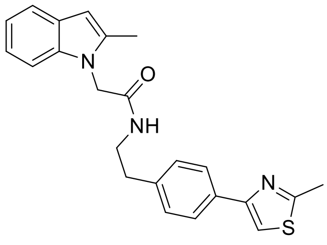 | 1.6 ± 0.0 | 87 |
| 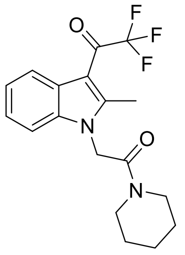 | >10 | -1 | 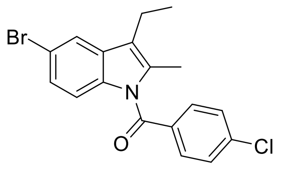 | not determined | -7 |
| 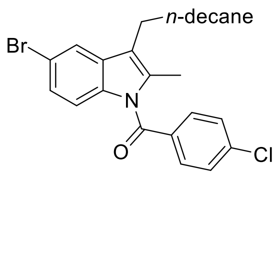 | not determined | 2 | 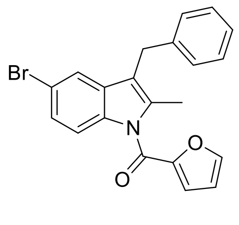 | >10 | 42 |
| 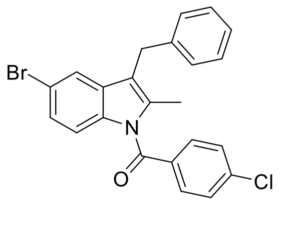 | not determined | 12 | 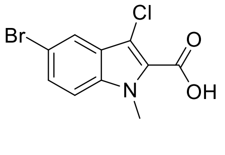 | not determined | 4 |

| entry | Kyn assay IC$_{50}$ [µM] | Kyn assay inhibition [%] | entry | Kyn assay IC$_{50}$ [µM] | Kyn assay inhibition [%] |
|---|---|---|---|---|---|
| 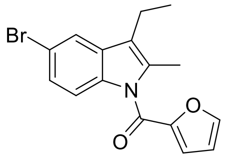 | not determined | 18 | 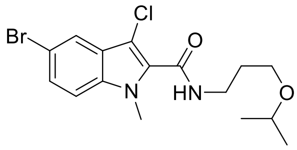 | not determined | -4 |

**Table S2:** Affinity-based chemical proteomics (pulldown). Enriched proteins on affinity probe **2b** as identified by HRMS (n=2, N=4, FDR 0.01). Proteins that were identified in both experimental replicates are highlighted in grey. Related to Figure 2.

| | n1, N=4 | | n2, N=4 | |
|---|---|---|---|---|
| **gene name** | **significance** (-log($p$)) | **difference** ($log_2$(2b-3b)) | **significance** (-log($p$)) | **difference** ($log_2$(2b-3b)) |
| EPHX2 | 2.14 | 1.99 | 4.74 | 2.54 |
| HIBCH | | | 4.47 | 1.11 |
| HPCAL1;HPCA | | | 3.54 | 1.63 |
| LYPLA2 | | | 4.32 | 1.14 |
| MTAP | 6.47 | 2.34 | 4.65 | 1.04 |
| RPL31 | | | 1.88 | 1.46 |
| SEH1L | 5.77 | 2.38 | 8.38 | 3.31 |
| TRA2B | 4.44 | 2.36 | | |

**Table S3:** Affinity-based chemical proteomics (pulldown). Enriched proteins on control probe **3b** as identified by HRMS (n=2, N=4, FDR 0.01). Proteins that were identified in both experimental replicates are highlighted in grey. Table is continued on the next pages. Related to Figure 2.

| | n1, N=4 | | n2, N=4 | |
|---|---|---|---|---|
| **gene name** | **significance** (-log(*p*)) | **difference** ($log_2$(2b-3b)) | **significance** (-log(*p*)) | **difference** ($log_2$(2b-3b)) |
| ABCB7 | | | 3.99 | -1.10 |
| ADK | 3.63 | -2.53 | 8.27 | -2.29 |
| AKAP8 | | | 1.95 | -4.07 |
| ALG5 | | | 2.87 | -1.30 |
| APOO | | | 5.01 | -1.54 |
| ARL6IP5 | | | 4.54 | -1.40 |
| ARL8B;ARL8A | | | 3.03 | -1.47 |
| ATL3 | | | 6.25 | -1.26 |
| ATP1B3 | | | 2.16 | -1.22 |
| ATP5EP2;ATP5E | | | 1.06 | -2.06 |
| ATP5F1 | | | 4.90 | -1.08 |
| ATP5J | 1.99 | -2.04 | | |
| ATP5J2;ATP5J2-PTCD1 | | | 4.92 | -1.08 |
| ATP5L | | | 4.75 | -1.05 |
| BCAP31 | | | 4.09 | -1.71 |
| BET1;DKFZp781C0425 | | | 1.61 | -1.71 |
| BST2 | | | 4.99 | -1.48 |
| C1GALT1 | | | 4.11 | -1.56 |
| CCDC134 | | | 1.98 | -1.64 |
| CD44 | | | 5.52 | -1.38 |
| CD97 | | | 4.32 | -1.08 |
| CDC42 | | | 3.03 | -1.19 |
| CERS2 | | | 1.23 | -1.78 |
| CHCHD3 | | | 4.86 | -1.25 |
| CISD1 | | | 5.84 | -1.59 |
| CISD2 | | | 5.47 | -1.79 |
| CLPTM1L | | | 5.41 | -1.34 |
| COMT | | | 4.56 | -1.18 |
| COX4I1 | | | 4.13 | -1.46 |
| COX5A | | | 4.64 | -1.11 |
| CSE1L | | | 5.70 | -1.04 |
| CYB5R3 | | | 4.03 | -1.14 |
| DAD1 | | | 2.98 | -1.72 |
| DEGS1 | | | 3.87 | -1.27 |
| DHCR7 | | | 5.03 | -1.17 |
| DHRS1 | | | 5.12 | -1.39 |
| DHRS3 | | | 1.78 | -1.73 |
| EDIL3 | | | 4.63 | -1.13 |
| ELOVL1 | | | 1.52 | -1.73 |
| ELOVL5 | | | 4.25 | -1.24 |
| EPHX1 | | | 5.28 | -1.33 |
| ERGIC1 | | | 5.45 | -1.18 |
| FADS1 | | | 6.51 | -1.37 |
| FADS2 | | | 5.42 | -1.08 |
| FAM105A | | | 3.82 | -1.31 |
| FKBP11 | 2.29 | -2.35 | 3.88 | -1.09 |
| FUNDC2 | 3.94 | -2.88 | 3.25 | -2.34 |
| GALNT2 | | | 4.70 | -1.22 |
| GLUD1; GLUD2 | 5.49 | -1.67 | | |
| GNB2 | | | 4.58 | -1.47 |
| GNG5 | | | 3.32 | -2.39 |
| GPR89A;GPR89B | | | 4.26 | -1.52 |
| GPX8 | 1.74 | -2.91 | 5.58 | -1.27 |
| HCCS | | | 4.07 | -1.51 |
| HEATR1 | | | 4.34 | -1.08 |
| HEBP1 | 5.95 | -2.58 | 7.07 | -1.71 |

| | n1, N=4 | | n2, N=4 | |
|---|---|---|---|---|
| **gene name** | **significance** (-log(*p*)) | **difference** ($log_2$(2b-3b)) | **significance** (-log(*p*)) | **difference** ($log_2$(2b-3b)) |
| HLA-A | | | 5.18 | -1.12 |
| HLA-B | | | 5.85 | -1.21 |
| HLA-C;HLA-Cw | | | 6.13 | -1.09 |
| HMOX2 | | | 2.90 | -1.19 |
| IKBIP | | | 3.59 | -1.09 |
| IMMT | | | 5.80 | -1.54 |
| ITPA | 7.11 | -2.85 | | |
| LCLAT1 | | | 4.33 | -1.19 |
| LMAN1 | | | 5.19 | -1.07 |
| LMAN2 | | | 4.80 | -1.93 |
| LMAN2L | | | 3.30 | -1.41 |
| LMF2 | | | 5.73 | -1.19 |
| MAGT1 | | | 3.07 | -1.27 |
| MBOAT7 | | | 3.91 | -1.07 |
| MCU | | | 4.73 | -1.13 |
| MGST1 | | | 7.32 | -1.55 |
| MLEC | | | 2.87 | -1.49 |
| MPC1;BRP44L | | | 3.22 | -1.41 |
| MPC2 | | | 2.09 | -2.13 |
| MPDU1 | | | 5.77 | -1.38 |
| MRRF | 1.72 | -2.19 | | |
| MT-ATP8 | | | 3.10 | -1.76 |
| MTCH2 | | | 4.98 | -1.12 |
| MT-CO2 | | | 5.61 | -1.26 |
| MTOR | | | 4.15 | -1.45 |
| MTX1 | | | 3.19 | -1.68 |
| NDC1 | | | 5.47 | -1.10 |
| NDUFA8 | | | 1.71 | -2.00 |
| NDUFAF2 | | | 3.64 | -1.78 |
| NDUFB11 | | | 3.56 | -2.54 |
| NDUFB3 | | | 2.91 | -1.38 |
| NDUFB5 | | | 2.58 | -1.33 |
| NT5C3B | | | 4.26 | -2.45 |
| PDCD6 | | | 4.93 | -1.04 |
| PFDN6 | 2.68 | -2.03 | | |
| PGM3 | | | 1.60 | -1.89 |
| PGRMC1 | | | 2.03 | -1.66 |
| PGRMC2 | 5.25 | -2.35 | 5.06 | -1.35 |
| PIGS | | | 4.47 | -1.31 |
| PIGT | | | 3.43 | -1.68 |
| PITPNB | 3.07 | -1.85 | | |
| POR | | | 2.96 | -1.15 |
| PRKDC | | | 6.33 | -1.04 |
| PSMD8 | 1.60 | -2.76 | | |
| PTGES2 | 2.46 | -1.91 | 4.52 | -1.10 |
| PTRH2 | | | 3.44 | -2.13 |
| RAB10 | | | 5.76 | -1.54 |
| RAB11B;RAB11A | | | 4.73 | -1.57 |
| RAB14 | | | 4.70 | -1.27 |
| RAB18 | | | 4.35 | -2.48 |
| RAB1A | | | 2.32 | -1.98 |
| RAB1B | | | 5.56 | -1.10 |
| RAB21 | | | 6.51 | -1.78 |
| RAB2A | | | 4.94 | -1.55 |
| RAB31;RAB22A | | | 4.19 | -1.76 |
| RAB5A | | | 1.75 | -1.80 |
| RAB5C | | | 5.12 | -1.35 |
| RAB6A; RAB6B | 3.54 | -1.75 | | |
| RAB7A | | | 5.39 | -1.45 |
| RAB8A | | | 4.36 | -1.10 |

| | n1, N=4 | | n2, N=4 | |
|---|---|---|---|---|
| **gene name** | **significance** (-log(*p*)) | **difference** ($log_2$(2b-3b)) | **significance** (-log(*p*)) | **difference** ($log_2$(2b-3b)) |
| RAP1B; RAP1A | 5.15 | -3.86 | 4.61 | -1.80 |
| RDH11 | | | 5.07 | -1.16 |
| RETSAT | | | 2.60 | -1.31 |
| RFT1 | | | 3.56 | -1.83 |
| RHOA | | | 1.44 | -1.98 |
| RHOG | | | 3.93 | -2.42 |
| S100A10 | | | 2.24 | -2.67 |
| SACM1L | | | 4.85 | -1.26 |
| SAMM50 | | | 6.98 | -1.74 |
| SAR1A | | | 2.86 | -1.74 |
| SCAMP3 | 4.90 | -2.82 | | |
| SCARB1 | | | 4.86 | -2.93 |
| SCCPDH | 3.90 | -1.94 | 2.87 | -1.17 |
| SCPEP1 | 3.14 | -2.01 | | |
| SEC11A | | | 6.78 | -1.37 |
| SEC22B | | | 4.69 | -1.43 |
| SEC61B | | | 6.41 | -1.59 |
| SEC61G | | | 5.67 | -1.83 |
| SEC63 | | | 1.62 | -2.21 |
| SECTM1 | | | 1.95 | -2.08 |
| SIGMAR1 | | | 2.04 | -2.18 |
| SLC30A7 | | | 2.68 | -1.65 |
| SMPD4 | | | 2.01 | -1.41 |
| SOAT1 | | | 4.68 | -1.17 |
| SPCS2 | | | 4.77 | -1.24 |
| SPCS3 | | | 4.57 | -1.17 |
| SPR | | | 6.49 | -1.58 |
| SSR3 | | | 1.19 | -1.65 |
| STT3A | | | 5.09 | -1.01 |
| SYPL1 | | | 2.75 | -1.36 |
| TAMM41 | | | 2.34 | -1.24 |
| TAPBP | | | 3.64 | -1.41 |
| TBL2 | 4.26 | -1.49 | 4.86 | -1.15 |
| TM9SF1 | | | 3.10 | -1.19 |
| TM9SF2 | | | 5.99 | -1.03 |
| TM9SF3 | | | 4.62 | -1.12 |
| TM9SF4 | | | 2.22 | -1.71 |
| TMED2 | | | 3.30 | -1.38 |
| TMED7-TICAM2;TMED7 | | | 4.25 | -1.18 |
| TMEM109 | | | 1.98 | -1.97 |
| TMEM205 | | | 1.66 | -2.06 |
| TMPO | | | 4.49 | -1.16 |
| TOMM20 | | | 1.36 | -2.28 |
| TOMM7 | | | 4.28 | -1.26 |
| TOR1AIP1 | | | 3.54 | -1.08 |
| TPP1 | | | 1.83 | -1.41 |
| TSPO | | | 4.84 | -1.35 |
| TUBGCP4 | | | 2.45 | -1.20 |
| VAPA | | | 4.92 | -1.51 |
| VAPB | | | 2.70 | -1.28 |
| VDAC1 | | | 5.11 | -1.31 |
| VMP1 | | | 1.88 | -1.51 |
| XPO7 | | | 6.26 | -1.04 |
| YIF1A | | | 3.68 | -1.24 |
| ZMPSTE24 | | | 5.52 | -1.69 |

**Table S4:** Affinity-based chemical proteomics (pulldown). LFQ intensities for proteins selectively enriched on affinity probe **2b** in comparison to control probe **3b** as determined by HRMS (n=2, N=4, FDR 0.01).

| | gene name | affinity probe 2b | | | | control probe 3b | | | |
|---|---|---|---|---|---|---|---|---|---|
| | | LFQ1 | LFQ2 | LFQ3 | LFQ4 | LFQ1 | LFQ2 | LFQ3 | LFQ4 |
| n1 | EPHX2 | 7,352,600 | 6,270,200 | 6,809,100 | 6,688,400 | 0 | 0 | 0 | 0 |
| | MTAP | 13,678,000 | 16,810,000 | 18,212,000 | 16,267,000 | 3,253,500 | 3,132,500 | 2,994,100 | 3,442,400 |
| | SEH1L | 75,267,000 | 88,492,000 | 108,110,000 | 80,662,000 | 15,245,000 | 18,745,000 | 16,230,000 | 17,053,000 |
| n2 | EPHX2 | 23,157,000 | 15787000 | 15518000 | 21115000 | 0 | 0 | 0 | 0 |
| | MTAP | 17,151,000 | 20,828,000 | 17,404,000 | 18,929,000 | 9,893,700 | 8,366,300 | 8,462,400 | 9,450,700 |
| | SEH1L | 281,169,984 | 254,340,000 | 262,070,000 | 260,070,000 | 24,289,000 | 25,395,000 | 28,607,000 | 28,273,000 |

**Table S5:** Structure-activity relationship (SAR) for reduction of Kyn levels (related toTable S1) and sEH-H inhibition. Kyn assay $IC_{50}$ values (mean values ± SD, n≥3) and Kyn assay inhibitions were determined in BxPC-3 cells using the automated Kyn assay. Inhibition of human sEH-H was measured by means of the conversion of the fluorogenic sEH-H substrate PHOME (mean values ± SD, n=3).

| entry | Kyn assay $IC_{50}$ | Kyn assay inhibition] | sEH-H assay $IC_{50}$ | sEH-H assay inhibition |
|---|---|---|---|---|
| 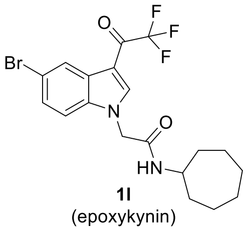 **1l** (epoxykynin) | 36.0 ± 15.0 nM | 98% | 6.7 ± 3.2 nM | 84% |
| 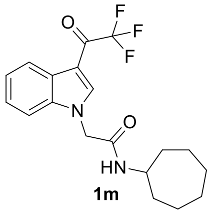 **1m** | 1.7 ± 0.1 µM | 55% | 50.1 ± 11.2 nM | 82% |
| 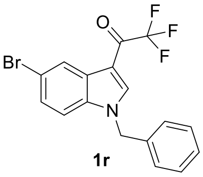 **1r** | >10 µM | 12% | >10 µM | n.d. |
| 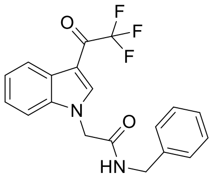 | >10 µM | 3% | 619.6 ± 71.2 nM | 44% |

**Table S6:** Data collection, processing, and refinement statistics for the sEH-H-epoxykynin complex (pdb 8qzd).

| | **8QZD** |
|---|---|
| **Data collection and reduction** | |
| Wavelength (Å) | 1.0 |
| Space group | $P22_12_1$ |
| Resolution range (Å) | 44.54 - 1.30 |
| Last resolution shell (Å) | 1.33 - 1.30 |
| Unit cell parameters | |
| a,b,c (Å) | 45.92, 80.31, 89.08 |
| α, β, γ (°) | 90, 90, 90 |
| Total number of observations | 518234 (21137) |
| Unique reflections | 80739 (3673) |
| Mosaicity (˚) | 0.20 |
| Multiplicity | 6.4 (5.8) |
| Mean I/σ(I) | 10.7 (1.5) |
| Completeness (%) | 99.6 (93.3) |
| $R_{merge}{}^{b}$ | 0.072 (1.071) |
| $R_{meas}{}^{c}$ | 0.085 (1.298) |
| $R_{pim}{}^{d}$ | 0.045 (0.723) |
| **Refinement** | |
| Resolution range (Å) | 44.54 - 1.30 |
| Number of reflections used | 76583 |
| Number of Free R flagged reflections | 4091 |
| $R_{cryst}{}^{e}$ | 0.16279 |
| $R_{free}{}^{f}$ | 0.18041 |
| rmsd Bond length (Å) | 0.012 |
| rmsd Bond angle (˚) | 1.926 |
| Ramachandran plot, residues in | |
| Most favored region (%) | 98.2 |
| Additionally allowed region (%) | 1.5 |
| Average B-factor ($Å^2$) | 18.140 |

[a]Values for the last resolution shell are in parentheses.

[b]$R_{merge} = \Sigma_{hkl}\Sigma_i|I_i(hkl)-\langle I(hkl)\rangle| / \Sigma_{hkl}\Sigma_i I_i(hkl)$, where I(hkl) is the intensity of reflection hkl

[c]$R_{meas} = \Sigma_{hkl}(n/(n-1))^{1/2}\Sigma_i|I_i(hkl)-\langle I(hkl)\rangle| / \Sigma_{hkl}\Sigma_i I_i(hkl)$

[d]$R_{pim} = \Sigma_{hkl}(1/(n-1))^{1/2}\Sigma_i|I_i(hkl)-\langle I(hkl)\rangle| / \Sigma_{hkl}\Sigma_i I_i(hkl)$

[e]$R_{cryst} = \Sigma_{hkl} ||F_{obs}|-|F_{calc}|| / \Sigma|F_{obs}|$

[f]$R_{free}$ is the cross-validation R-factor computed for the test set of unique reflections.

1.2 Supporting Figures

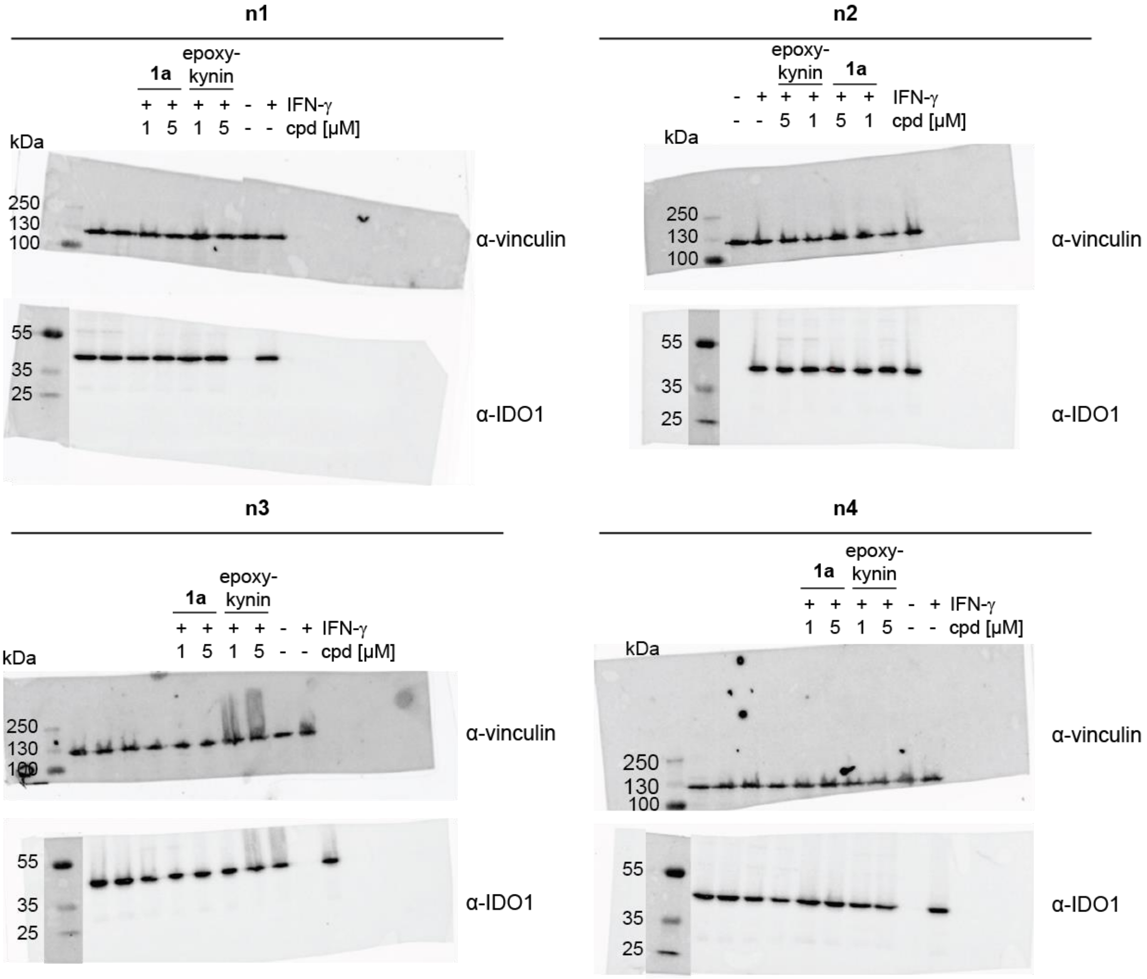

**Figure S1:** IDO1 protein levels in HeLa cells that were treated with IFN-γ and compound **1a** and epoxykynin or DMSO for 24 h prior to quantification of protein levels via immunoblotting. IDO1 protein bands were monitored in the IRDye800CW channel, the vinculin control bands and marker bands were monitored in the IRDye680RD channel. Related to Figure 1H.

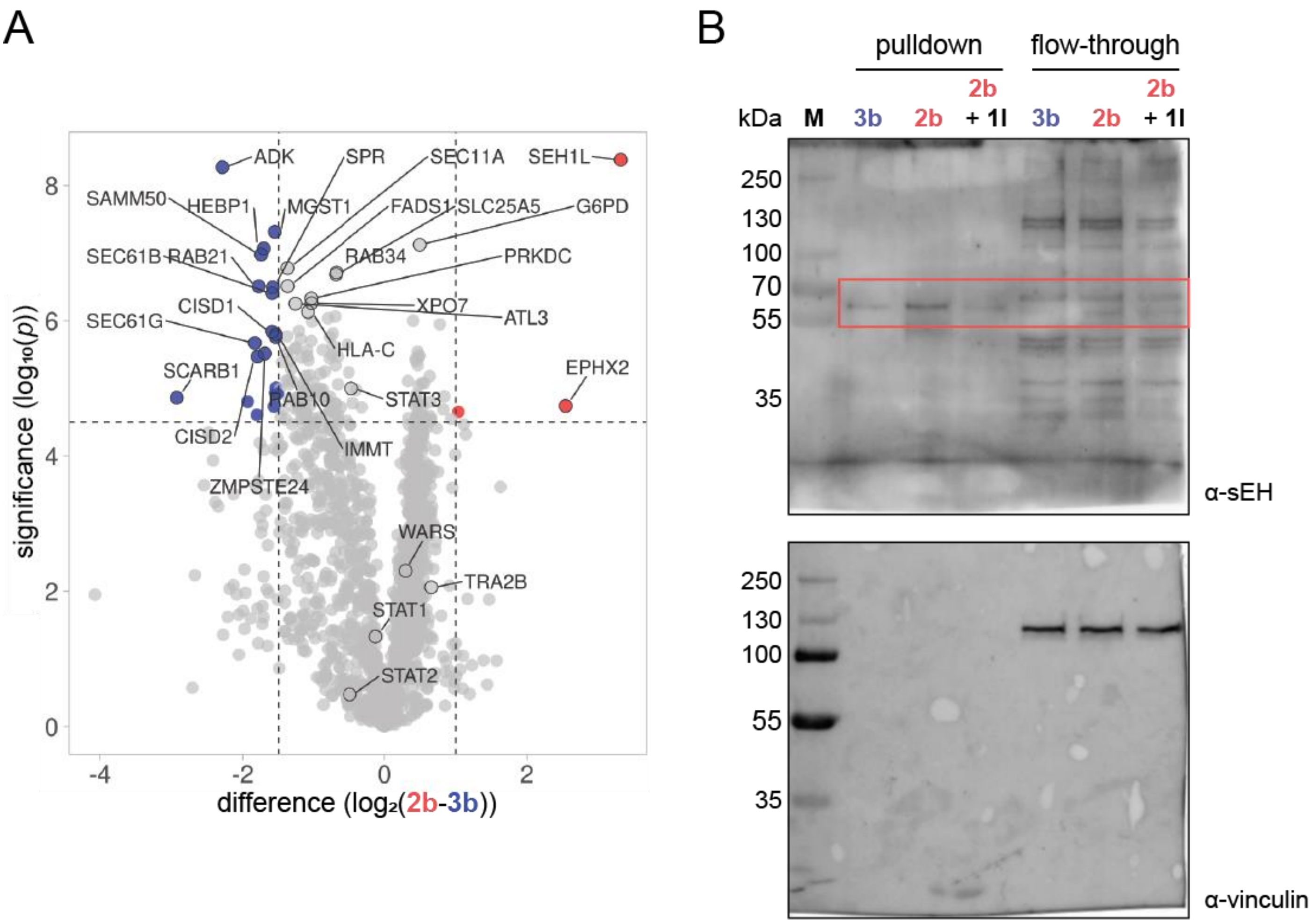

**Figure S2:** Affinity-based chemical proteomics (pulldown). A) The affinity probes **2b** and **3b** were immobilized on NHS-activated beads and incubated with HeLa cell lysate for 2 h at 4°C. Enriched proteins were analyzed using HRMS (n=2, N=4, FDR 0.01), representative replicate is shown, see also Table S2 and S3. Volcano plot for proteins enriched by probe **2b** (red) or probe **3b** (blue) created with VolcaNoseR.[1] B) Selective enrichment of EPHX2 (sEH) using the probe **2b** and competition with compound **1l** (epoxykynin). For the competition experiment, HeLa cell lysate was pre-incubated with 10 µM compound **1l** (epoxykynin) for 1 h at 4°C prior to incubation with the immobilized probes **2b** and **3b**. The enriched proteins were analyzed using immunoblot. sEH bands were monitored using an HRP-conjugated secondary antibody, the vinculin and marker (M) signal were monitored using an IRDye680RD secondary antibody. Related to Figure 2.

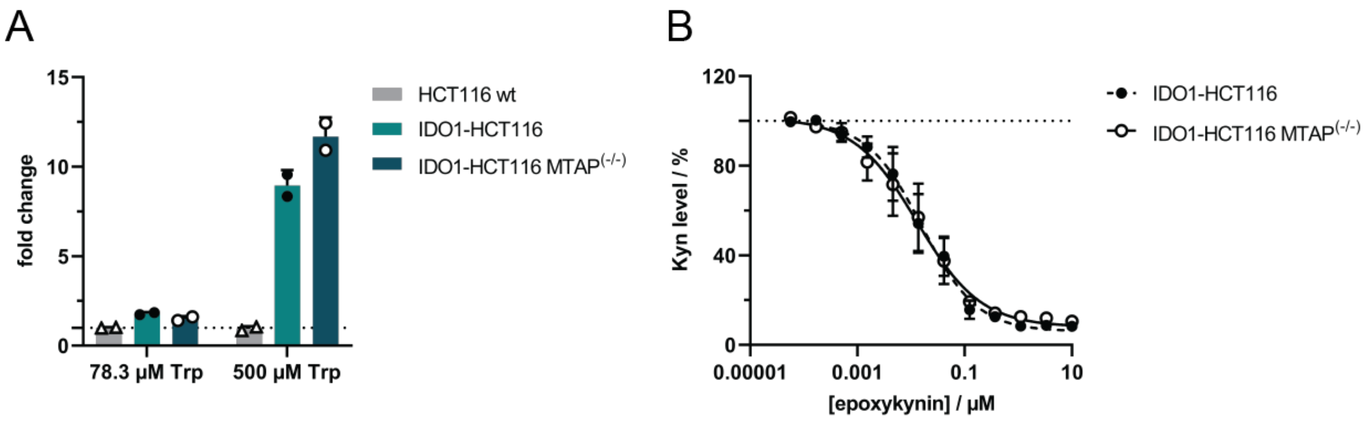

**Figure S3:** Devalidation of MTAP as target of epoxykynin using HCT116 MTAP$^{(-/-)}$ cells that transiently express IDO1. A) HCT116 wildtype (wt) and HCT116 MTAP (-/-) cells were transiently transfected with IDO1 expression construct for 24 h prior to addition of no (78.3 µM Trp already included in the medium) or 500 µM Trp for 36 h. Kyn levels were quantified using *p*-DMAB (mean values±SD, n=2). B) Kyn assay in HCT116 and HCT116 MTAP (-/-) cells that transiently express IDO1. Cells were transfected with IDO1 for 24 h prior to addition of 500 µM Trp and epoxykynin for 48 h. Kyn levels were quantified using *p*-DMAB (mean values ± SD, n=3). $IC_{50}$(IDO1-HCT116) = 20.7 ± 11.5 nM; $IC_{50}$(IDO1-HCT116 MTAP$^{(-/-)}$) = 20.5 ± 15.9 nM.

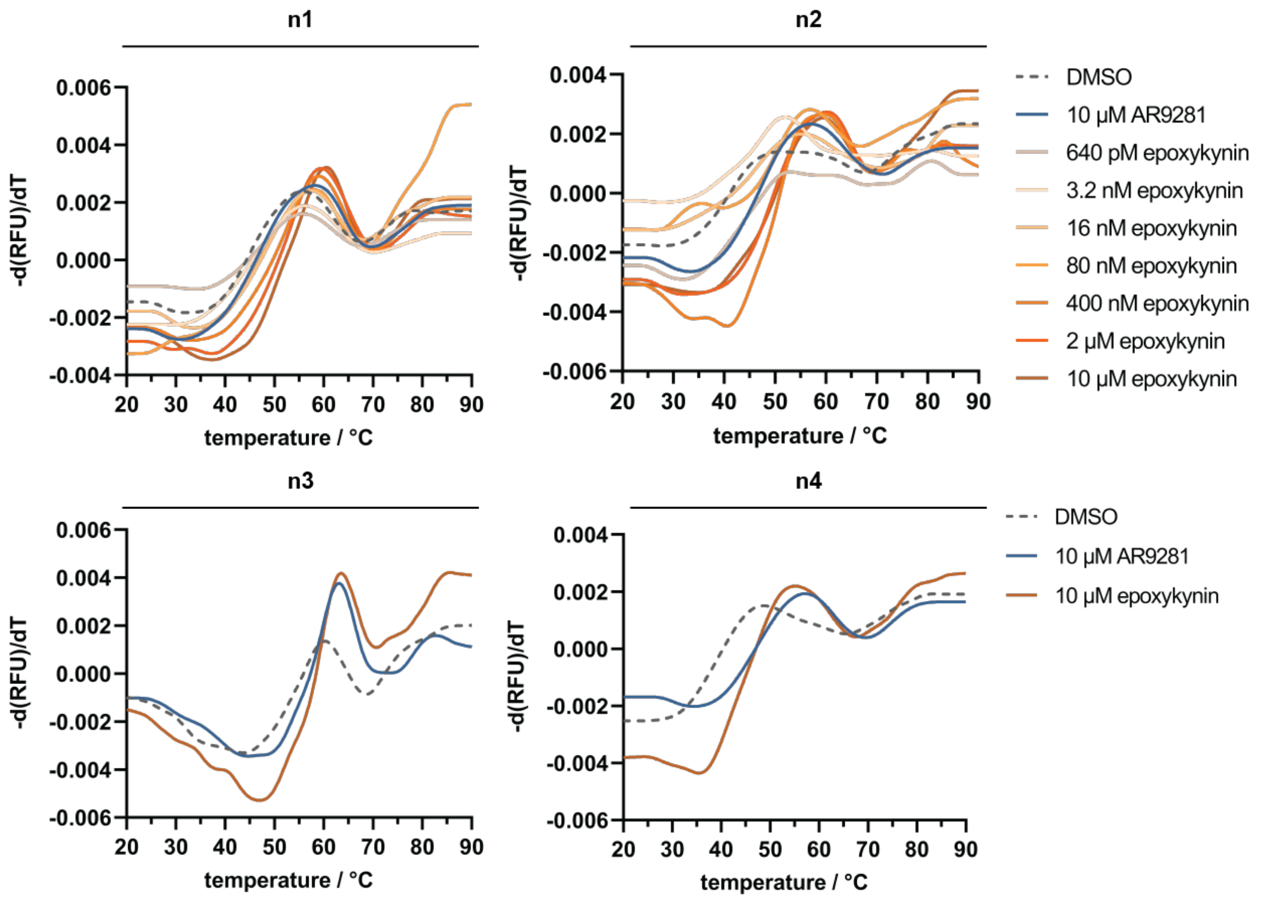

**Figure S4:** Dose-dependent $T_m$ shift of sEH upon binding of epoxykynin as detected using nanoDSF. Purified sEH was treated with epoxykynin or DMSO for 10 min at room temperature prior to detection of the intrinsic tryptophan/tyrosine fluorescence upon thermal denaturation. Related to Figure 3A.

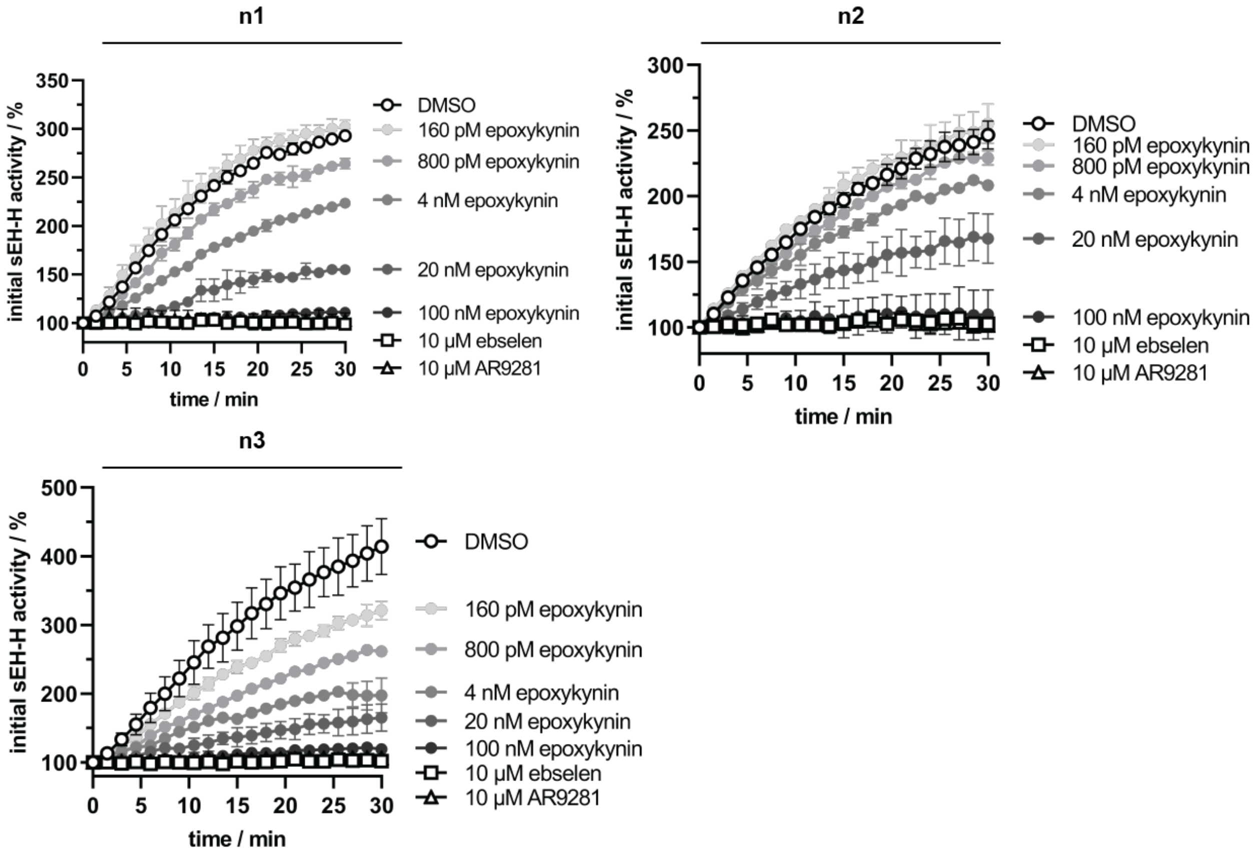

**Figure S5:** Dose-dependent inhibition of sEH-H by epoxykynin. The epoxide hydrolase activity of purified sEH (sEH-H) was measured by means of the conversion of the fluorogenic sEH-H substrate PHOME upon treatment with epoxykynin, 10 µM ebselen and 10 µM AR9281. Related to Figure 3B.

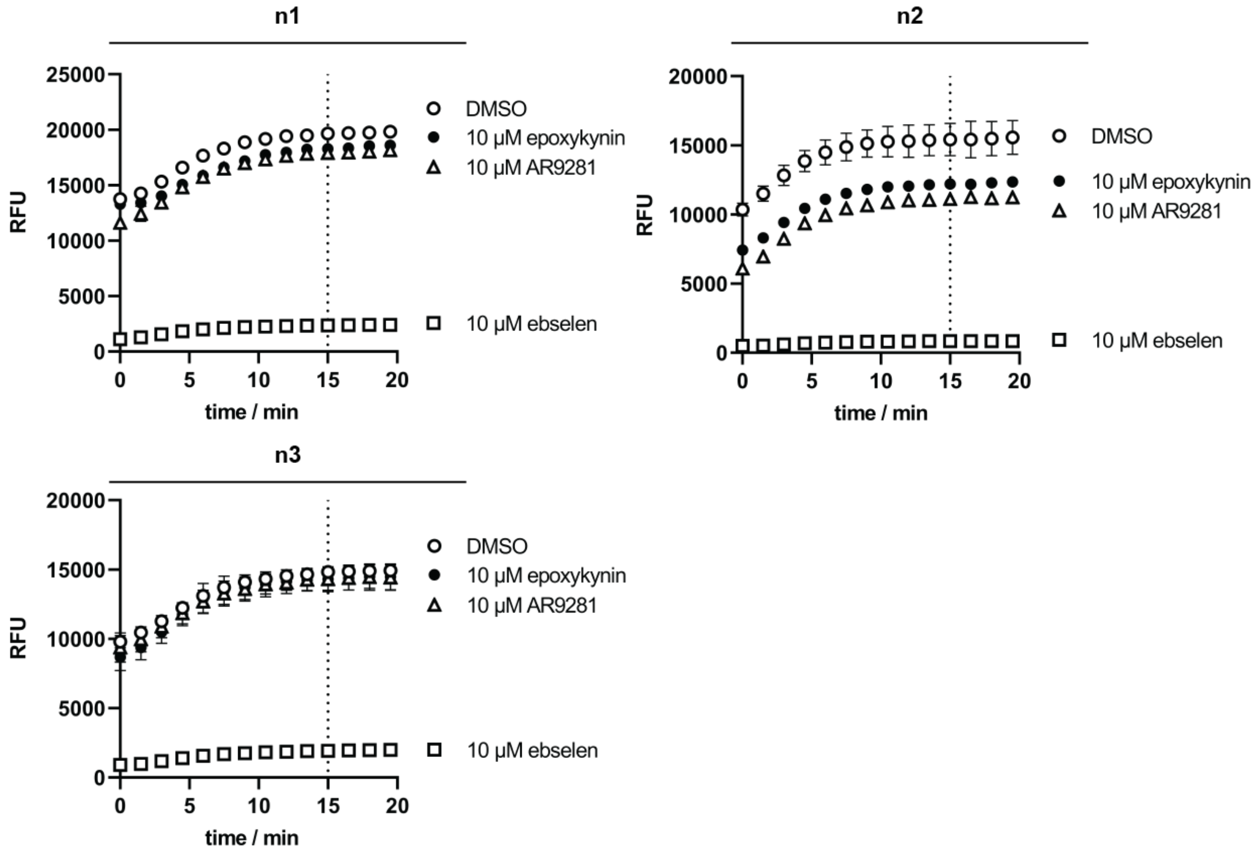

**Figure S6:** The phosphatase activity of purified sEH (sEH-P) was measured by means of an AttoPhos-based assay upon treatment with epoxykynin or AR9281 and ebselen as controls. Related to Figure 3C.

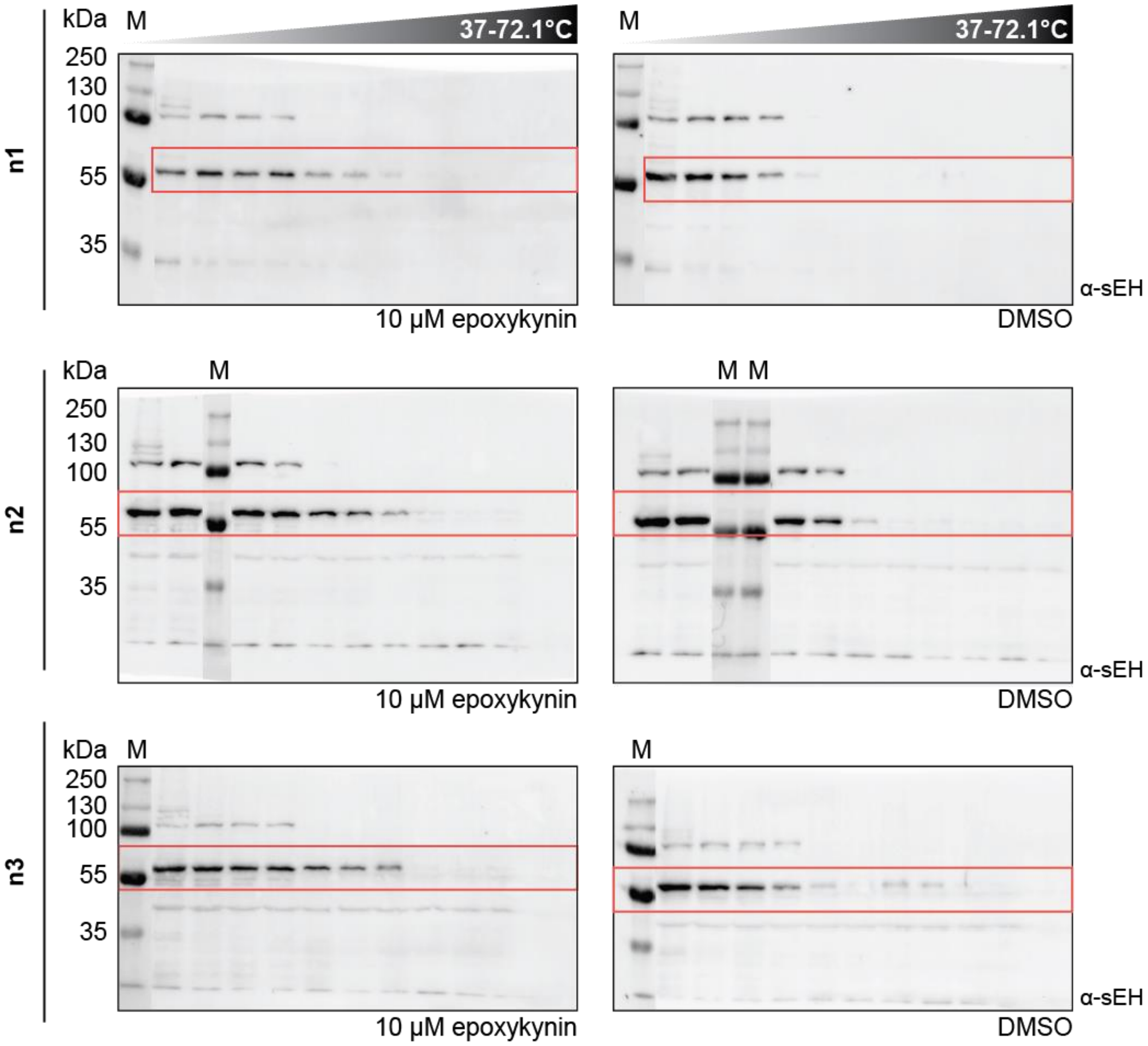

**Figure S7:** Cellular thermal shift assay (CETSA) for sEH in Jurkat cells. Cells were treated with 10 µM epoxykynin or DMSO for 15 min prior to heat treatment and cell lysis. Soluble proteins were analyzed using immunoblotting. sEH bands were acquired using an IRDye800CW secondary antibody, the marker (M) signal was acquired in the IRDye680RD channel. Related to Figure 3D.

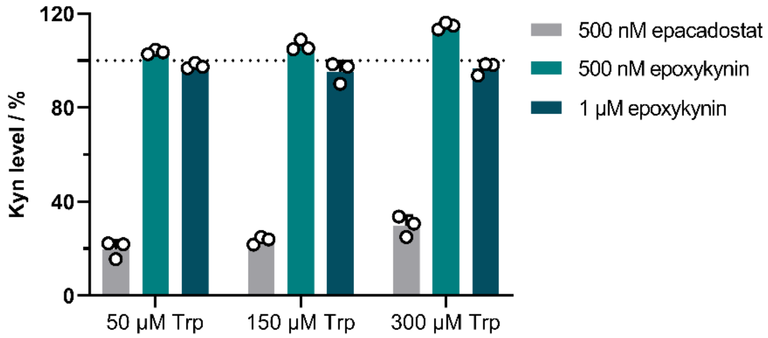

**Figure S8:** Kyn assay in HAP1 wt cells. HAP1 cells were treated with epacadostat or epoxykynin, Trp and IFN-γ for 48 h. Kyn levels were quantified using *p*-DMAB (mean values ± SD, n=3).

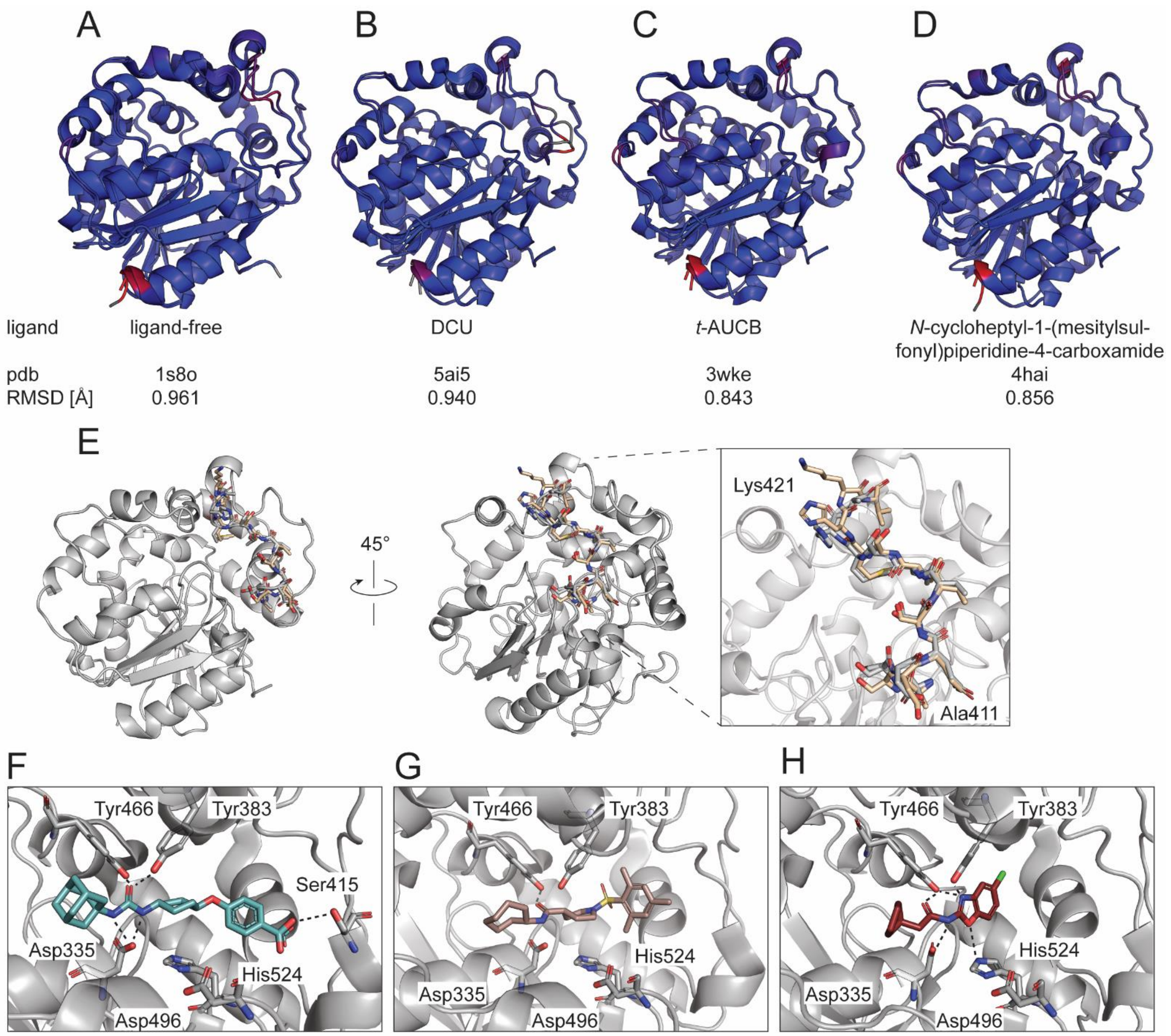

**Figure S9:** Alignments of co-crystal structures of epoxykynin bound to human sEH-H (pdb 8qzd) with ligand-free sEH-H (A, pdb 1s8o), *N*,*N*'-dicyclourea- (B, DCU, pdb 5ai5), *trans*-4-(4-(3-adamantan-1-ylureido)cyclohexyloxy)benzoic acid- (C, *t*-AUCB, pdb 3wke) and *N*-cycloheptyl-1-(mesitylsulfonyl)piperidine-4-carboxamide-bound sEH-H (D, pdb 4hai). Alignments were performed by superposition of the protein structures. The average distance between the atoms is represented by the root mean square deviation (RMSD). Blue color represents good alignment; red color represents high deviation; grey color represents unaligned residues. E) Overlay of co-crystal structures of epoxykynin (grey, pdb 8qzd) and *t*-AUCB (wheat, pdb 3wke) bound to human sEH-H. Residues Ala411 to Lys421 of the cap domain of sEH-H are shown as sticks. The amino acids in the active site are labeled with the three-letter code. Heteroatoms of the amino acid side chains are depicted in red (oxygen), blue (nitrogen) and yellow (sulfur). F-H) Crystal structures of *t*-AUCB (F, teal sticks, pdb wke), *N*-cycloheptyl-1-(mesitylsulfonyl)piperidine-4-carboxamide (G, brown sticks, pdb 4hai) and *N*-(5-chloro-1,3-benzoxazol-2-yl)-2-cyclopentylacetamide (H, red sticks, pdb 3pdc) bound to human sEH. Polar interactions are indicated by the dotted black lines. The amino acids in the active site are labeled with the three-letter code. Heteroatoms of the ligands and amino acid side chains are depicted in red (oxygen), blue (nitrogen), yellow (sulfur) and green (chlorine). Amino acids 497-500 are omitted for clarity.

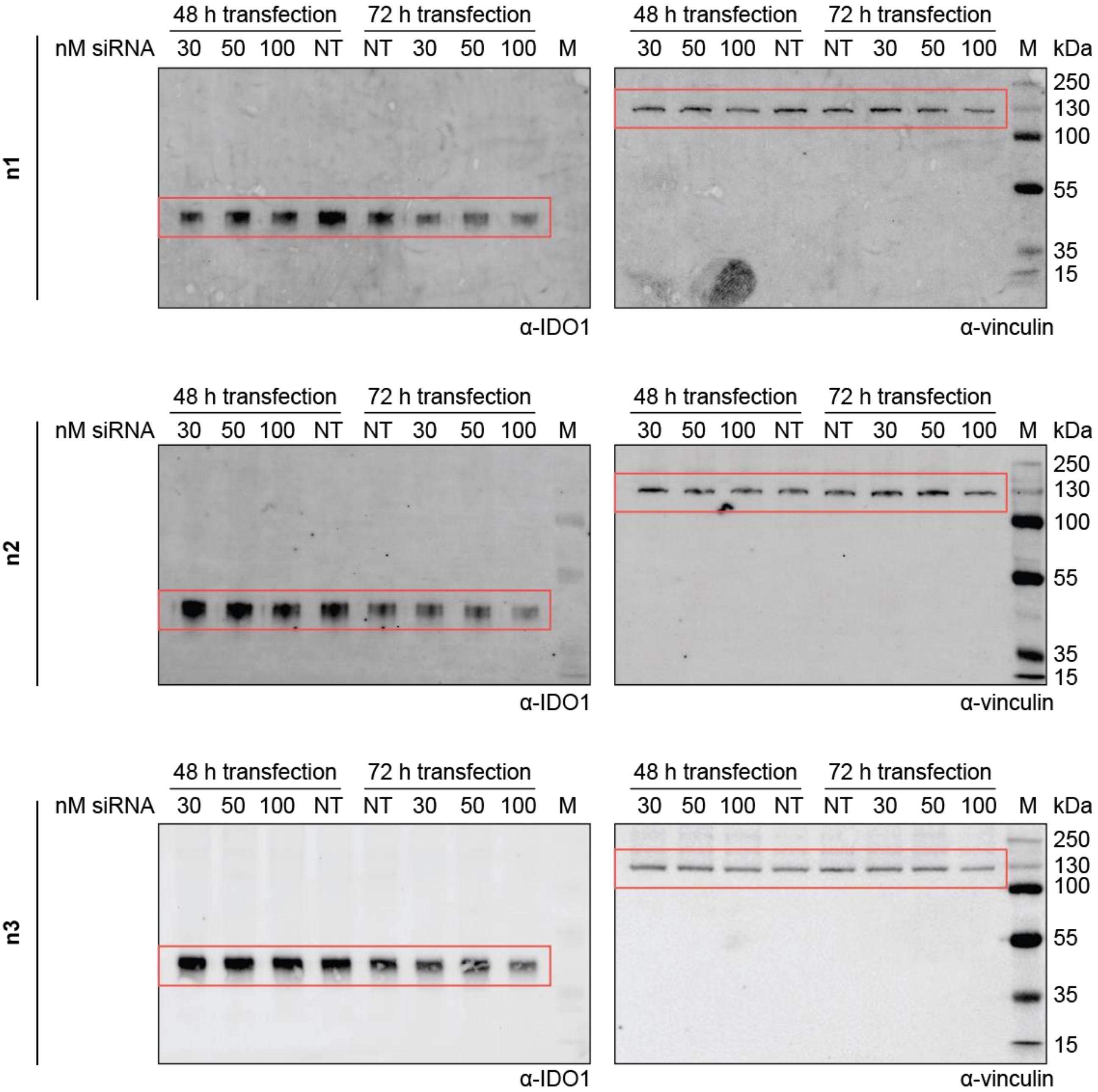

**Figure S10:** Knockdown (KD) of sEH resulted in decreased IDO1 levels. HeLa cells were transfected with non-targeting (NT) or sEH siRNA for 48-72 h and treated with IFN-γ for 48 h prior to quantification of protein levels *via* immunoblotting. IDO1 protein bands were acquired in the IRDye800CW channel, the vinculin control and marker (M) bands were acquired in the IRDye680RD channel. Related to figure 4A.

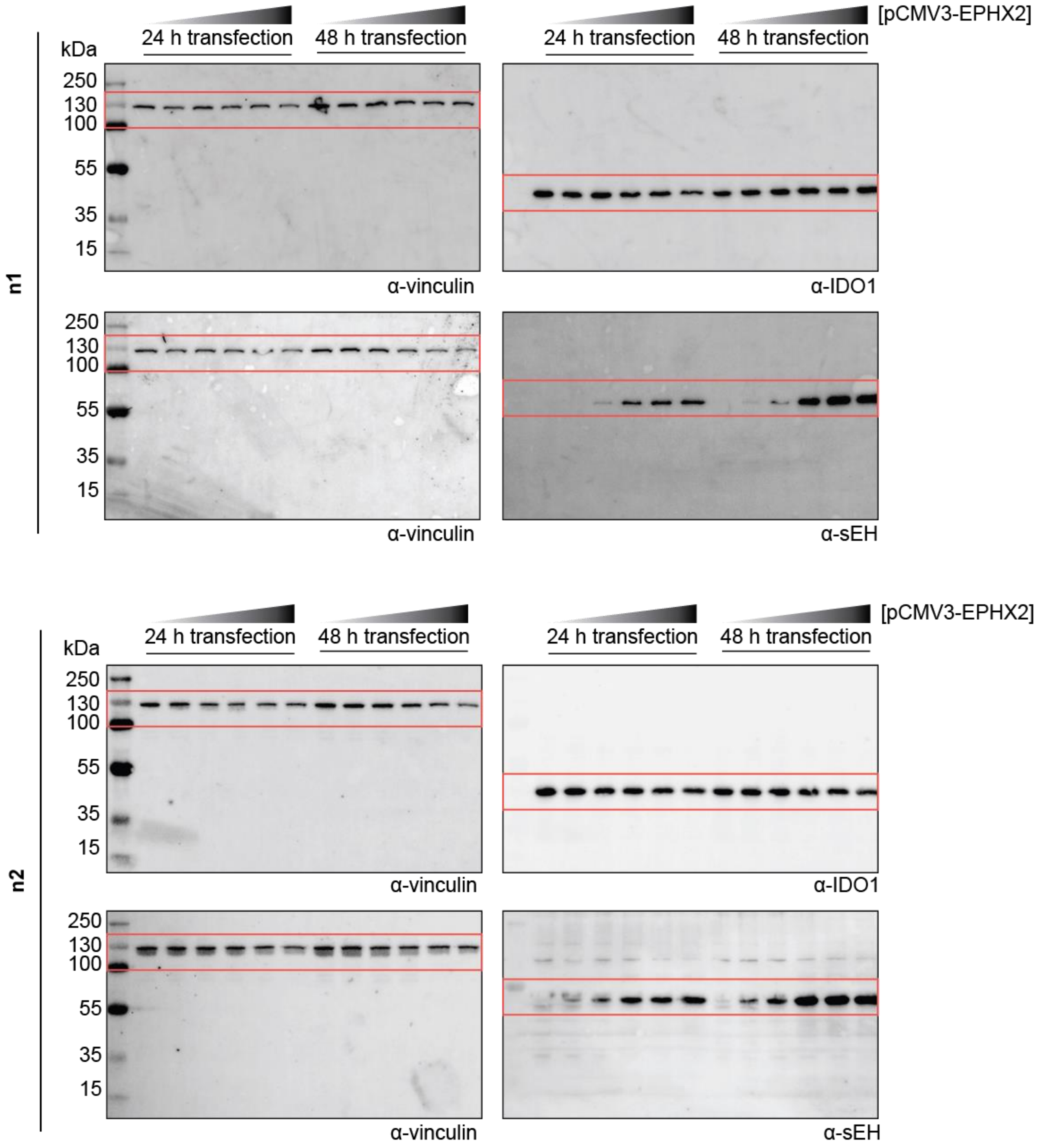

**Figure S11:** Overexpression of sEH resulted in increased IDO1 levels. HeLa cells were transfected with empty vector or pCMV3-EPHX2 for 24-48 h and treated with IFN-γ for 48 h prior to quantification of protein levels *via* immunoblotting. IDO1 and sEH protein bands were acquired in the IRDye800CW channel, the vinculin control and marker bands were acquired in the IRDye680RD channel. Related to figure 4B.

#### 2. Methods

2.1 Chemical Synthesis

**General Information**

The screening library of in total 157,332 compounds was composed of proprietary (10%) and commercially available compounds (90%). The commercial compounds included reference small molecules and natural products with annotated targets as well as compounds with drug-likeness and accordance to Lipinski's rule-of-five. Epoxykynin derivatives were purchased from ChemDiv, US.

Unless otherwise stated, all commercially available compounds were used as provided without further purification. Solvents for chromatography were technical grade.

$^{1}$H-NMR, $^{13}$C-NMR and $^{19}$F-NMR were recorded on Bruker DRX400 (400 MHz), Bruker DRX500 (500 MHz), INOVA500 (500 MHz) and Bruker DRX700 using $CD_2Cl_2$ as solvent and trimethoxybenzene as internal standard. Data is reported in the following order: chemical shift values (δ) are reported in ppm with the solvent resonance as internal standard ($CD_2Cl_2$: δ = 5.32 ppm for $^{1}$H, δ = 54.00 ppm for $^{13}$C). Multiplicities are indicated as br s: broad singlet, s: singlet, d: doublet, t: triplet, q: quartet, m: multiplet. Coupling constants (J) are given in Hertz.

HRMS spectra were acquired using the LTQ Fleet (Thermo Fisher Scientific, US), Ultimate 3000 HPLC (Thermo Fisher Scientific, US) and Xcalibur™ software (Thermo Fisher Scientific, US).

| time / min | %A | %B |
|---|---|---|
| 0 | 90 | 10 |
| 0.5 | 90 | 10 |
| 7.5 | 5 | 95 |
| 9.0 | 5 | 95 |
| 11 | 90 | 10 |

A: dd$H_2O$ + 0.1% (v/v) formic acid; B: acetonitrile + 0.1% (v/v) formic acid; flow rate:0.4 mL/min.

**Synthesis of the Affinity Pulldown Probes**

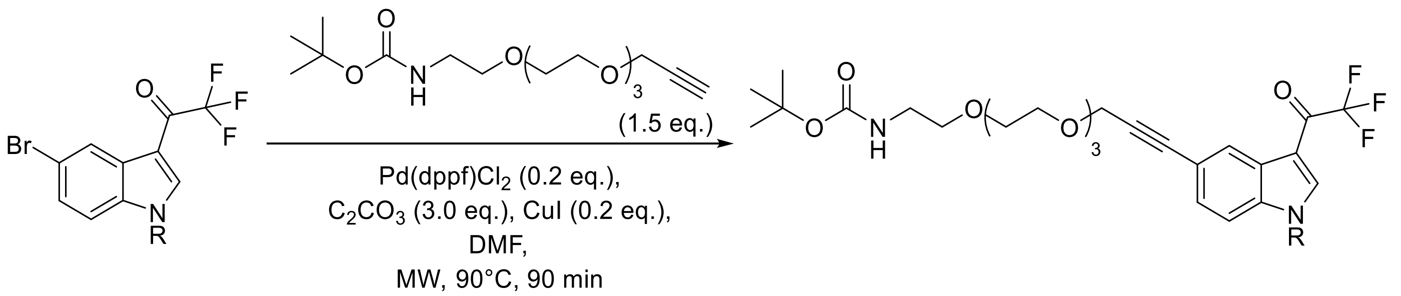

Into a reaction vessel were loaded the indole (1.0 eq.), the Pd(dppf)$Cl_2$ catalyst (0.2 eq.), $C_2CO_3$ (3.0 eq.) and CuI (0.2 eq.). The vessel was back-filled thrice with argon. 40 mL anhydrous DMF (0.05 M) was then added. Again, the vessel was back-filled thrice with argon. The mixture was stirred prior to addition of previously degassed amine-alkyne-PEG4 linker (1.5 eq.) in minimal amount of DMF. The reaction mixture was heated to 90°C for 90 min in a microwave reactor.

The mixture was poured into 20 mL 1 N HCl, extracted with 50 mL ethyl acetate and washed with brine. The organic phase was separated from the aqueous phase and washed with 20 mL brine and dried over $Na_2SO_4$ and concentrated *in vacuo*. The crude material was purified by silica gel column chromatography (Pent/EtOAc (1:1 to 1:2)) to yield a brown solid.

*tert*-butyl (15-(1-(2-(cycloheptylamino)-2-oxoethyl)-3-(2,2,2-trifluoroacetyl)-1*H*-indol-5-yl)-3,6,9,12-tetraoxa-pentadec-14-yn-1-yl)carbamate (**2a**)

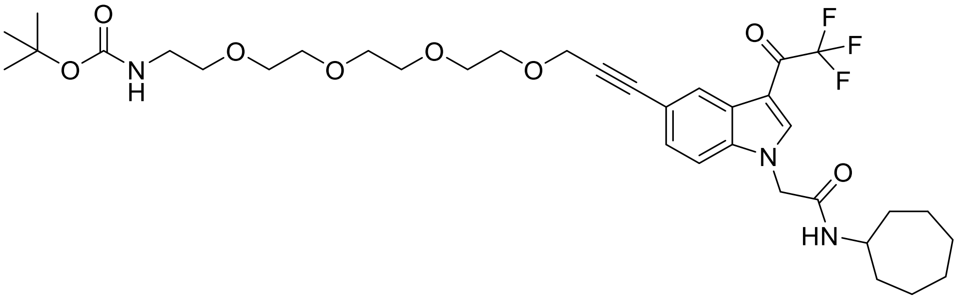

Colorless solid, 44.80% yield. **$^1$H-NMR (600 MHz – $CD_2Cl_2$):** δ 8.37 (d, *J* = 1.4 Hz, 1H), 7.95 (q, *J* = 1.7 Hz, 1H), 7.35 (dd, *J* = 8.5, 1.6 Hz, 1H), 7.24 (d, *J* = 8.5 Hz, 1H), 5.83 (bs, 1H, NH), 5.01 (bs, 1H, NH), 4.72 (s, 2H), 4.35 (s, 2H), 3.84 (tt, *J* = 8.5, 4.3 Hz, 1H), 3.69 – 3.66 (m, 2H), 3.62 – 3.59 (m, 2H), 3.57 – 3.48 (m, 8H), 3.41 (t, *J* = 5.2 Hz, 2H), 3.17 (q, *J* = 5.2 Hz, 2H), 1.77 (ddd, *J* = 12.8, 7.1, 3.4 Hz, 2H), 1.48 (p, *J* = 7.2, 6.6 Hz, 5H), 1.37 (d, *J* = 7.5 Hz, 4H), 1.34 (s, 9H), 1.32 – 1.27 (m, 1H); **$^{13}$C NMR (126 MHz – $CD_2Cl_2$):** δ 174.86 (d, *J* = 35.2 Hz), 163.96, 156.03, 139.57 (q, *J* = 4.8 Hz), 136.66, 128.57, 126.74, 126.19, 118.62, 117.97, 116.04, 110.73, 110.08, 86.31, 84.98, 78.96, 70.67 (2C), 70.58, 70.51, 70.41, 70.24, 69.39, 59.16, 51.33, 50.66, 34.91 (2C), 28.27(2C), 27.99 (3C), 24.13 (2C); **$^{19}$F NMR (565 MHz – $CD_2Cl_2$):** δ -71.66 (d, *J* = 3.0 Hz). **HRMS**: calculated for $[M+H]^+$, $C_{35}H_{49}F_3N_3O_8^+$ = 696.3466, found: 696.3464.

*tert*-butyl (15-(1-benzyl-3-(2,2,2-trifluoroacetyl)-1*H*-indol-5-yl)-3,6,9,12-tetraoxapentadec-14-yn-1-yl)carbamate (**3a**)

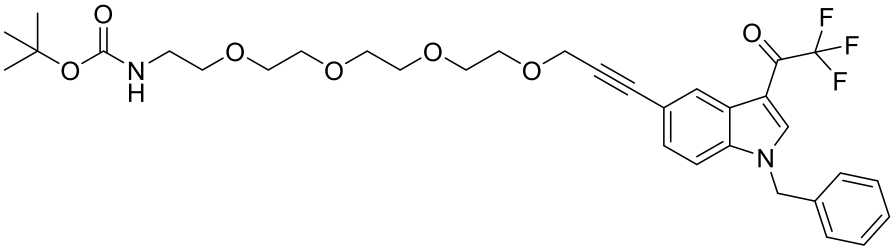

Brown solid, 18.28% yield. **$^1$H-NMR (500 MHz – $CD_2Cl_2$):** δ 8.49 (dd, *J* = 1.7, 0.7 Hz, 1H), 8.04 (d, *J* = 1.8 Hz, 1H), 7.41 (dd, *J* = 8.5, 1.6 Hz, 1H), 7.39 – 7.31 (m, 4H), 7.21 – 7.17 (m, 2H), 5.42 (s, 2H), 4.43 (s, 2H), 3.76 – 3.72 (m, 2H), 3.68 – 3.65 (m, 2H), 3.64 – 3.55 (m, 8H), 3.49 (t, *J* = 5.2 Hz, 2H), 3.26 (m, 2H), 1.41 (s, 9H). **$^{13}$C-NMR (176 MHz – $CD_2Cl_2$):** δ 175.06, 156.41, 139.31 – 139.07 (m), 137.05, 135.32, 129.72 (2C), 129.12, 128.73, 127.64 (2C), 127.60, 126.62, 118.91, 118.33, 111.70, 110.14, 86.78, 85.32, 79.32, 71.08 (2C), 71.01, 70.93, 70.82, 70.72, 69.76, 59.58, 52.10, 40.99, 28.68 (3C). **$^{19}$F-NMR (470 MHz – $CD_2Cl_2$):** δ -72.87 (d, *J* = 1.6 Hz). **HRMS**: calculated for $[M+H]^+$, $C_{33}H_{40}F_3N_2O_7^+$ = 632.2709, found: 632.6804.

**Deprotection of the Affinity Pulldown Probes**

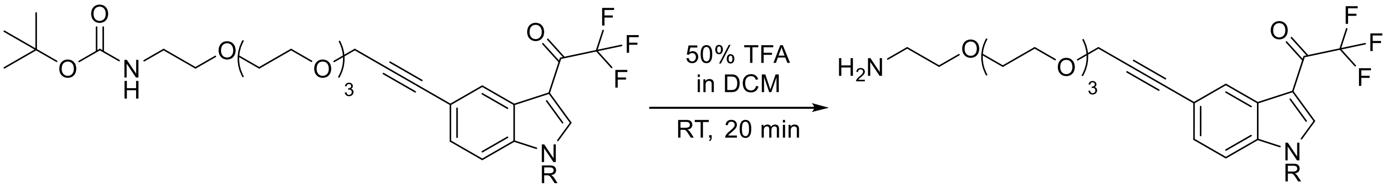

The boc-protected starting material was dissolved in 50% (v/v) TFA in DCM and stirred for 20 min at room temperature. The product was concentrated thrice with toluene *in vacuo* and lyophilized to afford the free amine probe.

2-(5-(3-(2-(2-aminoethoxy)ethoxy)prop-1-yn-1-yl)-3-(2,2,2-trifluoroacetyl)-1*H*-indol-1-yl)-N-cycloheptyl-acetamide (**2b**)

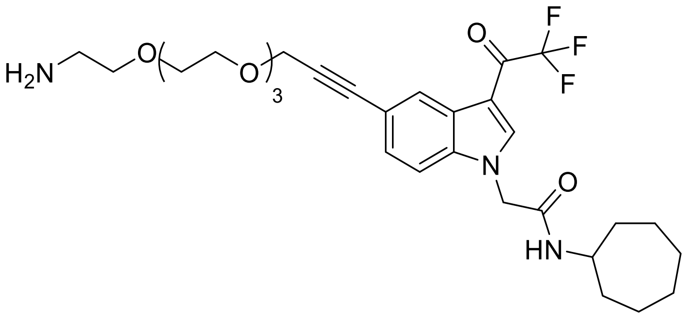

Brown solid, 98.12% yield. **HRMS**: calculated for $[M+H]^+$, $C_{30}H_{41}F_3N_3O_6^+$ = 696.29, found: 696.26; ; calculated for $[2M+H]^+$, $C_{60}H_{81}F_6N_6O_{12}^+$ = 1191.58, found: 1090.52.

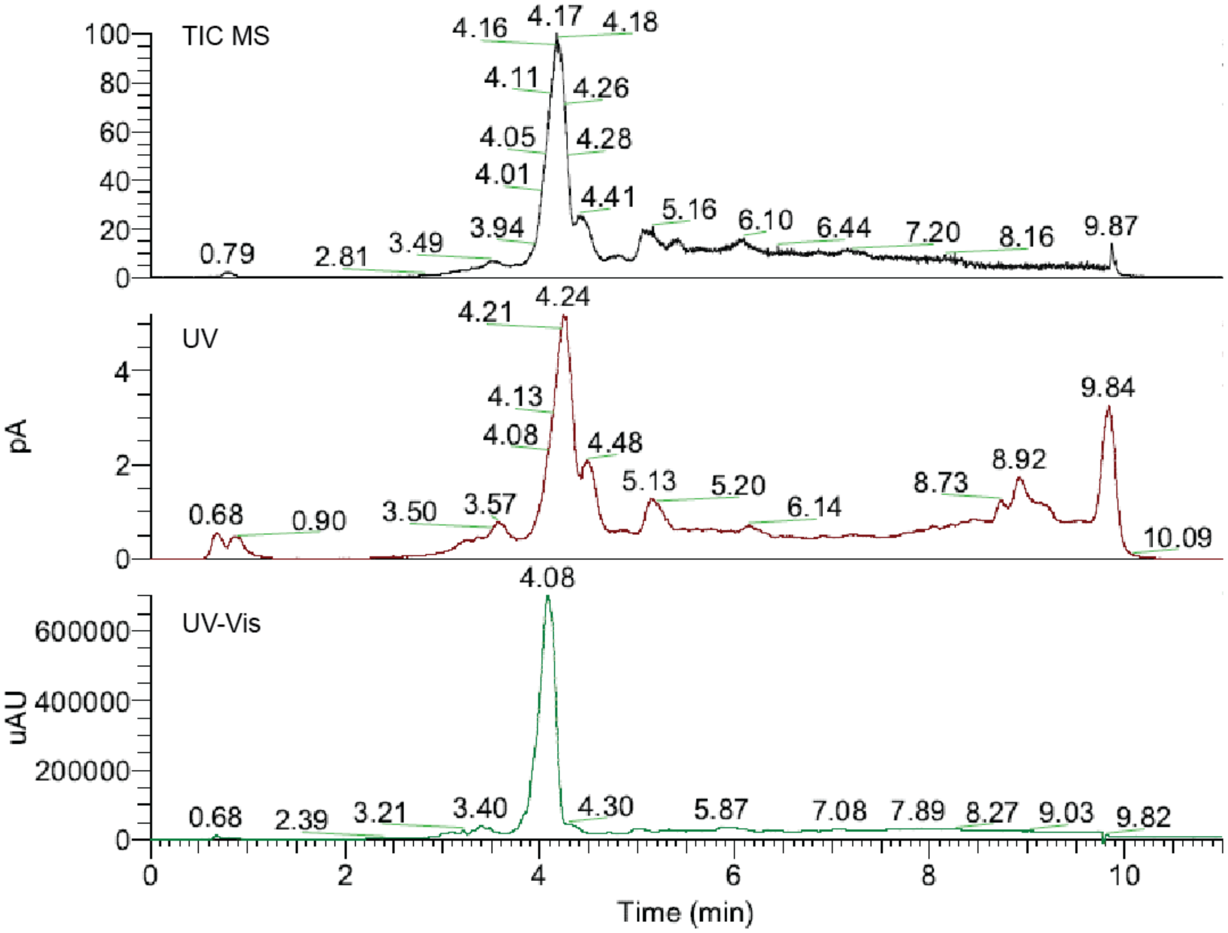
TIC MS
UV
UV-Vis
pA
uAU
Time (min)
4.16 4.17 4.18 4.11 4.26 4.05 4.28 4.01 4.41 3.94 5.16 6.10 6.44 7.20 8.16 9.87 0.79 2.81 3.49
4.21 4.24 4.13 4.08 4.48 9.84 8.92 8.73 5.13 5.20 6.14 0.68 0.90 3.50 3.57 10.09
4.08 0.68 2.39 3.21 3.40 4.30 5.87 7.08 7.89 8.27 9.03 9.82

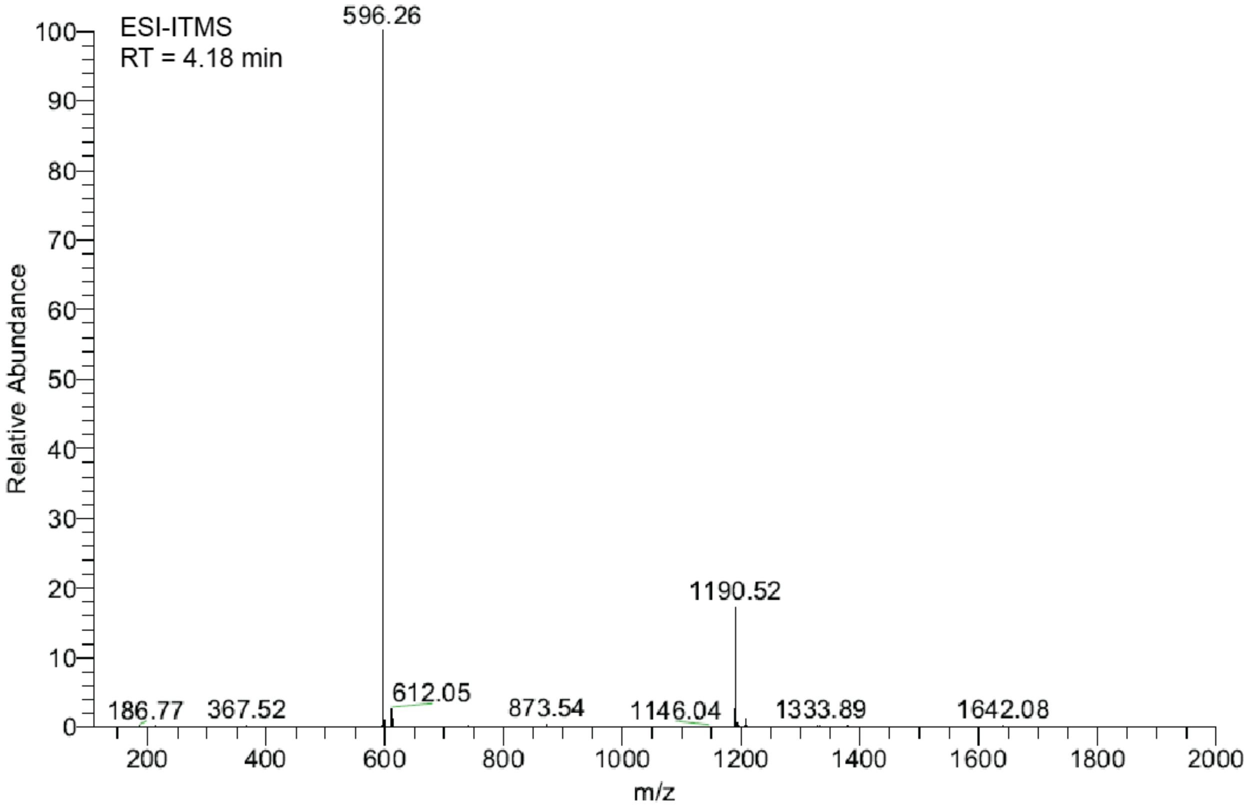
ESI-ITMS
RT = 4.18 min
Relative Abundance
m/z
596.26
1190.52
186.77 367.52 612.05 873.54 1146.04 1333.89 1642.08

1-(5-(3-(2-(2-aminoethoxy)ethoxy)prop-1-yn-1-yl)-1-benzyl-1*H*-indol-3-yl)-2,2,2-trifluoroethan-1-one (**3b**)

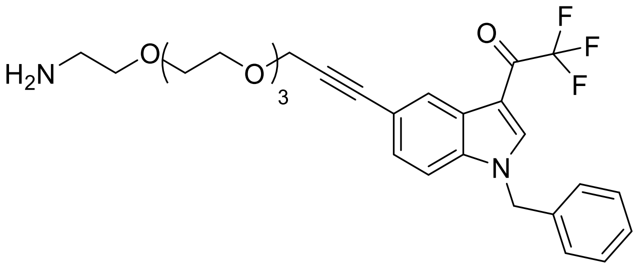

Brown solid, 99.79% yield. **HRMS**: calculated for [M+H]$^+$, $C_{28}H_{32}F_3N_2O_5^+$ = 533.23, found: 533.20; calculated for [2M+H]$^+$, $C_{56}H_{63}F_6N_4O_{10}^+$ = 1065.44, found: 1064.42.

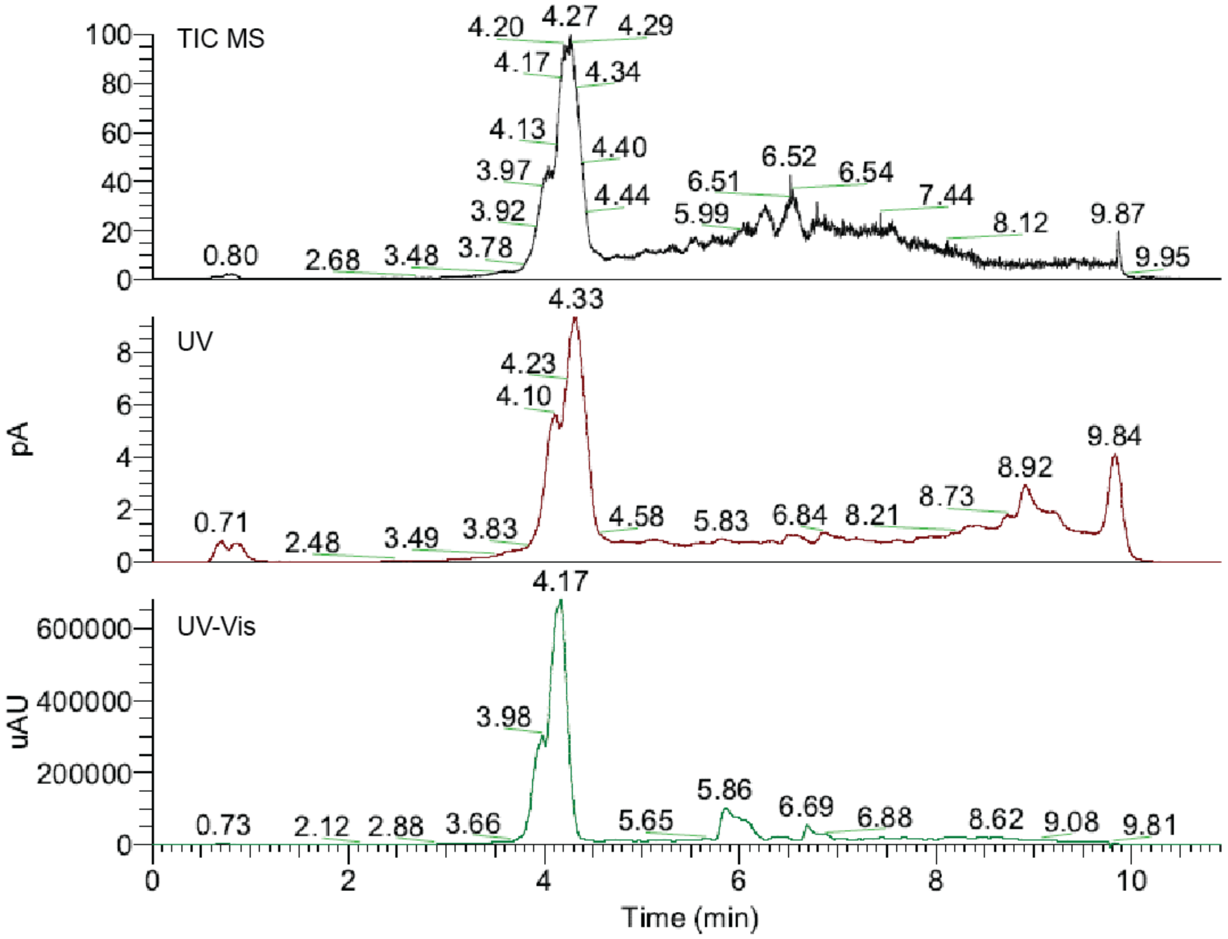

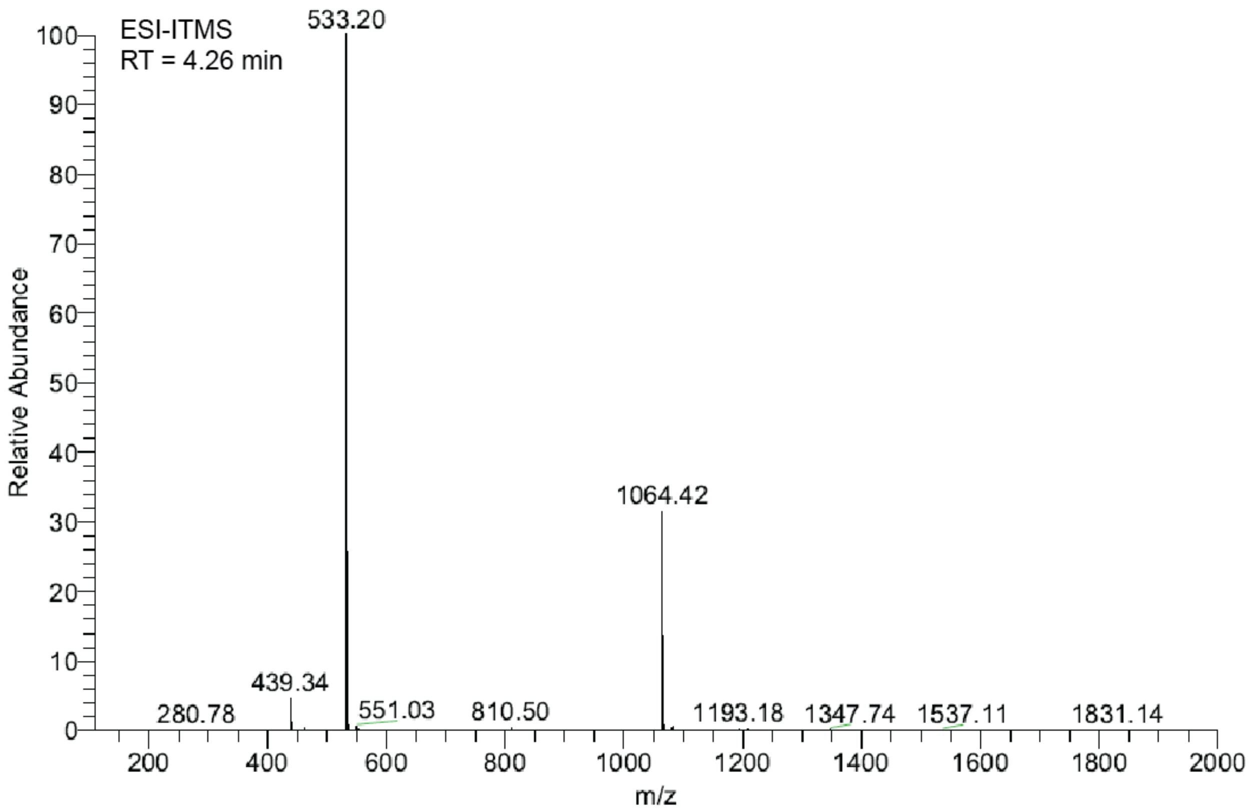
ESI-ITMS
RT = 4.26 min
533.20
1064.42
439.34
280.78
551.03
810.50
1193.18
1347.74
1537.11
1831.14
Relative Abundance
m/z

2.2 Biological Methods

**Key Resource Table**

| **REAGENT or RESOURCE** | **SOURCE** | **IDENTIFIER** |
|---|---|---|
| **Antibodies** | | |
| rabbit anti-IDO1 | Abcam | RRID:AB_2936946,<br>cat.# ab211017 |
| rabbit anti-sEH | ABclonal | RRID:AB_2763918,<br>cat.# A1885 |
| mouse anti-vinculin | Merck | RRID:AB_477629,<br>cat.# V9131 |
| **Chemicals, reagents and recombinant proteins** | | |
| AR-9281 | Hycultec | HY-111151 |
| acetic acid, glacial | Fisher Scientific | 10171460 |
| ascorbic acid | Sigma-Aldrich | A92902 |
| AttoPhos reagent | Promega | S1011 |
| catalase from bovine liver | Sigma-Aldrich | C9322 |
| 2-chloroacetamide | Sigma-Aldrich | C0267 |
| cOmplete™. EDTA-free protease inhibitor cocktail | Roche | 11873580001 |
| ebselen | Hycultec | HY-13750 |
| epacadostat | Selleckchem | S7910 |
| fetal bovine serum | Gibco | 10099-141 |
| IFN-γ | PeproTech | 300-02 |
| Intercept® (PBS) Blocking Buffer | LI-COR Biosciences | 927-70001 |

| REAGENT or RESOURCE | SOURCE | IDENTIFIER |
|---|---|---|
| Lipofectamine™ 2000 Transfection Reagent | Invitrogen Life Technologies | 11668-027 |
| NHS Mag Sepharose | Cytiva | 28-9440-09 |
| *para*-dimethylaminobenzaldehyde | Merck | 103058 |
| PhosphoStop | Roche | 4906845001 |
| trichloroacetic acid (TCA) | Sigma-Aldrich | T0699 |
| trifluoroacetic acid (TFA) | Thermo Fisher Scientific | 85183 |
| triethylammonium bicarbonate (TEAB) | Sigma-Aldrich | T7408 |
| trypsin, recombinant | Roche | 03708969001 |
| **Critical commercial assays** | | |
| *DC* Protein Assay | Bio-Rad Laboratories | 500-0116 |
| Dual-Glo® Luciferase Assay System | Promega | E2980 |
| Indoleamine 2,3-Dioxygenase 1 (IDO1) Activity Assay Kit | BioVision | K972-100 |
| MycoAltert™ Mycoplasma Detection Kit | Lonza | LT07-318 |
| QuantiTect® Reverse Transcription Kit | Qiagen | 205311 |
| RNase-Free DNase Set | Qiagen | 79254 |
| RNeasy® Mini Kit | Qiagen | 74106 |
| Soluble Epoxide Hydrolase Inhibitor Screening Assay Kit | Cayman | 10011671 |

| REAGENT or RESOURCE | SOURCE | IDENTIFIER |
|---|---|---|
| SsoAdvanced™ Universal SYBR® Green Supermix | Bio-Rad Laboratories | 1725274 |
| Deposited data | | |
| Co-crystal structure of epoxykynin bound to sEH-H | PDB | 8qzd |
| **Experimental models, cell lines** | | |
| BxPC-3 | DSMZ | RRID:CVCL_0186, cat.# 760 |
| HAP1 | Horizon Discovery | RRID:CVCL_Y019, cat.# c631 |
| HCT116 | DSMZ | RRID:CVCL_0291, cat.# 581 |
| HCT116 MTAP$^{(-/-)}$ | Horizon Discovery | cat.# HD R02-033 |
| HeLa | DSMZ | RRID: CVCL_0030, cat.# 57 |
| HEK293T | ATCC | RRID:CVCL_1926, cat.# 11268 |
| Jurkat | DSMZ | RID:CVCL_0065, cat.# 282 |
| **Recombinant DNA** | | |
| pCMV3-EPHX2 | Sino Biological | HG14856-UT |
| pCMV3-IDO1 | Sino Biological | HG11650-UT |
| pGEX6p-GST-IDO1 | kindly provided by A. De Antoni[2] | |
| pRL-TK | Promega | E2241 |

| REAGENT or RESOURCE | SOURCE | IDENTIFIER |
|---|---|---|
| pRSF-EPHX2 | kindly provided by E. Proschak[3] | |
| pXPG-IDO1 | kindly provided by G.M. Doody[4] | |
| **Software and algorithms** | | |
| Bio-Rad CFX96 Real-Time PCR Detection System | Bio-Rad | RRID:SCR_018064 |
| CCP4 suite | (Agirre *et al.*, 2023)[5] | RRID:SCR_007255 |
| ChemDraw v18.2 | PerkinElmer | RRID:SCR_016768 |
| Coot | (Emsley *et al.*, 2010)[6] | RRID:SCR_014222 |
| Fiji ImageJ | (Schindelin *et al*., 2012)[7] | RRID: SCR_002285 |
| Image Lab | Bio-Rad | RRID:SCR_014210 |
| Image Studio v5.2 | LI-COR | RRID:SCR_015795 |
| MaxQuant v1.6.3.4 | (Cox and Mann, 2008)[8] | RRID:SCR_014485 |
| Perseus v1.6.2.2 | (Tyanova and Cox, 2018)[9] | RRID:SCR_015753 |
| Prism v9.2.0 | GraphPad | RRID:SCR_002798 |
| XDS | (Kabsch, 2010)[10] | RRID:SCR_015652 |

#### Experimental Procedures

##### Cell culture

Human cell lines were maintained at 37°C and 5% $CO_2$ in a humidified atmosphere and sub-cultivated twice a week. All cell lines were tested regularly for mycoplasma contamination and were always free of mycoplasma. BxPC-3 cells (DSMZ#760, female, RRID:CVCL_0186) and Jurkat cells (DSMZ#282, male, RID:CVCL_0065) were grown in RPMI-1640 medium supplemented with 10% FBS, 2 mM L-glutamine, 1 mM sodium pyruvate, 4.5 g/L glucose, 10 mM HEPES and 1.5 g/L $NaHCO_3$. HeLa (DSMZ#57, female, RRID: CVCL_0030), HCT116 (DSMZ#581, male, RRID:CVCL_0291), HCT116 MTAP$^{(-/-)}$ (Horizon Discovery#HD R02-033, male, parental RRID:CVCL_0291) and HEK293T cells (ATCC#11268, female, RRID:CVCL_1926) were cultivated in DMEM supplemented with 10% FBS, 4.5 g/L glucose, 4 mM L-glutamine, 1 mM sodium pyruvate, 1% non-essential amino acids, 3.7 g/L $NaHCO_3$. HAP1 cells (Horizon Discovery#c631, male, RRID:CVCL_Y019) were maintained in IMDM medium supplemented with 10% FBS, 4.5 g/L glucose, 4 mM L-glutamine, 25 mM HEPES and 3.0 g/L $NaHCO_3$.

##### Kynurenine (Kyn) assays

*High-throughput Kyn assay*

Automated screening for modulators of Kyn levels in BxPC-3 cells was performed as published earlier.[11] Briefly, BxPC-3 cells (1,000 cells/well) were seeded in black 1536-well plates prior to incubation for 24 h and subsequent treatment with compounds, 380 µM Trp and 50 ng/mL IFN-γ for 48 h. Afterwards, trichloroacetic acid was added to a final concentration of 7% (v/v), the plates were incubated for 10 min at 37°C and centrifuged for 10 min at 1620 x g. Kyn was detected by addition of 17.5 µM Kyn sensor[12] in assay buffer (excitation: 535 nm, emission: 595 nm). In total, 157,332 compounds were screened at a concentration of 7.1 µM. The screening library (157,332 compounds) comprised 90% commercial (e.g. LOPAC and Prestwick Chemical Libraries) and 10% in-house synthesized compounds. The epoxykynin compound class was purchased from ChemDiv, US. Compounds that reduced Kyn levels by ≥70% were subjected to $IC_{50}$ determination. Cytotoxic compounds were excluded by nuclear staining with Hoechst 33342 prior to incubation for 30 min at 37°C and imaging with the ImageXpress Micro XL (excitation: 377/50 nm, emission: 447/60 nm, Molecular Devices, US). Image analysis was performed by means of cell count with the Cell Proliferation HT Application Module of the MetaXpress software (Molecular Devices, US).

*Manual Kyn assay*

For manual testing, BxPC-3 or HeLa cells (20,000 cells/well or 5,000 cells/well, respectively) were seeded in 96-well plates in medium without phenol red. After 24 h, the Kyn pathway was induced by addition of 50 ng/mL IFN-γ and 380 µM (BxPC-3) or 164.15 µM (HeLa) Trp with simultaneous treatment with the compounds at the indicated concentrations followed by incubation for 48 h. Subsequently, trichloroacetic acid was added to a final concentration of 7% (v/v), the plates were incubated for 15 min at room temperature and centrifuged for 10 min at 1800 x g. For determination of Kyn levels with the Ehrlich reagent, an equal volume of freshly prepared 2% (w/v) *p*-DMAB in glacial acetic acid (Ehrlich reagent) was added prior to measuring the absorbance at 492 nm and 650 nm on the Spark® Multimode Microplate Reader (Tecan, AT). The background absorbance $A_{650}$ was subtracted from the absorbance $A_{492}$ of the Kyn-*p*-DMAB adduct and normalized to the DMSO control. Data analysis was performed using a non-linear regression curve fit and GraphPad Prism 9.0 (GraphPad Software, Inc, US) to generate dose-response curves and obtain $IC_{50}$ values.

*Determination of Kyn levels using LC-MS*

For determination of Kyn levels *via* LC-MS, BxPC-3 (20,000 cells/well) were seeded in 96-well plates in medium without phenol red. After 24 h, the Kyn pathway was induced by addition of 50 ng/mL IFN-γ and 380 µM Trp with simultaneous treatment with the compounds at the indicated concentrations followed by incubation for 48 h. Subsequently, trichloroacetic acid was added to a final concentration of 7% (v/v), the plates were incubated for 15 min at room temperature and centrifuged for 10 min at 1800 x g. Kyn and Trp was quantified by HPLC-MS/MS using the LTQ Velos Pro and Dionex HPLC (Thermo Fisher Scientific, US). Data analysis was performed with Xcalibur™ (Thermo Fisher Scientific, US) and represented using GraphPad Prism 9.0 (GraphPad Software, Inc, US).

*Kyn assay in IDO1-HEK293T cells*

HEK293T cells (25,000 cells/well) were reverse-transfected with pCMV3-IDO1 (1 µg/96-well plate, Sino Biological, CN) using 3.9 µL Lipofectamine™ 2000 (Invitrogen, US) prior to incubation for 20 h. Subsequently, cells were treated with 500 µM Trp and the compounds for 24 h. Thereafter, trichloroacetic acid was added to a final concentration of 7% (v/v), the plates were incubated for 15 min at room temperature and centrifuged for 10 min at 1800 x g. Afterwards, an equal volume of freshly prepared 2% (w/v) *p*-DMAB in glacial acetic acid (Ehrlich reagent) was added prior to measuring the absorbance at 492 nm and 650 nm on the Spark® Multimode Microplate Reader (Tecan, AT). To determine the Kyn levels, the background absorbance $A_{650}$ was subtracted from the absorbance $A_{492}$ of the Kyn-*p*-DMAB adduct and normalized to the DMSO control. Data analysis was performed using a non-linear regression curve fit and GraphPad Prism 9.0 (GraphPad Software, Inc, US) to generate dose-response curves and obtain $IC_{50}$ values.

*Kyn assay in IDO1-HCT116 cells*

HCT116 wt and HCT116 $MTAP^{(-/-)}$ cells (30,000 cells/well) were reverse-transfected with pCMV3-IDO1 (2 µg/96-well plate, Sino Biological, CN) using 4 µL Lipofectamine™ 3000 and 4 µL P3000™ reagent (Invitrogen, US) prior to incubation for 24 h. Subsequently, cells were treated with 500 µM Trp and the compounds for 36 h. Thereafter, trichloroacetic acid was added to a final concentration of 7% (v/v), the plates were incubated for 15 min at room temperature and centrifuged for 10 min at 1800 x g. Afterwards, an equal volume of freshly prepared 2% (w/v) *p*-DMAB in glacial acetic acid (Ehrlich reagent) was added prior to measuring the absorbance at 492 nm and 650 nm on the Spark® Multimode Microplate Reader (Tecan, AT). To determine the Kyn levels, the background absorbance $A_{650}$ was subtracted from the absorbance $A_{492}$ of the Kyn-*p*-DMAB adduct and normalized to the DMSO control. Data analysis was performed using a non-linear regression curve fit and GraphPad Prism 9.0 (GraphPad Software, Inc, US) to generate dose-response curves and obtain $IC_{50}$ values.

*Kyn assay in sEH-HAP1 cells*

HAP1 cells (20,000 cells/well) were reverse-transfected with pCMV3-EPHX2 (1 and 3 µg/96-well plate, Sino Biological, CN) using 2 µL Lipofectamine™ 3000 and 2 µL P3000™ reagent (Invitrogen, US) prior to incubation for 20 h. Subsequently, cells were treated with 10 ng/mL IFN-γ and 100 µM Trp and the compounds for 48 h. Thereafter, trichloroacetic acid was added to a final concentration of 7% (v/v), the plates were incubated for 15 min at room temperature and centrifuged for 10 min at 1800 x g. Afterwards, an equal volume of freshly prepared 2% (w/v) *p*-DMAB in glacial acetic acid (Ehrlich reagent) was added prior to measuring the absorbance at 492 nm and 650 nm on the Spark® Multimode Microplate Reader (Tecan, AT). To determine the Kyn levels, the background absorbance $A_{650}$ was subtracted from the absorbance $A_{492}$ of the Kyn-*p*-DMAB adduct and normalized to the DMSO control. Data analysis was performed using a non-linear regression curve fit and GraphPad Prism 9.0 (GraphPad Software, Inc, US) to generate dose-response curves and obtain $IC_{50}$ values.

*In vitro Kyn assay*

To detect direct inhibition, 1 µM recombinant human IDO1 was incubated with compounds for 40 min at 37°C in 50 mM potassium phosphate buffer (16.9 mM $K_2HPO_4$, 33.1 mM, $KH_2PO_4$, pH 6.5). Subsequently, 10 mM ascorbic acid, 10 µM methylene blue, 2 mM Trp and 100 µg/mL catalase were added and samples were incubated for 60 min at room temperature or 37°C. Trichloroacetic acid was added to a final concentration of 7% (v/v) and samples were incubated for 30 min at 70°C. Afterwards, an equal volume of freshly prepared 2% (w/v) *p*-DMAB in glacial acetic acid (Ehrlich reagent) was added prior to measuring absorbance at 492 nm and 650 nm on the Spark® Multimode Microplate Reader (Tecan, AT). To determine the Kyn levels, the background absorbance $A_{650}$ was subtracted from the absorbance $A_{492}$ of the Kyn-*p*-DMAB adduct and normalized to the DMSO control. Data analysis was performed using a non-linear regression curve fit and GraphPad Prism 9.0 (GraphPad Software, Inc, US) to generate dose-response curves and obtain $IC_{50}$ values.

*Kyn assay in lysates*

The Kyn assay in lysate was performed using the commercial Indoleamine 2,3-Dioxygenase 1 (IDO1) Activity Assay Kit (cat# K972-100, BioVision Inc, US). Therefore, BxPC-3 cells (1 x $10^6$ cells) were seeded in 60 mm-dishes and incubated for 24 h. On the next day, cells were treated with 50 ng/mL IFN-γ prior to incubation for 48 h. If the cells were pre-treated with compound or siRNA, the compounds or siRNA were added simultaneously with IFN-γ. The plates were washed twice with warm PBS prior to harvesting the cells with 750 µL trypsin/EDTA solution. The cells were washed with ice-cold PBS and collected by centrifugation at 4°C and 300 x g for 5 min prior to resuspension in 300 µL IDO1 assay buffer containing protease inhibitors. The cells were lysed by three consecutive freeze/thaw cycler.

For the Kyn detection, the 2X reaction premix was prepared by diluting the antioxidant mix 50-fold in IDO1 assay buffer. A 10X working solution of the IDO1 substrate Trp was prepared by 10-fold dilution of the stock in IDO1 assay buffer (final Trp concentration in the assay: 100 µM). In a final volume of 11 µL containing 5.5 µL 2X reaction premix, 7 µg of cell lysate was treated with the compounds prior to reaction by addition of 1.1 µL of 10X Trp. The plate was sealed with a sticky silver sealer and incubated for 45 min at 37°C and

300 rpm in the dark prior to addition of 5 µL of the fluorogenic developer solution. The plate was incubated for another 3 h at 45°C and 300 rpm in the dark. After cooling down for at least 1 h, the fluorescence was measured at an excitation wavelength of 402 nm and an emission wavelength of 488 nm on the Spark® Multimode Microplate Reader (Tecan, AT).

**Expression and purification of recombinant IDO1**

Recombinant human GST-IDO1 (rhIDO1) was expressed in *E. coli* BL21 DE3 cells using a pGEX6p-2rbs-GST-IDO1 and purified as published previously.[2, 13] Bacteria were transformed with the vector by a heat shock at 42°C for 60 s and cells derived from a single colony were used to inoculate LB medium containing 100 µg/mL ampicillin and 30 µg/mL chloramphenicol prior to incubation overnight at 37°C with rotation. On the next day, 2,500 mL LB medium were inoculated with 50 mL of the bacteria suspension and IDO1 protein expression was induced by addition of 200 µM isopropyl β-d-1-thiogalactopyranoside prior to incubation at 18°C overnight.

Afterwards, the cells were pelleted by centrifugation at 3600 x g and 4°C for 20 min and resuspended in buffer 1 (50 mM TRIS-HCl, pH 7.4, 100 mM NaCl) and homogenized by sonication, followed by mechanical cell lysis. The soluble fraction was separated from the cell debris by centrifugation at 13,000 x g and 10°C for 35 min and transferred to a GSTrap™ HP column (Cytiva, US). GST-IDO1 was washed on column with buffer 2 (50 mM TRIS-HCl, pH 7.4, 100 mM KCl), followed by on-column cleavage overnight with 3.2 mg/mL PreScission protease (Cytiva, US) in buffer 3 (50 mM TRIS-HCl, pH 7.0, 150 mM NaCl, 1 mM DTE, 200 µM hemin). The eluted protein was purified by size exclusion chromatography with a HiLoad™ 16/600 Superdex™ 200 pg column (Cytiva, US) and purity of the protein was confirmed by SDS-PAGE.

***IDO1* promoter reporter gene assay**

HEK293T cells (25,000 cells/well) were reverse-transfected with pXPG-IDO1[14] (4 µg/96-well plate, kindly provided by Gina M. Doody, Leeds, UK) and pRL-TK (300 ng/96-well plate, Promega, US) using 12.9 µL Lipofectamine™ 2000 (Invitrogen, USA) prior to incubation for 24 h. Afterwards, cells were treated with 50 ng/mL IFN-γ and the compounds for 48 h. Luminescence generated by the Fluc reporter and the control reporter Rluc was measured using the Dual-Glo ® Luciferase Assay System (Promega, US) on the Spark® Multimode Microplate Reader (Tecan, AT). To determine the *IDO1* promoter activity, Fluc signals were divided by the Rluc signals and normalized to the values of the DMSO control. Data analysis was performed using a non-linear regression curve fit and GraphPad Prism 9.0 (GraphPad Software, Inc, US) to generate dose-response curves and obtain $IC_{50}$ values.

**Trp uptake assay**

BxPC-3 cells (30,000 cells/well) were seeded in 96-well plates in Trp-free RPMI 1640 medium and incubated for 48 h prior to addition of 50 ng/mL IFN-γ. Trp starvation was continued for another 24 h. Afterwards, the medium was exchanged for Trp-free medium containing the control inhibitors 5 mM L-Leu (inhibitor of system-L amino acid transporters (LAT)), 1 mM 1-methyl-L-tryptophan (inhibitor of tryptophanyl-tRNA synthetase (TrpRS)) and compounds, and cells were incubated for 30 min. Subsequently, 50 µM Trp was added and samples were incubated for another 30 min. The supernatant was transferred to a new plate, trichloroacetic acid was added to a final concentration of 7% (v/v) and plates were incubated for 15 min at room temperature. Samples were centrifuged for 10 min at 1800 x g and Trp was quantified by HPLC-MS/MS using the LTQ Velos Pro and Dionex HPLC (Thermo Fisher Scientific, US). Data analysis was performed with Xcalibur™ (Thermo Fisher Scientific, US) and represented using GraphPad Prism 9.0 (GraphPad Software, Inc, US).

**RNA purification and RT-qPCR**

HeLa cells (250,000 cells/well) were seeded in 6-well plates and incubated for 24 h prior to addition of 50 ng/mL IFN-γ and compounds. After 24 h, RNA was extracted using the RNeasy Plus Mini Kit (QIAGEN, DE) following the manufacturer's procedure. DNA was removed on column by DNase digestion with RNase-free DNase Set (QIAGEN, DE) according to the manufacturer's instructions. Following the manufacturer's protocol, cDNA templates were synthesized from 800 ng total RNA using the QuantiTect Reverse Transcription Kit (QIAGEN, DE).

The expression levels of the *IDO1* gene and the reference *GAPDH* were assessed by reverse trancription quantitative PCR (qPCR). Therefore, 100 ng cDNA was amplified using 500 nM of gene-specific primers and SsoAdvanced Universal SYBR Green Supermix (Bio-Rad Laboratories, DE) in a total volume of 10 µL for 50 cycles using the CFX96 Touch™ Real-Time PCR Detection System (Bio-Rad Laboratories, DE). Relative *IDO1* and *EPHX2* expression levels were calculated using the $\Delta\Delta C_t$ method with *GAPDH* as reference gene[15] and represented using GraphPad Prism 9.0 (GraphPad Software, Inc, US).

The sequences of the primers for *IDO1* were 5'-GCCTGATCTCATAGAGTCTGGC-3' (forward) and 5'-TGCATCCCAGAACTAGACGTGC-3' (reverse). The sequences of the primers for *EPHX2* were 5'-CCTTCATACCAGCAAATCCCAACA-3' (forward) and 5'-TTCAGCCTCAGCCACTCCT-3' (reverse).[16] The sequences of the primers for *GAPDH* were 5'-GTCTCCTCTGACTTCAACAGCG-3' (forward) and 5'-ACCACCCTGTTGCTGTAGCCAA-3' (reverse).

**Immunoblotting**

HeLa cells (250,000 cells/well) were seeded in 6-well plates and incubated for 24 h prior to addition of 50 ng/mL IFN-γ and compounds. After 24 h, cells were lysed using 1XLaemmli buffer (8% glycerol, 4.4%

0.5 M TRIS, pH 6.8, 3.1 mM SDS, 4.44 mM DTT, 33.2 mM bromophenol blue) and homogenized by sonication. Protein concentration was determined using the DC Protein Assay (Bio-Rad Laboratories, DE). 20 µg to 100 µg total protein was separated using 10% polyacrylamide gels under reducing/denaturing conditions in a TRIS/glycine-based system. Proteins were transferred to a PVDF membrane for 60 min at 100 V (Thermo Fisher Scientific, USA) using wet tank transfer (192 mM glycine, 25 mM TRIS, 10% methanol) in a Mini Trans-Blot® Cell (Bio-Rad Laboratories, DE). Afterwards, membranes were blocked with 5% non-fat milk in PBS with 0.1% (v/v) Tween-20 (PBS-T) or 50% Odyssey® Blocking Buffer (PBS) (LI-COR Biosciences, US) in PBS-T for 60 min at room temperature followed by overnight incubation at 4°C with the primary antibodies. For detection of IDO1, sEH and the reference protein vinculin, the primary antibodies anti-IDO1 (1:5,000 in 5% milk in PBS-T, ab211017, Abcam, UK, RRID:AB_2936946), anti-sEH (1:1,000 in 50% Odyssey® Blocking Buffer in PBS-T, A1885, ABclonal Science, US, RRID:AB_2763918) and anti-vinculin (1:10,000 in 5% milk in PBS-T or 50% Odyssey® Blocking Buffer in PBS-T, V9131, Merck KGaA, DE, RRID:AB_477629) were used. Protein bands were visualized with secondary antibodies conjugated to IRDye® Infrared Fluorescent Dyes (1:5,000 in 50% Odyssey® Blocking Buffer in PBS-T, LI-COR Biosciences, US) using the ChemiDoc MP Imaging System (Bio-Rad Laboratories, DE). IDO1 and sEH protein levels were normalized to the levels of the reference protein vinculin.

**Affinity-based chemical proteomics (pulldown)**

HeLa cells (5.54 x $10^5$ cells/flask) were seeded in T175 flasks and incubated for 72 h at 37°C and 5% $CO_2$ in humidified atmosphere. The medium was exchanged for fresh culture medium containing 50 ng/mL IFN-γ prior to incubation for 24 h. The cells were harvested by trypsinization, resuspended in ice-cold PBS and washed thrice in ice-cold PBS. The cell pellets were resuspended in 500 µL NP-40-based lysis buffer (150 mM sodium chloride, 0.4% NP-40 alternative, 50 mM TRIS-HCl, pH 8.0), incubated on ice for 30 min and ultra-centrifuged for 20 min at 100,000 x g and 4°C.

25 µL of NHS Mag Sepharose beads (cat# 28-9440-09, Cytiva, US) were equilibrated in 500 µL ice-cold 1 mM HCl, the equilibration solution was removed and 500 µL of 10 µM the free amine probes **2b** and **3b** (0.1% (v/v)) in 150 mM triethanolamine, 500 mM sodium chloride, pH 8.3 were added. The beads were coated with the probes by overhead rotation for 1 h at room temperature. Subsequently, the residual active groups were quenched by i) washing with 500 µL block buffer 1 (500 mM ethanolamine, 500 mM sodium chloride, pH 8.3), ii) washing with 500 µL block buffer 2 (100 mM sodium acetate, 500 mM sodium chloride, pH 4.0), iii) addition of 500 µL block buffer 1 and incubation for 15 min with overhead rotation, iv) washing with 500 µL block buffer 2, v) washing with 500 µL block buffer 1, vi) washing with 500 µL block buffer 2.

After the compound immobilization, the beads were washed with 500 µL NP40-based lysis buffer prior to incubation with cell lysate (500 µL of lysate with a protein concentration of 3 g/L) for 2 h at 4°C with overhead rotation. For the competition experiment, the cell lysate was pre-incubated with 10 µM unmodified epoxykynin for 1 h at room temperature prior to addition to the coated beads. Subsequently, the supernatant was removed and the beads were washed twice with 50 mM PIPES (pH 7.4), 50 mM sodium chloride, 75 mM magnesium chloride, 5 mM EGTA, 0.1% NP-40 alternative, 0.1% Triton X-100 and 0.1% Tween20 and twice with PBS for 10 min under overhead rotation at room temperature. The supernatant was removed and the samples were subjected to on-bead tryptic digestion.

Samples were resuspended in 50 µL denaturing/reducing buffer (8 M urea, 50 mM TRIS-HCl (pH 7.5) and 1 mM DTT) and incubated for 30 min at room temperature. Afterwards, samples were alkylated by addition of 5.55 µL 50 mM 2-chloroacetamide in denaturing/reducing buffer prior to incubation for 30 min at room temperature. Afterward, 1 µg Lys-C (0.5 µg/µL in ddH$_2$O, FUJIFILM Wako Pure Chemical Corporation, JP) was added and samples were incubated for 1 h at 37°C. The supernatants were transferred to new tubes. 165 µL 50 mM TRIS-HCl, pH 7.5 containing 1 µg trypsin (Roche, US) were added and incubated for 1 h at 37°C. The supernatants from both the Lys-C and trypsin digestion were combined and another 1-2 µg trypsin was added. The digestion was proceeded overnight at 37°C and the reaction was stopped by addition of 10% (v/v) TFA on the next day. The peptides were desalted by StageTip Purification.

Two layers of EmporeTM High Performance Extraction C18 Disks (3M Bioanalytical Technologies, US) were stacked on top of each other in a 200 µL pipette tip using a syringe. The filters were activated by addition of 100 µL methanol and centrifugation until all liquid has passed through the disks. The filters were washed and equilibrated once with 100 µL of 0.1% formic acid and twice with 100 µL of 80% acetonitrile, 0.1% formic acid prior to loading of the samples and incubation for 1 min. The solution was removed by centrifugation and the filters were washed with 100 µL of 0.1% formic acid. 20 µL of 80% acetonitrile, 0.1% formic acid were added and samples were incubated for 1 min prior to elution of the sample by centrifugation at 1,500 x g for 5 min. The previous step was repeated once and the supernatants were combined prior to evaporation using a vacuum concentrator at 30°C. The dried peptides were dissolved in 20 µL of 0.1% TFA and analyzed by nanoHPLC-MS/MS using an Ultimate 3000 RSLC nano-HPLC system and a Hybrid-Orbitrap mass spectrometer Q Exactive Plus or Q Exactive HF, respectively (Thermo Fisher Scientific, US, different instruments were used for the two biological replicates; technical replicates were analyzed on the same instrument). 1 µL (Q Exactive HF) or 2 µL (Q Exactive Plus) of the peptide solution was injected and enriched on a C18 PepMap 100 column (5 mm, 100 Å, 300 mm ID * 5 mm, Dionex$^{TM}$, Thermo Fisher Scientific, US,) using 0.1% TFA, at a flow rate of 30 µL/min, for 5 min and separated on a C18 PepMap 100 column (3 mm, 100 Å, 75 mm ID * 50 cm) using a linear gradient (5-30% ACN/H$_2$O + 0.1% formic acid over 90 min) with a flow rate of 300 nL/min. The nano-HPLC apparatus was coupled online with the mass spectrometer using a standard coated PicoTip® emitter (ID 20 µm, Tip-ID 10 µM, New Objective, US). Signals in the mass range of m/z 300 to 1650 were acquired at a resolution of 70,000 (Q Exactive Plus) or 60,000 (Q Exactive HF) for full scan, followed by up to ten (Q Exactive Plus) or 15 (Q Exactive HF) high-energy collision-dissociation (HCD) MS/MS scans of the most intense at least doubly charged ions at a resolution of 17,500 (Q Exactive Plus) or 15,000 (Q Exactive HF), respectively. Proteins were relatively quantified by using MaxQuant[8, 17] v.2.0.3.0, including the Andromeda search algorithm and searching in parallel the Homo sapiens reference proteome of the UniProt database and a contaminants database implemented in MaxQuant. Briefly, an MS/MS ion search was performed for enzymatic trypsin cleavage, allowing two missed cleavages. Carbamidomethylation was set as a fixed protein modification, and oxidation of methionine and acetylation of the N-terminus were set as variable modifications. The mass accuracy was set to 20 parts per million (ppm) for the first search, and to 4.5 ppm for the second search. The false discovery rates for peptide and protein identification were set to 0.01. Only proteins for which at least two peptides were quantified were chosen for further validation. Relative quantification of proteins was performed by using the label-free quantification algorithm implemented in MaxQuant.

Statistical data analysis of pulldown samples was performed separately for both biological replicates (four technical replicates each) using Perseus[9, 18] v.1.6.14.0 including proteins which were identified in at least three out of four technical replicates in at least one of the two compared conditions. Label-free quantification (LFQ) intensities were log-transformed ($log_2$); replicate samples were grouped together. Missing values were imputed using small normally distributed values, and a two-sided t-test was performed. Volcano plots were generated using the VolcaNoseR[1, 19] web app. Proteins with $log_2$-fold changes <1 or >1 and $-log_{10}(p) > 4.5$ were considered as statistically significant enriched.

**sEH-H assay**

Inhibition of the C-terminal lipid epoxide hydrolase domain of the soluble epoxide hydrolase (seH-H) was tested using the Soluble Epoxide Hydrolase Inhibitor Screening Assay Kit (cat#10011671, Cayman, US). Briefly, 2.5 µL of 50X diluted sEH protein was treated with compounds at indicated concentrations and the reaction was initiated by addition of 250 nM fluorogenic sEH-H substrate PHOME (3-phenyl-cyano(6-methoxy-2-naphthalenyl)methyl ester-2-oxiraneacetic acid)). The final volume for the assay was 100 µL in all wells. After 10 s, the fluorescence was recorded using an excitation wavelength at 330 nm and an emission wavelength at 465 nm at 25°C on the Spark® Multimode Microplate Reader (Tecan, AT) for up to 30 min in a kinetic loop with intervals of 90 s between measurements. The background was subtracted from all values and all values were normalized to the initial activity of each condition. The $IC_{50}$ value was determined by area-under-curve analysis using GraphPad Prism 9.0 (GraphPad Software, Inc, US).

**sEH-P assay**

Inhibition of the N-terminal lipid phosphatase domain of the soluble epoxide hydrolase (sEH-P) was tested using an AttoPhos-based assay as described previously.[20] 5 µL of 50X diluted sEH protein (cat#600037, Cayman, US, from the Soluble Epoxide Hydrolase Inhibitor Screening Assay Kit) was incubated with compounds at indicated concentrations in AttoPhos assay buffer (25 mM BIS-TRIS, 1 mM $MgCl_2$ x 6 $H_2O$, 0.1 mg/mL BSA, pH 7.0) for 5 min at 23°C prior to addition of 7.5 µL AttoPhos solution (166.7 µM AttoPhos reagent (cat#S1011 Promega, US) in AttoPhos assay buffer to a final concentration 25 µM. The samples were incubated for 60 min at 23°C in the dark under mild shaking. Afterwards, 25 µL of AttoPhos stop solution (100 mM NaOH in AttoPhos buffer) was added and the fluorescence was recorded using an excitation wavelength at 435 nm and an emission wavelength at 555 nm on the Spark® Multimode Microplate Reader (Tecan, AT) for up to 30 min in a kinetic loop with intervals of 90 s between measurements. The final volume for the assay was 75 µL in all wells. The analysis was performed using the data from 15 min after the stop solution was added, i.e. when the fluorescence signal became stable. Therefore, all values were subtracted by the background and normalized to the DMSO control.

**nanoDSF**

Purified sEH protein (cat#600037, Cayman, US, from the Soluble Epoxide Hydrolase Inhibitor Screening Assay Kit) was diluted 18X in AttoPhos assay buffer and incubated with compounds at indicated concentrations for 10 min at 22°C. The thermal protein stability from 20°C to 90°C (1°C/min) was measured by means of the intrinsic tryptophan/tyrosine fluorescence using high sensitivity capillaries in the Prometheus™ NT.48 (NanoTemper® Technologies, DE). Melting scans, first derivatives of melting scans and melting temperatures were analyzed using the PR.ThermControl software (NanoTemper® Technologies, DE).

**In-Cell CETSA**

Jurkat cells (10 x $10^6$ cells/flask) were seeded in two T25 tissue culture flasks and treated with 10 µM compound or DMSO for 20 min at 37°C. Cells were harvested and washed thrice in ice-cold PBS. Compound- and DMSO-treated samples were distributed equally into ten tubes each and subjected to heating at different temperatures ranging from 37°C to 72.1°C (37°C, 51.5°C, 55.2°C, 58.1°C, 61.9°C, 63.8°C, 65.7°C, 67.6°C, 69.2°C, 72.1°C) in the Mastercycler X50s (Eppendorf SE, DE). Afterwards, cOmplete™ EDTA-free Protease Inhibitor Cocktail (Roche, CH) and NP-40 alternative, sodium chloride and TRIS-HCl, pH 8.0 (final concentrations of 0.4% (v/v), 150 mM and 50 mM, respectively) were added and cells were lysed by three consecutive freeze/thaw cycles. Soluble fractions were separated from denatured proteins by centrifugation at 20,000 x g and 4°C for 25 min. Supernatants were transferred to new tubes and subjected to immunoblot analysis.

**sEH NanoBRET**

Cellular target engagement was studied with the NanoBRET technology using a full-length sEH-NanoLuc construct and the fluorescent sEH-H ligand **4** as published previously.[21] HEK293T cells (5 x $10^5$ cells/well) were seeded in 6-well plates containing 2 mL growth medium and incubated for 24 h. Afterwards, the medium was removed and the cells were washed with PBS prior to transfection with sEH-aa1-aa555_C-NanoLuc_pF32Kp[21] plasmid (2.5 µg/well) using 3.75 µL Lipofectamine™ 3000 and 5 µL P3000™ reagent (Invitrogen, US) in Opti-MEM according to the manufacturer's protocol. The cells were incubated for 4 h and subsequently, the transfection mix was replaced by growth medium and the cells were incubated for additional 20 h. On the next day, the cells were washed thrice with PBS and harvested in Opti-MEM. The cell titer was determined and the concentration was adjusted to 5 x $10^4$ cells/mL.

4.25 x $10^4$ cells, 60 nM of the fluorescent sEH-H tracer **4** and epoxykynin at indicated concentrations were mixed in a total volume of 20 µL in a white 384-well plate. The plate was sealed with an AeraSeal™ film (Sigma-Aldrich, US) and incubated at 37°C and 5% $CO_2$ for 5 h. 9 µL NanoGlo substrate and 3 µL extracellular NanoLuc inhibitor (Intracellular TE Nano-Glo® Kit, Promega, US) were diluted in Opti-MEM with a total volume of 1.5 mL to yield the NanoGlo mixture. Afterwards, 10 µL of the NanoGlo mixture was added to each well prior to short centrifugation and measurement of the BRET signal. Firstly, the donor luminescence was monitored at 458 nm with a bandwidth of 25 nm.for 500 ms on the Spark® Multimode Microplate Reader (Tecan, AT). Subsequently, the acceptor fluorescence was measured at 578 nm with a bandwidth of 25 nm.for 500 ms. The BRET ratio was determined by multiplying the acceptor signal by 1000 and dividing by the donor signal. After plotting of the BRET ratio against the logarithmic compound concentration, data analysis was performed using a non-linear regression curve fit in GraphPad Prism 9.0 (GraphPad Software, Inc, US) to generate dose-response curves and obtain $IC_{50}$ values.

**Co-crystallization of epoxykynin and human sEH-H**

The C-terminal hydrolase domain of human sEH (sEH-H, aa222-aa555) was cloned and expressed in *E. coli* as described previously.[3, 22, 23] Briefly, sEH-H protein was expressed in *E. coli* BL21(DE3) using ZYP5052 autoinduction medium[24] at 16°C for 36 h. sEH-H was purified by nickel affinity chromatography in 50 mM TRIS-HCl, pH 8.0, 500 mM NaCl, 70 mM imidazole-HCl and eluted in 50 mM TRIS-HCl, pH 8.0, 500 mM NaCl, 400 mM imidazole-HCl, followed by size-exclusion chromatography using a HiLoad 16/60 Superdex 200 column (GE Healthcare, US) in 50 mM NaCl, 50 mM sodium phosphate, 10% (v/v) glycerol (98%), 2 mM DTT, pH 7.4. The pure protein was concentrated with an Amicon Ultra-15 (10 kDa cutoff).

440 µM sEH-H (16.7 mg/mL) was mixed with 1 mM epoxykynin for 1 h on ice. Initial crystallization trials were performed using commercially available screens. A drop volume of 200 nL was equilibrated against 20 µL of reservoir solution in 96-well sitting drop plates (SWISSCI, UK). The crystallization plates were incubated at 20°C. The crystals were obtained in 25% PEG Smear High, 0.1 M PIPES, pH 7.0, 0.1 M magnesium formate, 0.1 M rubidium chloride.

A single crystal was picked from the crystallization plate, treated with 25% ethylene glycol in reservoir solution as the cryoprotectant, and frozen in liquid nitrogen for each complex. Synchrotron X-ray diffraction data was acquired from the X06SA beamline at the Swiss Light Source at the Paul Scherrer Institute, CH. The temperature was maintained at 100 K during the data collection. A total of 900 images were collected, and the data were processed using XDS.[10] The intensities were scaled and converted into structure factors using AIMLESS[25, 26] of the CCP4 suite.[5] The structure was solved by molecular replacement using the MOLREP[27] program of the CCP4 suite with the published sEH-H structure from pdb 7P4K[28] as the search template. The model was built manually using COOT[6] and the structure was refined by REFMAC of the CCP4 suite. The model was validated using the wwPDB Validation System (https://validate-rcsb-2.wwpdb.org/) prior to deposition. The data collection, processing, and refinement statistics are shown in Table S6. The coordinate was deposited in the Protein Data Bank with the pdb ID 8QZD.

**Knockdown experiments**

HeLa cells (150,000 cells/well) were seeded in 6-well plates in medium without phenol red and incubated for 24 h. On the next day, cells were treated with 50 ng/mL IFN-γ and 164.15 µM Trp and transfected with siRNA. The siRNAs (ON-TARGETplus Human EPHX2 siRNA SMARTPool (cat# L-010006-00-0005, Horizon Discovery, UK) and ON-TARGETplus Non-targeting Control siRNAs (cat# D-001810-01-05, Horizon Discovery, UK)) were diluted to 2 µM in Opti-MEM in a total volume of 50 µL (final siRNA concentration: 30 nM to 100 nM). The DharmaFECT 1 reagent was diluted in Opti-MEM in a total volume of 50 µL (siRNA to transfection reagent ratio of 1 to 15). Both siRNA and transfection reagent were incubated for 5 min at room temperature prior to addition of the DharmaFECT solution to the siRNA solution. The transfection mixture was incubated for another 20 min at room temperature to allow formation of the siRNA:lipid complex. Subsequently, 1.9 mL of culture medium was added to the transfection mixture and the culture medium on the cells was replaced by the transfection mixture. The cells were incubated for up to 96 h. To quantify Kyn levels, 100 µL of the supernatant was transferred to a transparent 96-well plate and Kyn levels were determined with *p*-DMAB. The remaining supernatant was aspirated and RNA was extracted for qPCR analysis as described above or cells were lysed for immunoblotting. Therefore, the cells were resuspended in 500 µL NP-40-based lysis buffer (150 mM sodium chloride, 0.4% NP-40 alternative, 50 mM TRIS-HCl, pH 8.0), incubated on ice for 30 min and centrifuged for 25 min at 20,000 x g and 4°C. The supernatant was subjected to immunoblotting.

**Overexpression experiments**

HeLa cells (50,000 to 100,000 cells/well) were seeded in a 12-well plate in medium without phenol red and incubated for 24-48 h. Subsequently, cells were treated with 50 ng/mL IFN-γ and 164.15 µM Trp and transfected with 55 to 438 ng/well pCMV3-EPHX2 (Sino Biological, CN) using Lipofectamine™ 3000 (2 µL transfection reagent per µg plasmid DNA, Invitrogen, US) prior to incubation for 24-48 h. To quantify Kyn levels, 100 µL of the supernatant was transferred to a transparent 96-well plate and Kyn levels were determined with *p*-DMAB. The remaining supernatant was aspirated and the cells were resuspended in 500 µL NP-40-based lysis buffer (150 mM sodium chloride, 0.4% NP-40 alternative, 50 mM TRIS-HCl, pH 8.0), incubated on ice for 30 min and centrifuged for 25 min at 20,000 x g and 4°C. The supernatant was subjected to immunoblotting.

### 4. NMR Spectra

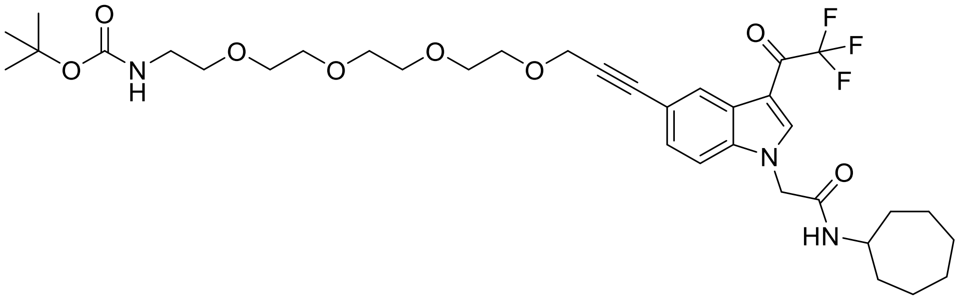

**(2a)**

1H NMR

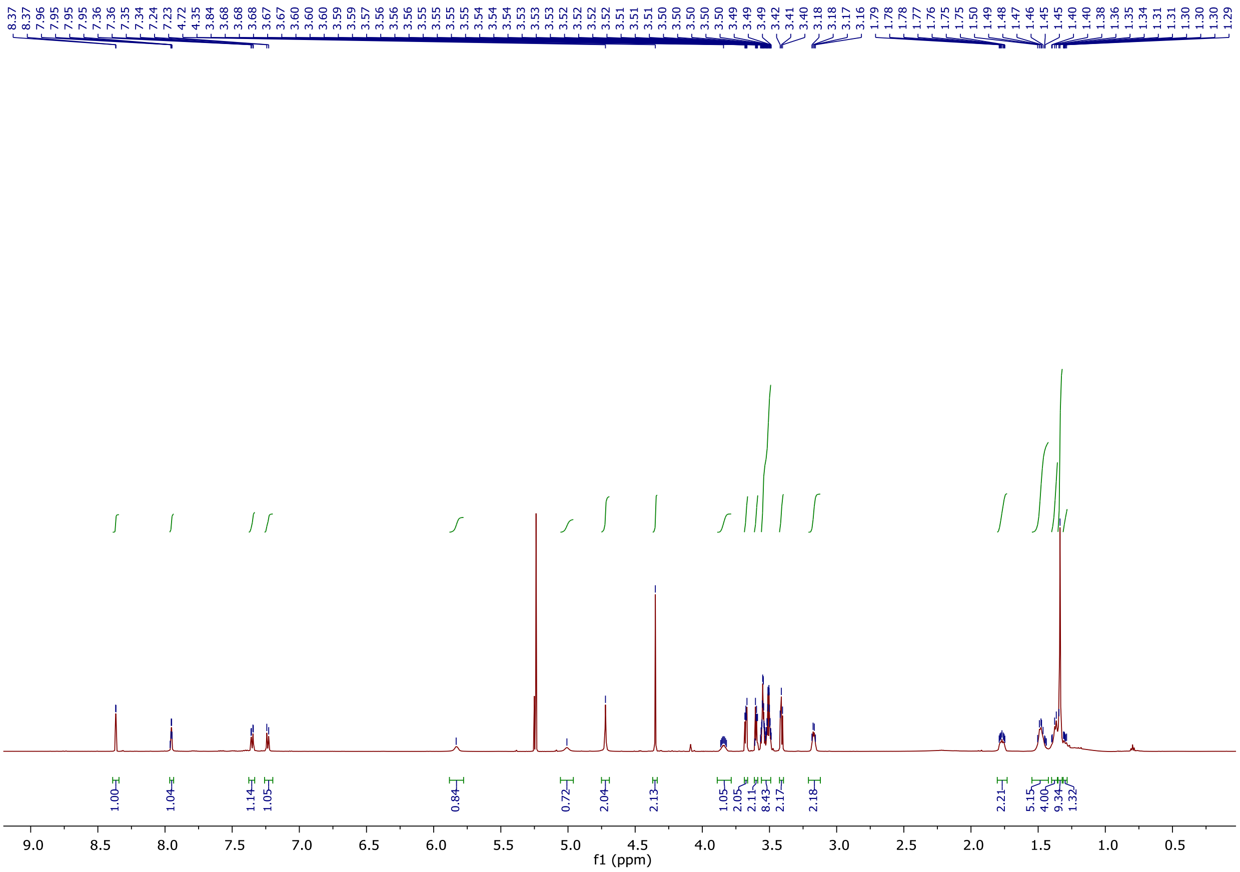

13C NMR

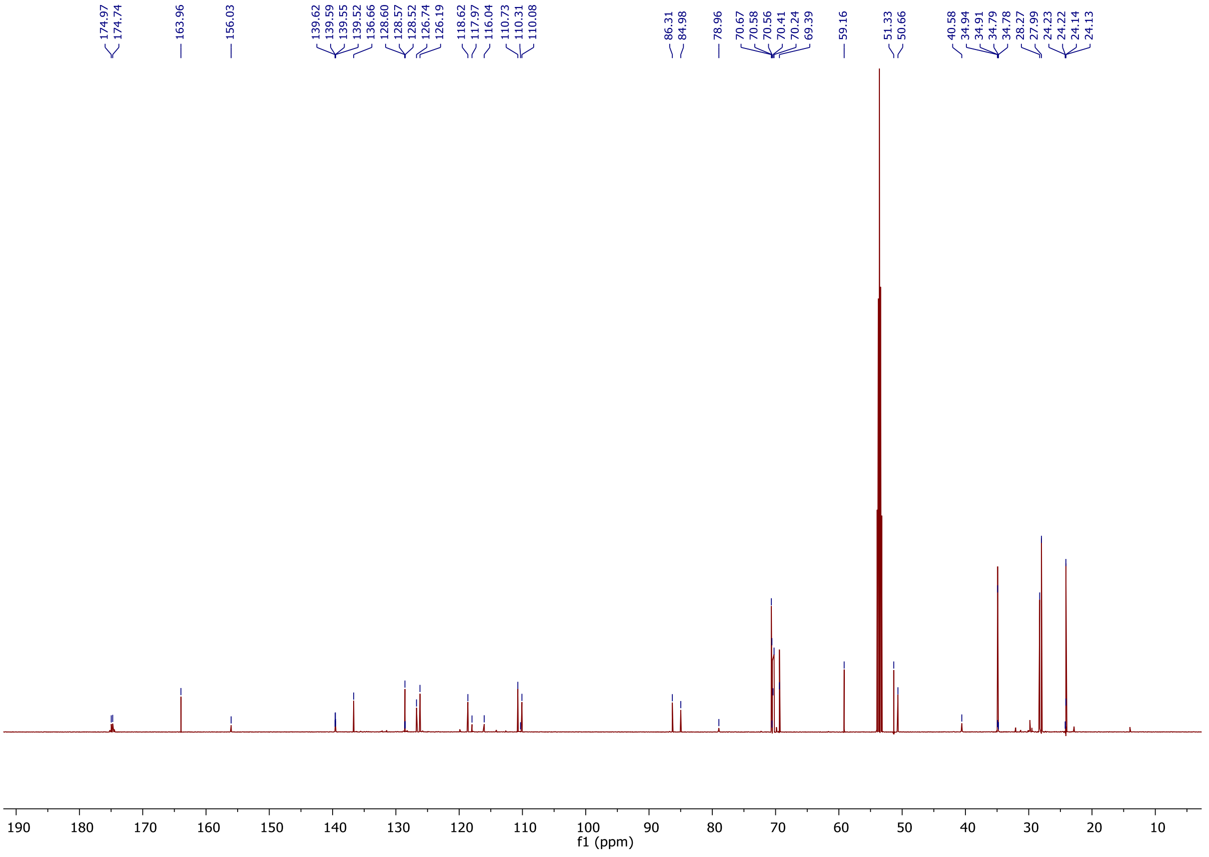

19 F NMR

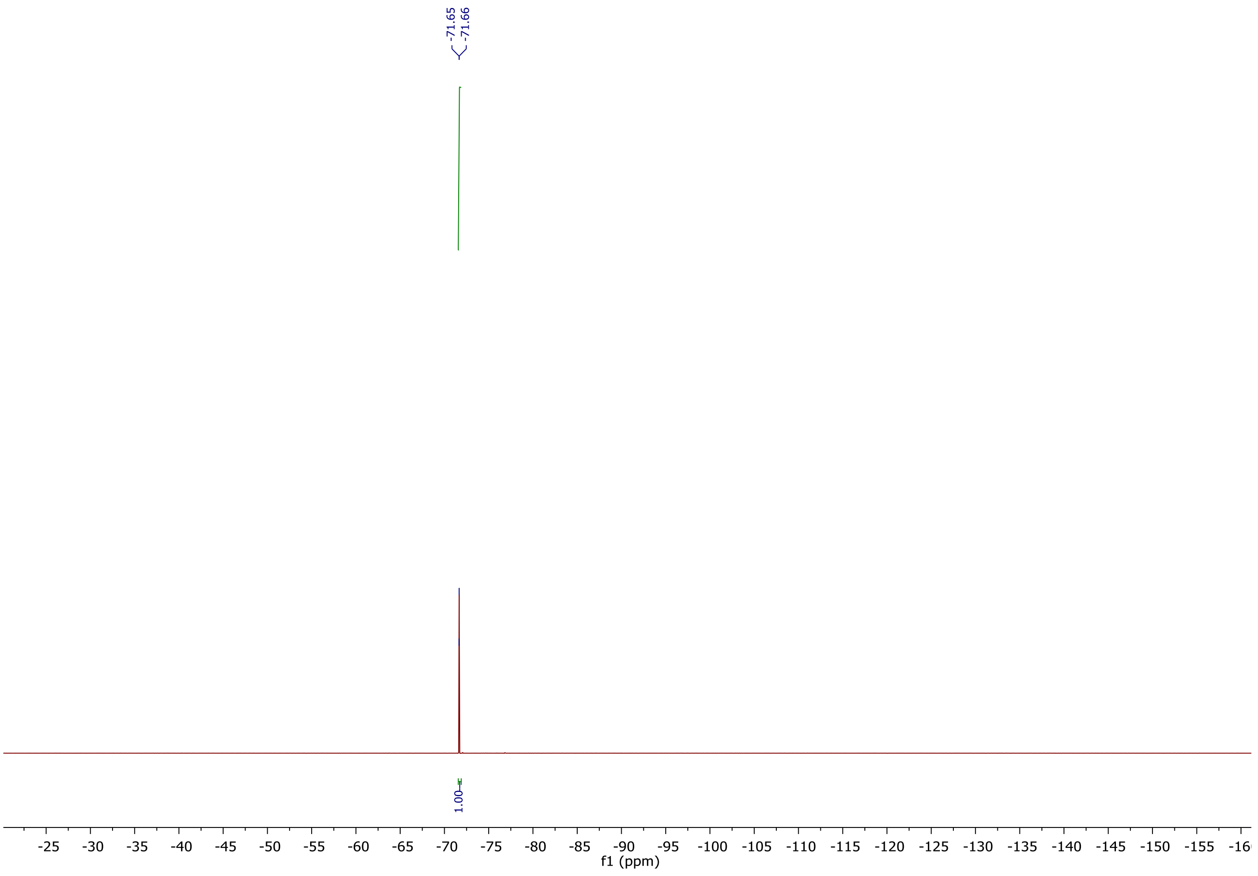

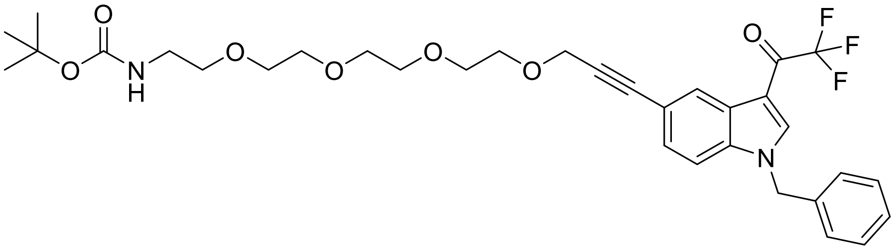
O
O
N
H
O
O
O
O
O
F
F
F
N

(3a)

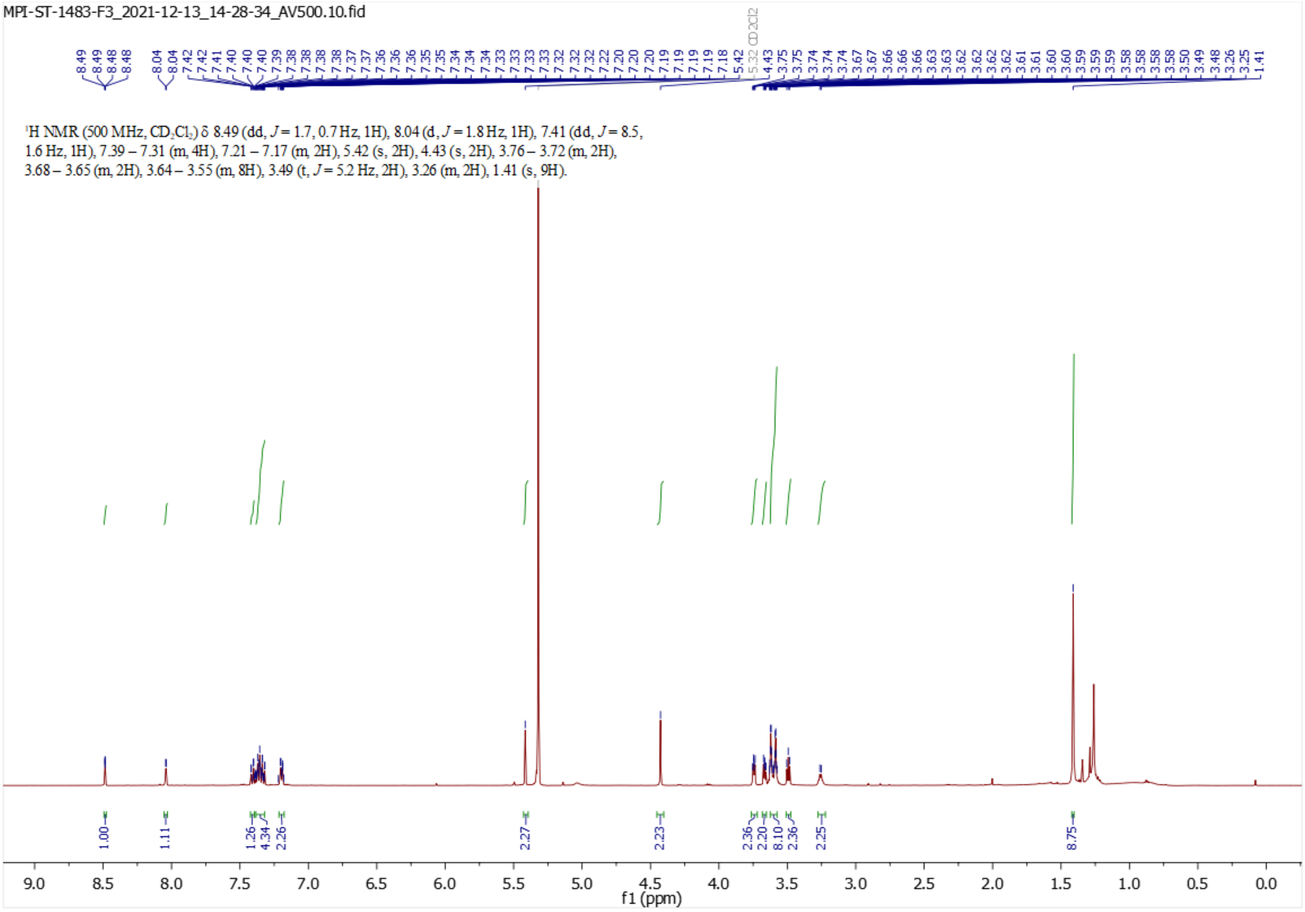
MPI-ST-1483-F3_2021-12-13_14-28-34_AV500.10.fid
CD2Cl2
1H NMR (500 MHz, CD2Cl2) δ 8.49 (dd, J = 1.7, 0.7 Hz, 1H), 8.04 (d, J = 1.8 Hz, 1H), 7.41 (dd, J = 8.5, 1.6 Hz, 1H), 7.39 – 7.31 (m, 4H), 7.21 – 7.17 (m, 2H), 5.42 (s, 2H), 4.43 (s, 2H), 3.76 – 3.72 (m, 2H), 3.68 – 3.65 (m, 2H), 3.64 – 3.55 (m, 8H), 3.49 (t, J = 5.2 Hz, 2H), 3.26 (m, 2H), 1.41 (s, 9H).
1.00
1.11
1.26
4.34
2.26
2.27
2.23
2.36
2.20
8.10
2.36
2.25
8.75
9.0
8.5
8.0
7.5
7.0
6.5
6.0
5.5
5.0
4.5
4.0
3.5
3.0
2.5
2.0
1.5
1.0
0.5
0.0
f1 (ppm)

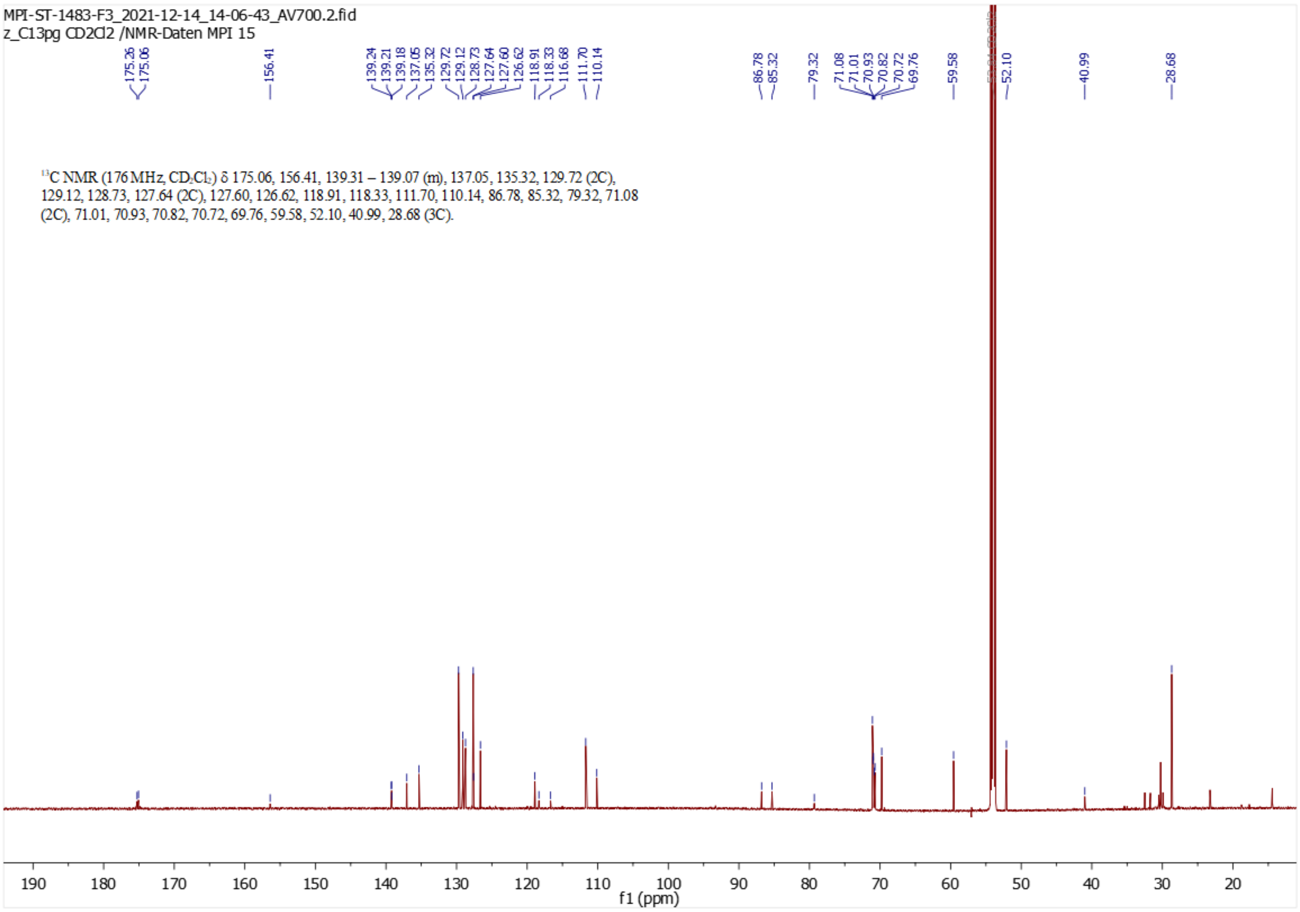
MPI-ST-1483-F3_2021-12-14_14-06-43_AV700.2.fid
z_C13pg CD2Cl2 /NMR-Daten MPI 15
13C NMR (176 MHz, CD2Cl2) δ 175.06, 156.41, 139.31 – 139.07 (m), 137.05, 135.32, 129.72 (2C), 129.12, 128.73, 127.64 (2C), 127.60, 126.62, 118.91, 118.33, 111.70, 110.14, 86.78, 85.32, 79.32, 71.08 (2C), 71.01, 70.93, 70.82, 70.72, 69.76, 59.58, 52.10, 40.99, 28.68 (3C).
f1 (ppm)

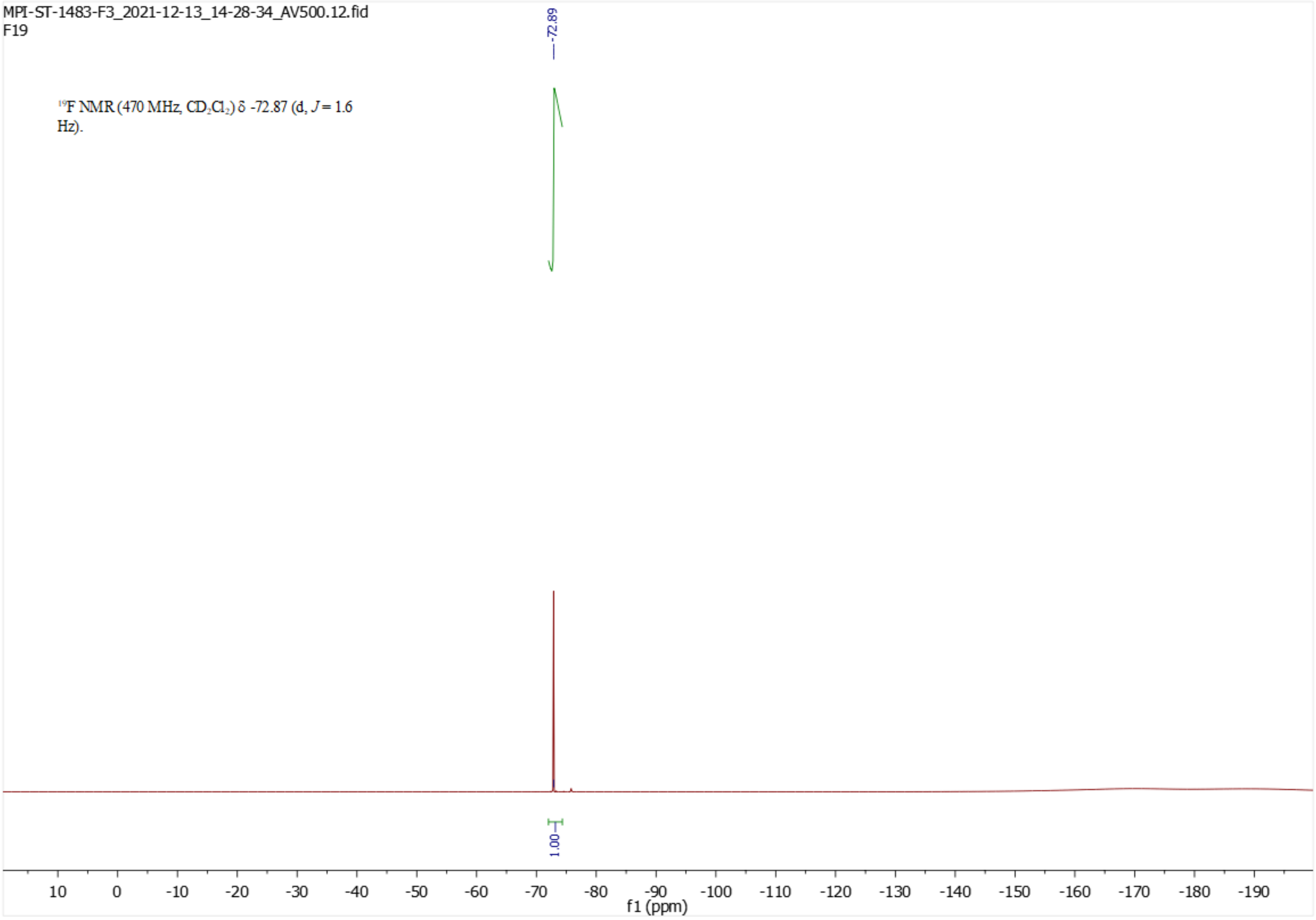
MPI-ST-1483-F3_2021-12-13_14-28-34_AV500.12.fid
F19
-72.89
19F NMR (470 MHz, CD2Cl2) δ -72.87 (d, J = 1.6 Hz).
1.00
f1 (ppm)
